# A single freshwater invasion gave rise to the world’s dominant freshwater fish group

**DOI:** 10.64898/2026.10.26.761952

**Authors:** Delson H. Hays, Matthew A. Burnett, Aintzane Santaquiteria, Dahiana Arcila, Alexander Burnett, Olga Otero, Diogo Mayrinck, Ricardo Betancur-R

## Abstract

The ultraspeciose clade Otophysa — catfishes, carps, minnows, tetras, and their kin — accounts for most vertebrate diversity in modern freshwater ecosystems. However, a controversial assemblage of largely-fragmentary marine fossils has led some authors, most recently Liu et al., to advocate parallel freshwater invasions for this otherwise inland lineage. Here, we test this hypothesis under multiple sources of uncertainty. We provide the first phylogenetic analysis of the recently-named marine fossil genus †*Sooinichthys*, resolving it as sister to all modern otophysans. We also find that †*Santanichthys*, long treated as an early marine otophysan, lacks a Weberian apparatus and plausibly belongs outside Ostariophysi entirely. When increasing the taxon and marker sampling of Liu et al.’s genomic dataset, standardizing their ecological coding, accounting for alternative interpretations of important morphological characters, and applying a suite of biogeographic models, we find a marine origin in only 6 of 132 analyses, and only when enforcing fossil placements inconsistent with our own phylogenetic results. We contend that the parallel freshwater invasion hypothesis of otophysan diversification is an artifact of sparse taxon sampling, taphonomic biases, model selection errors, and fossil overinterpretation. The result is a cautionary case for inferences about the cradle of any ancient freshwater clade.

## Introduction

One in every seven vertebrate species belongs to Otophysa, a freshwater fish clade comprising catfishes, tetras, carps, South American knifefishes, lutefishes, and their relatives. This taxonomic superdiversity (over 12,000 species divided amongst five orders and 95 families, 1–2) is coupled with unparalleled ecological breadth and a worldwide distribution. Otophysan fishes natively occur in every major freshwater system on Earth, including stagnant tropical swamps, fast-flowing temperate rivers, frigid polar streams, and even subterranean habitats throughout Africa, Eurasia, and the Americas (3–4). Yet despite this ecological breadth, more than 99% of extant otophysan species are limited to freshwater ecosystems; true marine species occur in only two catfish families (Ariidae and Plotosidae), which colonized the sea in parallel following the end-Cretaceous mass extinction (4–5).

Biogeographic analyses with extensive fossil sampling indicate that all living actinopterygians are ultimately descended from marine ancestors, and therefore, that freshwater endemism is a derived condition in fishes (6). However, the freshwater invasion preceding otophysan diversification has long been difficult to pinpoint phylogenetically. Three orders collectively comprising the closest relatives to Otophysa — Gonorynchiformes, Alepocephaliformes, and Clupeiformes — are a heterogeneous assemblage of freshwater, euryhaline, and oceanic taxa, with some members restricted to shallow tropical floodplains and others to the marine aphotic zone, while still others are diadromous and/or euryhaline (2–3). Despite this rootward uncertainty, the modern-day uniformity of freshwater life histories within Otophysa has long indicated that the otophysan radiation traces to a single ancient invasion of continental waters by a marine ancestor, with subsequent diversification occurring in the rivers and lakes of Pangaea and/or its daughter continents (7–9).

However, several studies (10–11) have leveraged paleontological data to challenge this assumption, most recently Liu et al. (12–13). Early fossil otophysans, though rarely described and often poorly preserved, are characterized by a surprising overrepresentation of marine taxa (14). While this incongruence can be partially explained by known taphonomic biases disfavoring the recovery of freshwater fauna (15–16), the prevalence of marine forms indicates that early otophysans unequivocally lacked the freshwater homogeneity seen today, a pattern observed in several ancient fish clades (17–19). However, unlike in other clades, the affinities of virtually all early ostariophysan fossils remain contentious. Many such taxa, such as the marine genera †*Santanichthys*, †*Chanoides*, and †*Nardonoides*, are poorly-characterized, while others are only recently described (i.e., the marine †*Sooinichthys varii*, potentially a stem otophysan but without phylogenetic analysis to substantiate its affinities (20), and the freshwater †*Acronichthys maccagnoi*, resolved inside the otophysan crown by Liu et al. (12)). The placements of these fossil lineages constitute the primary line of evidence in characterizing the habitat transition(s) that gave rise to the bulk of modern freshwater vertebrate biodiversity.

Here, we contend that previous works (10–13) recovering multiple otophysan freshwater invasions are affected by four shortcomings: (i) the phylogenetic irresolution of key fossil taxa; (ii) the exclusion of major extant lineages; (iii) the application of inappropriate habitat coding schemes; and (iv) the misspecification of biogeographic models during ancestral habitat reconstruction. We replicate the most recent analysis of historical otophysan habitat use (12) with a more conservative morphological coding for four fossil taxa, and with alternative coding schemes to account for uncertainty. We further increase taxon sampling from 46 to 88 species (gaining several extant families and one major fossil lineage), estimate and calibrate a novel suite of total-evidence phylogenies with a 7x-increased genomic dataset, and perform ancestral habitat reconstructions using a simplified character coding, under different taxon samplings, topologies, and biogeographic models. Our results consistently indicate that the modern-day abundance of otophysan fishes can be traced to a single freshwater invasion no later than the early Jurassic.

### Anatomical diagnoses of fossil otophysans

Otophysans are diagnosed by the Weberian apparatus, a sound-amplifying organ formed by a chain of small bony elements (21–27). Four pairs of ossicles (the claustra, scaphia, intercalaria, and tripodes) transmit auditory information posteriorly, from the inner ear to the swim bladder, which functions as a resonance chamber. Additional modifications to the adjacent cranial bones, vertebrae, and pleural ribs accommodate these ossicles. Through improving nonvisual sensory capacity in turbid estuarine and/or riverine waters (22, 28), the evolution of this apparatus may have partially facilitated the otophysan freshwater invasion(s) (12). With the exception of a small number of electroreceptive clades (e.g., gymnotiforms and some siluroid catfishes), which appear to have secondarily lost their claustra (21), these eight bones and their associated skeletal modifications are found in all otophysan fishes, making them highly valuable in defining the phylogenetic affinities of fossil taxa.

However, the small size of the Weberian ossicles, together with the imperfect preservation of many early otophysan fossils make it difficult to construct reliable morphological matrices. The recently-named North American genus †*Acronichthys*, treated as Otophysa *incertae sedis* and putatively resolved as a stem member of the otherwise-African Cithariniformes, is described from three specimens that suffer heavily from this conundrum. The coding of †*Acronichthys* by Liu et al. (12) has been called into question (29), and after conducting a revision of available materials for this taxon, we find that fewer than 50% of characters are reliably codable given the available fossil material. These characters include fewer than 40% of those corresponding to the Weberian anatomy (Supplementary Figure SX01; Supplementary Table TX1), though we do find acceptable evidence for a primitive Weberian apparatus. A similar problem is also noted in †*Nardonoides*, for which we find that only about 80% of characters employed by Liu et al. (12) are codable.

Two other fossil genera currently ascribed to Otophysa are better preserved: †*Chanoides* and †*Santanichthys* (8, 13, 25, 29–30), which jointly drive the recovery of a marine otophysan origin in the analyses of Liu et al. (12). However, our review of these fossils indicated that both taxa are miscoded in the matrix of Liu et al. (12). Taphonomy has caused disparate bony elements to artefactually resemble the Weberian ossicles in these forms, thereby confounding phylogenetic and biogeographic analyses. We contend that — despite exhibiting a supraneural pattern akin to that of modern otophysans — †*Santanichthys* entirely lacks a Weberian apparatus, and that the structures previously interpreted as Weberian ossicles are a displaced scale, rib, and fragments of cranial bone (see Supplementary Materials, section S7). This precludes it from placement within crown Otophysa, a result that is further supported by our interpretation of several traits in the caudal fin, jaw, and first four vertebrae (Supplementary Table TX6), as well as by our phylogenetic analyses. †*Chanoides* and †*Nardonoides* are considered to have primitive and poorly-characterized Weberian apparati, if any such apparati at all, with important skeletal articulations missing and some ossicles only dubiously observable (Supplementary Figure SX01; Supplementary Table TX1). They are likely non-otophysans.

### Phylogenetic inference and morphological matrix sensitivity

Within Otophysa, several relationships have been reliably resolved using both morphological and molecular datasets (14, 23, 26–27, 31–36). Most notably, otophysans are stably divided into two reciprocally monophyletic superorders: the Cypriniphysae (order Cypriniformes and their prehistoric kin) and the Characiphysae (orders Characiformes, Cithariniformes, Gymnotiformes, and Siluriformes) (1, 36). Otophysans in turn form a clade (Ostariophysi) with the anotophysans, which are subdivided into the marine gonorynchoids, freshwater knerioids, and euryhaline chanoids (36).

We evaluated these relationships by combining our revised morphological matrix modified from Liu et al. (12; 63 characters) with their 353-locus UCE alignment (∼95 kb) and a novel alignment of 1,133 single-copy exons following Hughes et al. (37, ∼617 kb; Supplementary Table TX2). In addition to substantially increasing the size of our genomic dataset, this allowed us to incorporate 42 additional taxa, including key lineages (Denticipitoidei, Diplomystoidei, Cobitoidei, and Alepocephaliformes) missing from Liu et al. (12)’s analyses. We used this total-evidence matrix to estimate phylogenies with RAxML-NG (38; Fig. 1), which were then calibrated under the Fossilized Birth Death (FBD) model and a Bayesian framework (39–40). The resultant topologies broadly concur with previous literature in the relationships of major extant lineages, including resolving the reciprocal monophylies of Characiphysae and Cypriniphysae, and Otophysa and Anotophysa.

**Fig 1.**
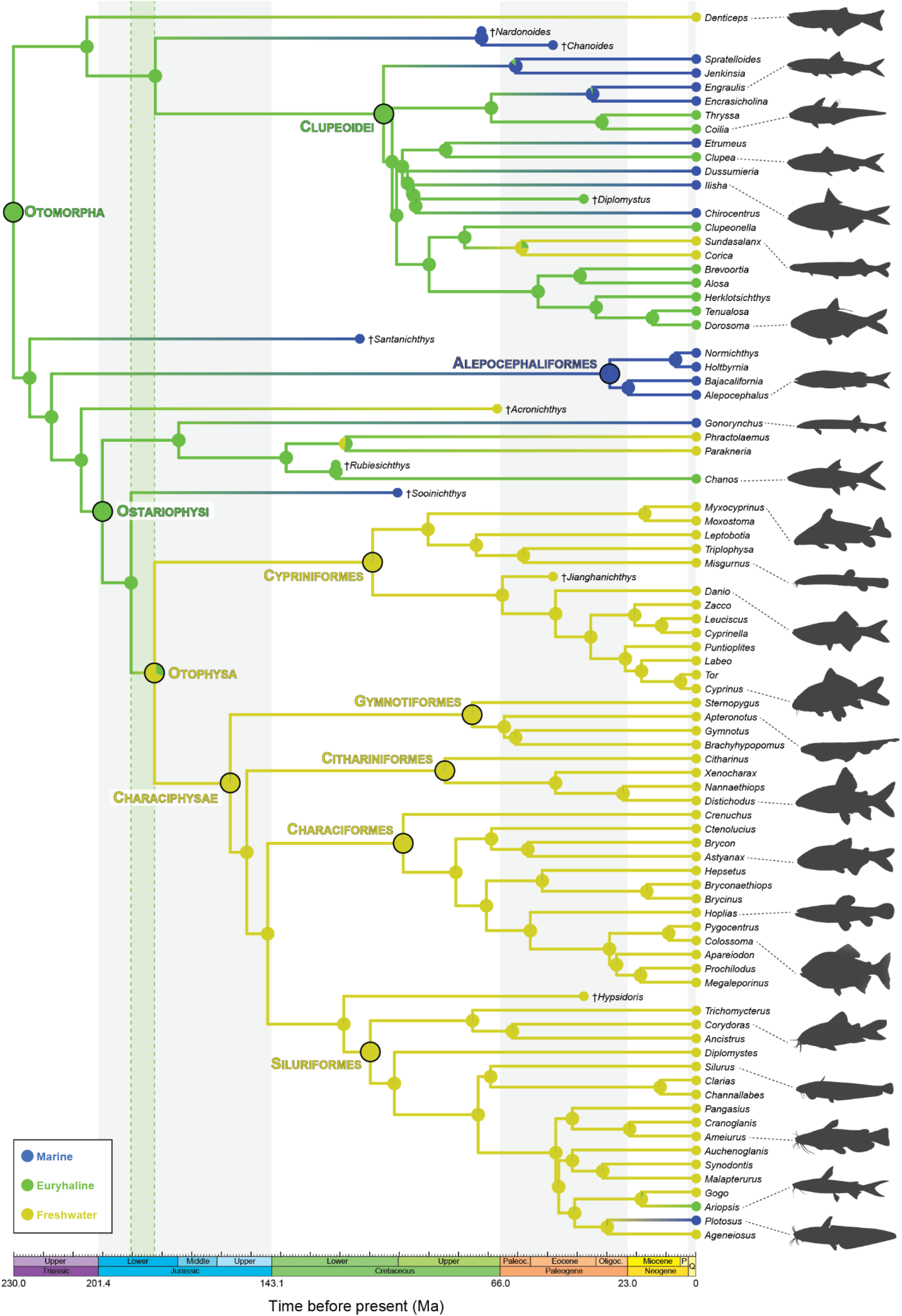
Ancestral habitat reconstruction of otocephalan fishes using a total-evidence (morphology, exons, and UCEs) phylogenetic analysis, calibrated under the fossilized birth death (FBD) model. The best-fitting BioGeoBEARS model (BAYAREA+J) fitted to the best-supported RAxML topology recovers euryhalinity throughout ancient Otomorpha, persisting into the crowns of Gonorynchiformes and Clupeoidei. Otophysans became restricted to freshwater habitats between 182 and 191 million years ago (highlighted with vertical green bar), during the breakup of Pangaea. Branches colored by the most-likely state selected by BioGeoBEARS, with transitions represented by color gradients, and individual node probabilities represented with pie charts. Major taxa labelled in bold. Silhouettes of representative taxa shown at right.

We resolve new putative relationships for five otocephalan fossil genera considered *incertae sedis* at the order level. Most pivotal of these genera is †*Santanichthys*, which has long been included in the crown of Otophysa (30), but is herein removed from Ostariophysi entirely. Across the five best-supported topologies recovered by our phylogenetic analyses, †*Santanichthys* is never resolved within the crown of Otophysa (see Figs. SX02–23 for exact placements); most frequently and in the best-supported topology, it forms an apparent trichotomy with Clupeomorpha and Alepocephaliformes–Ostariophysi. We provisionally consider it the sister clade to Alepocephaliformes–Ostariophysi, based on a suite of skeletal characters that unite these taxa to the exclusion of Clupeiformes (Supplementary Table TX6). †*Chanoides* and †*Nardonoides* are recovered in Clupeomorpha, stem Otophysa, or Alepocephaliformes, depending upon analysis; †*Acronichthys* is recovered in the stems of Ostariophysi, Otophysa, or Cithariniformes (as by Liu et al. (12)); and †*Sooinichthys* in stem Otophysa.

To isolate the impact of our morphological reinterpretation, we also estimated morphology-only phylogenies both with and without a constraint to the topology proposed by Liu et al. (12) across three analytical frameworks (Fig. 2). Maximum parsimony (MP) analyses performed in TNT (44–45) strongly favored the unconstrained topology, resolving all four marine fossil genera (†*Chanoides*, †*Nardonoides*, †*Santanichthys*, and †*Sooinichthys*) outside the otophysan crown across all equal and implied weighting schemes, though their precise placement varied slightly with the scaling of the concavity constant (*k*; Supplementary Table TX7). Maximum likelihood (ML) analyses largely mirrored these results, decisively placing †*Santanichthys* outside Ostariophysi and the remaining marine fossils as stem otophysans (Fig. 2; Supplementary Table TX8), whereas Bayesian Inference (BI) recovered all the marine fossils as stem otophysans (Fig. 2; Supplementary Table TX9). Both tests supported our unconstrained topology when using the revised matrix. The AU test rejected the constrained topologies but failed to reject the unconstrained topology (pAU = 0.966), while the Bayes factor strongly favored the unconstrained topology (BF = 33.572; Supplementary Table TX9). Using the matrix of Liu et al., the AU test also failed to reject our unconstrained topology (pAU = 0.077).

**Fig 2.**
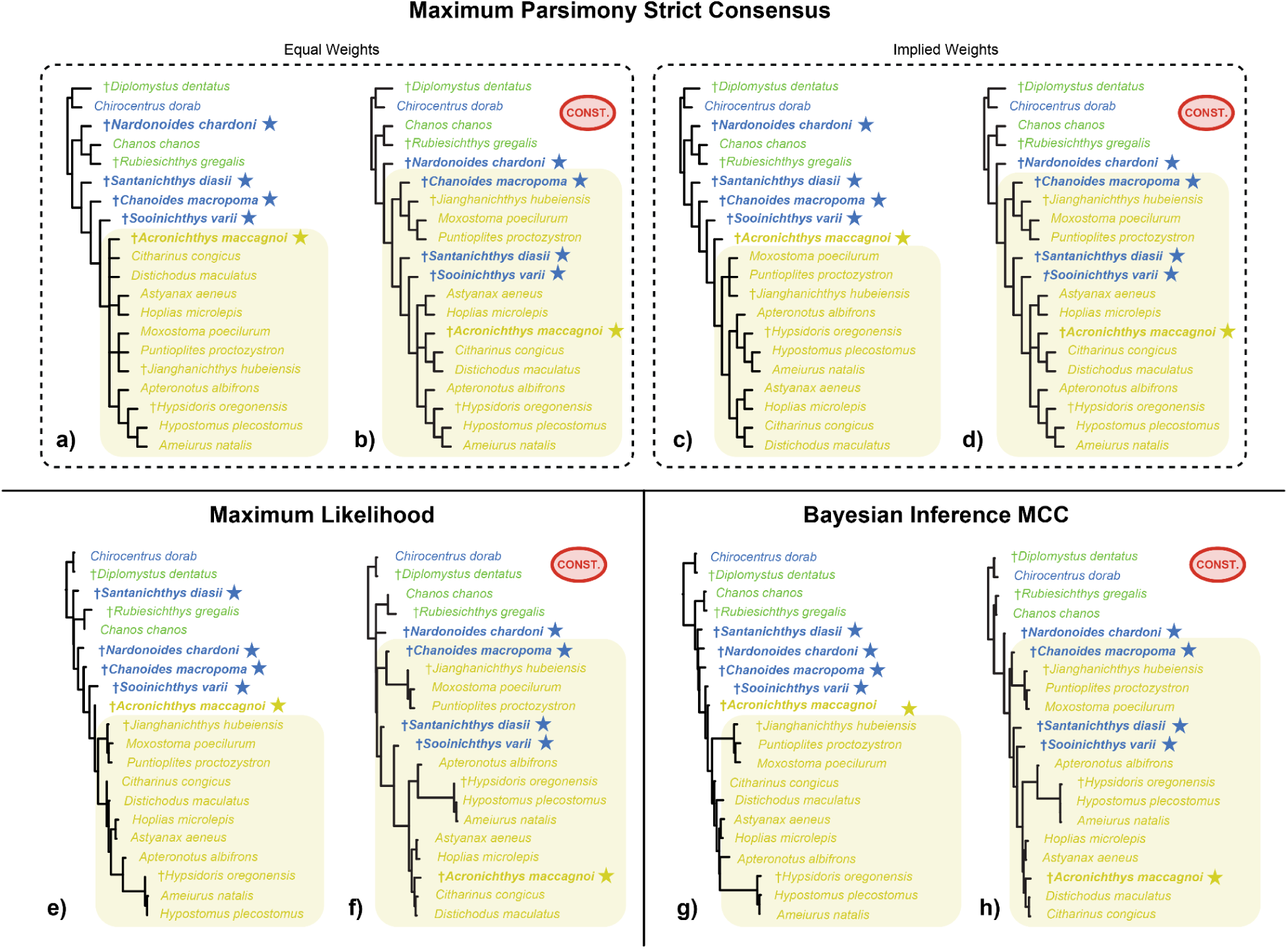
Morphological trees estimated under different optimality criteria (Maximum Parsimony, Maximum Likelihood, and Bayesian inference) concur in resolving no marine fossils within the otophysan crown. Conversely, analyses constrained to the topology of Liu et al. (12) (indicated with red CONST. ellipses) resolve †*Sooinichthys* alongside †*Chanoides* and †*Santanichthys* in the otophysan crown. Taxa are color coded by habitat occupancy (blue = marine, green = euryhaline, yellow = freshwater), and starred taxa are recoded or newly coded in the present study. Yellow highlight corresponds to crown Otophysa. **a–d)** MP phylogenies estimated in TNT (45) using either equal weights (**a–b**) or implied weights (**c–d**); the topology shared by the most values of the concavity constant (*k*) is presented (Supplementary Materials, section S1). **e–f)** ML phylogenies estimated using IQ-Tree (47). **g–h)** Maximum-clade consensus (MCC) trees of Bayesian analyses performed using MrBayes (48).

To further investigate the impact of our morphological revision, we conducted sensitivity analyses using three “hybrid” matrices. Each hybrid matrix was built by substituting the coding of the original matrix (12) with our alternative coding for one of three key anatomical regions: Weberian apparatus, caudal fin, and jaws(Fig. SX01). Morphological trees for each hybrid matrix were estimated without topological constraint, using the same set of MP, ML, and BI frameworks (see Supplementary Materials, section S1). All trees using revised coding for the caudal region and Weberian apparatus resolved all four marine genera outside of the otophysan crown (Supplementary Figure SX24), though these placements were, once again, dependent on the optimality criterion. Conversely, our revised coding for the jaw region, when paired with the original coding of the Weberian apparatus and caudal fin, resolves †*Santanichthys* and †*Sooinichthys* as crown Characiphysae. This result suggests that the spurious placement of †*Santanichthys* within Otophysa is affected by miscoding of traits in both the caudal and the Weberian anatomy.

### A freshwater (Pangaean) origin of Otophysa

Though the parallel freshwater invasion hypothesis has been advanced previously (10–11), Liu et al. (12) offer the first explicit biogeographic analysis to accompany this claim, proposing that two parallel freshwater invasions occurred during the mid-to-early Cretaceous — one in Gondwana (Characiphysae) and the other in Asia (Cypriniphysae) (12). We contend that such biogeographic questions cannot be addressed with their dataset. The sparse taxon sampling of that study, which only captures about one quarter (26 / 95) of living otophysan families and less than 0.5% of living species, precludes such an analysis for a clade in which many speciose families have transcontinental distributions (e.g., Nemacheilidae, Leuciscidae, Danionidae, Cyprinidae, Ariidae, Clariidae, etc.). Numerous diverse, geographically distinct, and/or early-branching lineages, including crenuchids, cobitoids, gyrinocheiloids, diplomystoids, and ∼75% of siluroid families, are also omitted, biasing ancestral range inferences within three of five orders. Similar issues arise from the exclusion of the freshwater Denticipitoidei, an ancient freshwater lineage sister to all other clupeiforms, and of Alepocephaliformes, the marine clade sister to Ostariophysi. Limited taxon sampling artefactually recovers a single marine return in the common ancestor of plotosid and ariid catfishes, which are instead resolved as independent invasions when additional families are incorporated (49–50; Supplementary Figures SX02–13). The overrepresentation of North American taxa in Cypriniformes (8.4% of species vs 40% of tips) and Siluriformes (1.1% of species vs 20% of tips) similarly confounds biogeographic inferences within these clades (51).

Importantly, the splitting of “freshwater” into five categories also biases reconstructions to favor a marine alternative. Liu et al. (12) themselves demonstrated this artefact, recovering ∼80% support for a marine otophysan origin when using six tip states, and nearly-even support for marine and freshwater origins when using a binary (freshwater vs marine) framework (12: figs. S15 and S20).

To correct for this, we bin all taxa as freshwater, marine, or both/euryhaline, without any biogeographic considerations. While true euryhalinity is infrequent among living otophysans (occurring only in some siluroids and the diadromous cypriniform genus *Pseudaspius*) (52), it is exceptionally common in related lineages — one of three gonorynchiform suborders and a majority of clupeoid genera natively inhabit a wide range of salinities (2–3). Given the prevalence of entirely-marine (Alepocephaliformes, Gonorynchoidei) and entirely-freshwater (Denticipitoidei, Knerioidei) otocephalan clades, euryhaline transitional forms were likely more abundant amongst early otocephalans, and should be considered a plausible condition in ancestral habitat reconstructions. This character state is not permitted by Liu et al. (12) in their primary analyses, and only allowed in two supplementary analyses where several freshwater species are erroneously coded as euryhaline (Supplementary Materials, section S6). When allowing for three tip states (freshwater, marine, and brackish) and significantly overcoding extant freshwater tips as euryhaline, Liu et al. resolve a probability of less than 50% for a stenohaline marine otophysan ancestor (12: fig. S16).

We performed an assessment of ancestral habitat occupancy within Otophysa and related lineages using 84 independent BioGeoBEARS analyses (Figs 1 and 3A–G; 53). From 20 independent runs of RAxML-NG, we selected the five best-supported topologies and performed downstream analyses on each. In the case of the single best-supported topology, which resolves a functional trichotomy formed by †*Santanichthys*, Clupeiformes, and Alepocephaliformes–Ostariophysi, we calibrated the tree rooted on each of these clades and performed analyses on each chronogram independently (seven chronograms total). To account for differences in fossil sampling, we performed model-fitting with three different fossil sampling schemes on each chronogram: (i) all fossil taxa included, (ii) after removing †*Sooinichthys*, which was not yet published at the time of Liu et al.’s analyses (12), and (iii) after removing the “high missing data” taxa †*Acronichthys* and †*Nardonoides*. We then performed BioGeoBEARS on the resulting 21 trees, using two models for each: the best-fitting biogeographic model as decided by corrected Akaike Information Criterion (BAYAREALIKE+J in all cases but one; Supplementary Table TX4), and the model selected by Liu et al. (12), DEC. Finally, for each combination of chronogram, model, and fossil sampling scheme, we also performed an additional analysis with †*Santanichthys* coded as euryhaline instead of marine, consistent with its reported presence in lacustrine fossil assemblages near its type locality (30, 54).

**Fig 3.**
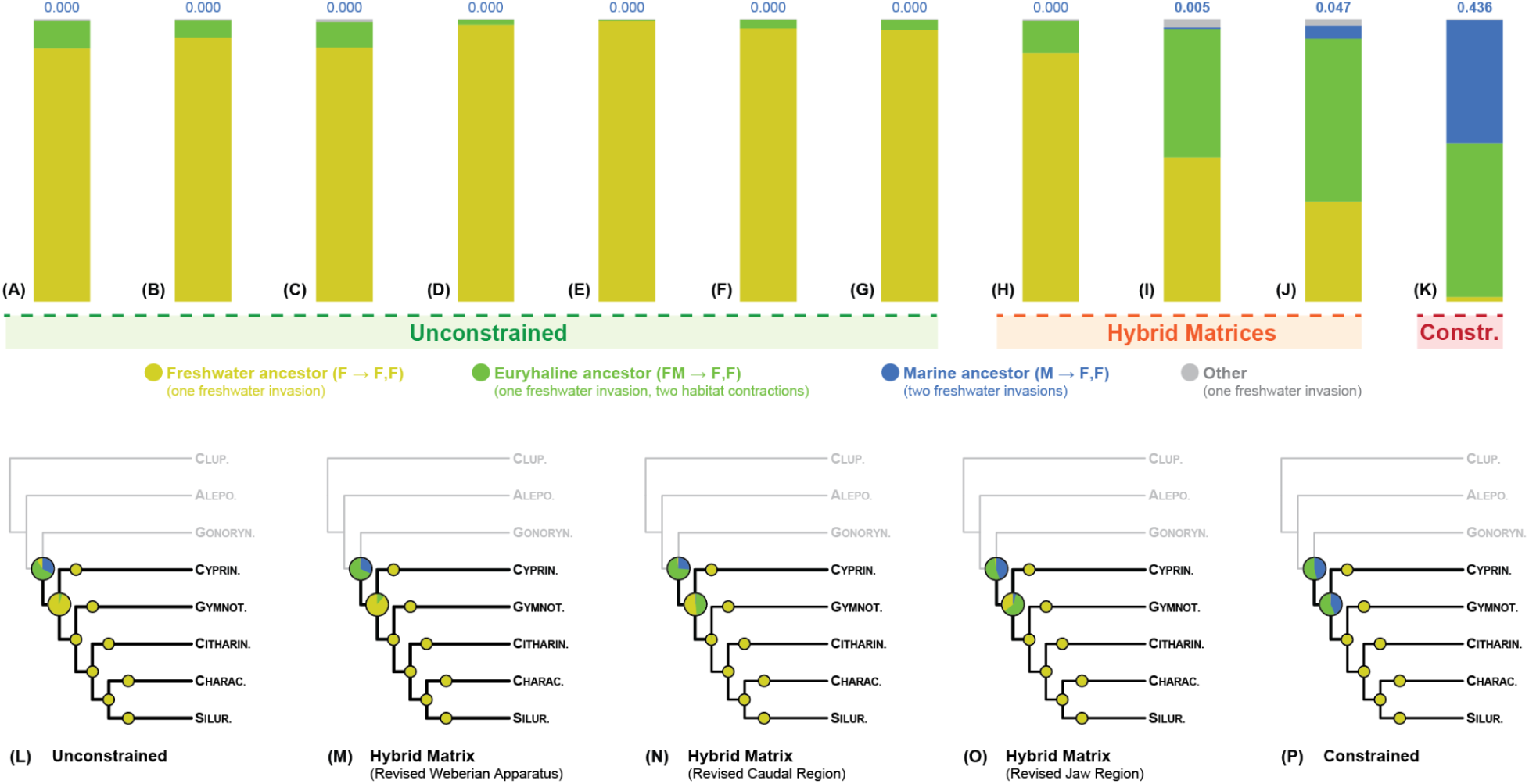
Ancestral habitat states of key nodes among extant otocephalan taxa across 132 BioGeoBEARS analyses. **(A–K)** Probabilities of different habitat transition schemes in ancient otophysans, measured as the joint probabilities of each habitat occupancy for crown Otophysa, Characiphysae, and Cypriniphysae. Yellow indicates a single marine-fresh transition (freshwater at all three nodes), green indicates a single marine-fresh transition followed by two habitat contractions (euryhaline Otophysa, freshwater Characiphysae and Cypriniphysae), and blue indicates two independent freshwater invasions (marine Otophysa, freshwater Characiphysae and Cypriniphysae). Gray indicates schemes in which crown Characiphysae and/or Cypriniphysae are resolved as euryhaline, with crown Otophysa freshwater or euryhaline (never marine). Analyses are binned by topology: **(A)** best-supported topology, rooted on Clupeiformes; **(B)** best-supported topology, rooted on Clupeiformes–†*Santanichthys*; **(C)** best-supported topology, rooted on †*Santanichthys*; **(D)** second-best topology; **(E)** third-best topology; **(F)** fourth-best topology; **(G)** fifth-best topology; **(H)** best topology recovered when recoding only the Weberian apparatus (traits 1–27); **(I)** best topology recovered when recoding only the caudal region (traits 28–41); **(J)** best topology recovered when recoding only the jaw (traits 42–49); and **(K)** best topology recovered when fossils are constrained to their placement in Liu et al. (12). Two independent invasions are not the preferred result under any topology, and only surpass 5% when fossils are constrained to their placement by Liu et al. (12). **(L–P)** Averaged probabilities of major node states across analysis types; upper and lower enlarged pie charts are the crowns of Ostariophysi and Otophysa, respectively. **(L)** compiles the average node states across all unconstrained analyses (n=84); **(M)** the average node states in the Weberian apparatus hybrid analyses (n=12); **(N)** the average node states in the caudal region hybrid analyses (n=12); **(O)** the average node states in the jaw hybrid trees (n=12); and **(P)** the average node states in the constrained analyses (n=12). Individual BioGeoBEARS results can be viewed in the Supplement (Figs. SX02–23, A–F).

Across 84 ancestral habitat reconstructions, we recover a stenohaline freshwater origin of Otophysa in all 84 trials (node probabilities between 0.541–1.00, mean 0.945) (Fig. 3A–G; Supplementary Figures SX02–15). The probability of a stenohaline marine origin never exceeds 0.001. By contrast, neither Anotophysa nor Ostariophysi can be stably resolved as any single state, though they are strongly coupled (same state at both nodes in 74 / 84 trials; Supplementary Tables TX7–8).

To investigate the sensitivity of this result, we performed 36 additional BioGeoBEARS trials on trees inferred using alternative morphological codings (Fig. 3H–J; Supplementary Figures SX16–21). Total-evidence trees estimated using the hybrid morphological matrices and the concatenated genomic dataset produced a mixed signal, with 21 trials indicating a freshwater origin of Otophysa and 15 indicating a euryhaline origin. Notably, a freshwater origin was recovered unanimously when we used the Liu et al. matrix only recoded for the Weberian apparatus. Finally, an additional 12 analyses with the fossil placements recovered by Liu et al. (12)’s ML analysis favor a euryhaline (6 trials) or marine (6 trials) origin of Otophysa, dependent on fossil sampling, BioGeoBEARS model, and †*Santanichthys* habitat coding. Across these twelve analyses, which adopt the exact fossil placements recovered by Liu et al. (12), only half support a stenotopic marine origin of Otophysa, and in no case does a stenotopic marine origin attain 90% support. In sum, across all 132 BioGeoBEARS analyses, we recover a freshwater origin of Otophysa in 106, a euryhaline origin in 20, and a marine origin in only 6, all of which enforce the fossil placements of Liu et al. (12).

The reconciliation of phylogenetically inferred divergence times with paleogeographic events remains a controversial undertaking (17, 55–60). This approach was used by Liu et al. (12) to contend that a mid-Jurassic divergence between cypriniphysean and characiphysean otophysans — one that postdates the beginning of the Pangaean breakup — requires two independent freshwater invasions. This claim hinges upon the accurate estimation of divergence times, as well as the presence of a completely impermeable geographic barrier between the nascent continents of Gondwana and Laurasia.

Phylogenetic estimates of the crown otophysan age are consistently between 145 and 175 Ma when using large nuclear datasets and dense fossil sampling (7, 31–32, 34, 61). This study similarly resolves a cypriniphysean-characiphysean split during the Jurassic, amidst the Pangaean breakup (roughly 200 to 140 Ma) (28, 62–63) (Fig. 1). While Gondwana and Laurasia grew increasingly disconnected during this time, they remained ecologically interconnected by ephemeral land bridges in the present-day North Atlantic, as late as the Barremian (c. 126–121 Ma, 64). These marginal habitats, although inhospitable to most modern-day otophysans, provide evidence that the nascent barriers between protocontinents would have remained at least transiently permeable to freshwater fishes throughout the Jurassic, until such land bridges completely disappeared (64).

The viability of a single freshwater invasion by otophysan fishes is also not dependent on the chronology of their diversification, as modern otophysans exhibit many widely-accepted examples of transoceanic dispersal. Within Leuciscidae (minnows and kin), two North American lineages diverged from their Eurasian relatives only 15–25 Ma ago (31, 65), contemporary with the separation between North American and Chinese catostomids (31, 65) but long after land connectivity between the two landmasses had dissipated (66). Siluroid catfishes — which famously harbor freshwater clades endemic to the Americas, Eurasia, India, Africa, Australia–New Guinea, and Madagascar — radiated long after the disintegration of Pangaea (31, 61, 67), although virtually none can survive in seawater for prolonged periods. An especially pertinent example is the rare genus *Lacantunia*, a Central American endemic deeply nested among African families (68). Irrespective of otophysan crown state, Liu et al.’s own chronogram requires transoceanic dispersals within four of the five otophysan orders (12). Though instances of such dispersal are likely rare, the assumption that marine barriers firmly preclude long-distance dispersal in freshwater fishes is untenable, and dispelled by Liu et al.’s own analyses (12).

Inferring major biogeographic events in vertebrate lineages is compelling but inherently difficult. The fossil record, especially for fishes and even more so for freshwater fishes, is fragmentary by nature, and taphonomic biases can easily obscure evolutionary history to the detriment of otherwise well-founded analyses. By applying a more conservative osteological interpretation of key otophysan fossils, increasing taxon and marker sampling, using less restrictive ancestral state coding, and applying best-fitting biogeographic models, we show how poorly preserved marine taxa overrepresented in the fossil record can distort ancestral habitat reconstructions. We do not argue that marine origins for major freshwater radiations are implausible in principle, only that any such claim must be robust to the methodological choices that produced it.

## Acknowledgements

The authors wish to thank Zach Heiple (UCSD, SIO) and Aline Paiva Medeiros (OU, UCSD, SIO) for useful insights in performing biogeographic analyses; Milton Tan (UIUC) for discussions of otophysan systematics and phylogenetics; and Guillermo Ortí (GWU) and Lily C. Hughes (NCMNS) for their leadership in the FishLife project, which provided much of the data used in these analyses. We would like to thank the curators and institutions for allowing us to consult fossils: Jesús Alvarado (National Autonomous University of Mexico), Gaël Clément (Muséum National d’Histoire Naturelle, Paris), Zerina Johanson (the Natural History Museum, London), John Maisey (American Museum of Natural History), and Anna Vaccari (Museo Civico di Storia Naturale di Verona). Additional thanks to the former and current members of the Scripps Institution of Oceanography Fish Evolution Lab, and to the 2023 cohort of the Summer Systematics Institute (SSI) at the California Academy of Sciences (CAS), for their continued and enthusiastic support. Analyses by D.H.H. were performed on the OSCER High Performance Computing cluster at Oklahoma University and CIPRES, hosted by the University of California San Diego. Analyses by M.A.B and A.B. were performed on the Pegasus High Performance Computing Cluster at the George Washington University, as well as through the TNT GUI made freely available thanks to the Willi Hennig Society.

## Author contributions

Conceptualization: R.B.R., O.O., and D.M.; Methodology: D.H.H., M.A.B., A.S., D.A., O.O., D.M., and R.B.R.; Investigation: D.H.H., M.A.B., A.S., A.B., O.O., and D.M.; Visualization: D.H.H., M.A.B., A.B., O.O, and D.M.; Project administration: R.B.R., A.S., and D.A.; Supervision: R.B.R. and A.S.; Writing — original draft: D.H.H., M.A.B., O.O., and D.M.; Writing — review & editing: D.H.H., M.A.B, A.S., D.A., A.B., O.O., D.M., and R.B.R.

## Supplementary Materials

### S0. Fossil material examined

Most of the character states coded for the fossil taxa are taken from published articles (8, 12, 20, 25, 30). For four species (all recoded taxa other than †*Acronichthys maccagnoi*), additional personal observations were made by DM based on the following specimens:

†*Chanoides macropoma*: The material examined includes four specimens housed in the Natural History Museum, London (NHMUK PV P.1867; P.1868b; P.28397; P.51216), one housed in the American Museum of Natural History (AMNH 4108), and 11 specimens deposited in the Muséum National d’Histoire Naturelle, Paris (MNHN Bol 0077; Bol 0078; Bol 0464; Bol 0465; Bol 0443; Bol 0444; Bol 0447; Bol 0468; Bol 0469; Bol 0500; Bol 0504).

†*Nardonoides chardoni*: The material examined is housed in the collections of the Museo Civico di Storia Naturale di Verona (MCSNV Na 500; Na 62; Na 108; Na 109; Na 173; Na 174; Na 175; Na 214; Na 8; Na 176; Na 197; Na 204; Na 205; Na 213; Na 215; Na 216; Na 217; Na 219; Na 246; Na 517; Na 523; Na 525).

†*Santanichthys diasii*: The material examined is housed in the American Museum of Natural History (specimens AMNH 12826; AMNH 13981, a mass mortality plate featuring specimens labelled from A to Z; AMNH 19439; AMNH 22652 (originally, 11922); and AMNH 22654 (originally 11954)).

†*Sooinichthys varii*: The material examined is housed in the Colección Nacional de Paleontología at the Instituto de Geología, Universidad Nacional Autonoma de México (specimens IGM 14347 and IGM 14348–14377).

### S1. Morphology-only phylogenetic tree inference

The revised morphological matrix for these studies was modified from Liu et al. (12) and included 63 characters and 20 taxa. One of these taxa, †*Sooinichthys varii*, is included in a phylogenetic context for the first time. Four of these taxa,†*Nardonoides chardoni*, †*Chanoides macropoma*, †*Santanichthys diasii*, and †*Acronichthys maccagnoi* were recoded as described above. The matrix comprised four character suites: features of the Weberian Apparatus, features of the caudal fin, features of the jaws, and additional features not corresponding to a unified anatomical region. All analyses were run without partitioning the data.

For maximum parsimony in TNT (44, 45), we performed exhaustive searches via iterative enumeration under both equal weighting and a comprehensive range of implied character weights (*k* = 1-10,12,15,20,50). Implied weighting acts to downweight homoplastic characters, with larger concavity constants (*k*) producing results increasingly similar to equal weighting (75). All most parsimonious trees for a weighting scheme were summarized using strict consensus.

For model-based frameworks, we implemented the Mk model under maximum likelihood in IQ-Tree (47) and Bayesian inference in MrBayes (48). For maximum likelihood, 1000 ultrafast bootstraps (77) were used to assess support. For Bayesian inference, posterior probabilities were estimated using MCMC with 650,000 generations with a 25% burn-in, four metropolis-coupled chains, and a heating parameter of 0.2. Trees and parameter values were sampled every 100 generations, and convergence diagnostics were calculated every 1,000 generations.

Trees were estimated under each optimality criterion in two ways: first without topological constraints and second with a constraint matching the topology of Liu et al. (12) for topological hypothesis testing.

Constrained and unconstrained parsimony trees were compared based on tree length. Maximum likelihood trees were compared using an approximately biased (AU) test (46). Using our revised morphological matrix under the Mk model, statistical support for the alternative topologies was assessed using 10,000 resampled estimated log-likelihood bootstrap replicates. In addition, we conducted an AU test using the unmodified Liu et al. matrix (12) under the same parameters.

For Bayesian Inference trees, we calculated Bayes factors (BF) via marginal likelihoods estimated using a stepping-stone sampler in MrBayes (50 steps, 1,000,000 generations, 25% burn-in) (97). While the Liu et al. constraint was initially favored over our unconstrained topology, this was artefactual due to the unconstrained run searching an exponentially larger prior tree space which in turn penalizes the marginal likelihood (70, 88). To correct for this, we constrained a run to match the unconstrained maximum clade credibility (MCC) topology.

Three combination matrices were created by substituting recoded character suites into the original Liu et al. matrix (12), one for each character suite: weberian apparatus, caudal, jaws. For each of these matrices, as well as the unmodified Liu et al. matrix, unconstrained trees were inferred for each optimality criterion, following the same parameters as above.

### S2. Total-evidence phylogenetic tree inference

To interrogate historical habitat transitions within Otophysa, we estimated a new total-evidence phylogeny of Otocephala with broadened taxonomic sampling. Our alignment was assembled from three components: a 63-character morphological matrix revised from Liu et al. (12), the 95,274-bp UCE alignment from Chakrabarty et al (1) as used by Liu et al., and a novel 616,966-bp alignment using the “FishLife” exon set of Hughes et al (37). To produce this exonic alignment, we produced *de novo* assemblies from 27 sets of raw genomic reads (Supplementary Table TX2) using MEGAHIT v1.2.9 (81). We then used HMMER v3.4 (96) to mine 1,123 nuclear and 10 mitochondrial exons from each MEGAHIT assembly as well as 52 publicly-available genomes from the NCBI genome database (Supplementary Table TX2). Exons were then aligned using MAFFT v7.505 (79). This expanded alignment allowed us to incorporate exonic data for all 36 tips in the original Liu alignment, as well as 26 families (including the major lineages Alepocephaliformes, Cobitoidei, Denticipitoidei, and Diplomystoidei) not present in their analyses. The three matrices (one morphological and two molecular) were concatenated using AMAS (71).

In order to prevent data non-overlap, we treat 17 tips as composite taxa, with UCEs and exons from two different congeneric (n = 15) or confamilial (n = 2) species (Supplementary Table TX2). To ensure no violation of monophyly assumptions, all composite taxa are lone representatives of stable taxonomic lineages. 13 composite tips enclose the only included data for stable monophyletic families (Sternopygidae, Ctenoluciidae, Bryconidae, Characidae, Erythrinidae, Loricariidae, Pangasiidae, Ictaluridae, Mochokidae, Malapteruridae, Ariidae, and Auchenipteridae, respectively). Within Dorosomatidae, the two species of the “*Dorosoma*” composite form a stable North American clade to the exclusion of the Indo-Pacific *Herklotsichthys* and *Tenualosa* (74, 90). Within Catostomidae, the two species of the “*Moxostoma*” composite form a stable North American clade to the exclusion of the China-endemic *Myxocyprinus* (69, 84). Within Distichodontidae, the species pairs of the “*Xenocharax*” and “*Distichodus*” composites form morphologically and ecologically distinct clades across several previous analyses (80, 93), with *Xenocharax* representing the first-branching distichodontid genus. Within Alestidae, the two species of the “*Bryconaethiops*” composite form a stable clade to the exclusion of *Brycinus* (86, 94), which is the first-branching alestid lineage.

Total-evidence phylogenetic trees were inferred using RAxML-NG v1.0.2 (38). We generated an initial pair of trees using the modified matrix, one in which species with morphological data were constrained to their placements in Liu et al., and one with no topological constraint. We repeated this with the original matrix of Liu et al., first unconstrained and then again constrained to their placement following the unconstrained, revised-matrix phylogeny. We repeated this once more with three “hybrid” matrices, each with no constraint, and using the original Liu et al. matrix but with our revised coding for only (a) the Weberian apparatus, (b) the caudal fin, or (c) the jaw. All runs of RAxML-NG used five partitions: one for the morphological matrix (MULTI4_Mk), one for the UCE dataset, and one for each codon position of the exonic dataset.

### S3. Divergence time estimation

We used BEAST v2.7.8 (40) to time-calibrate eleven RAxML topologies under the Fossilized Birth Death (FBD) model and a Bayesian framework (39). We calibrated all three unconstrained trees recovered using hybrid matrices, plus the constrained topology and the five best supported unconstrained topologies recovered using revised matrix. Because the single best supported unconstrained tree did not conclusively resolve †*Santanichthys* as a member of either Clupeomorpha or Alepocephaliformes—Ostariophysi we calibrated this topology with three alternative rooting schemes: (a) on †*Santanichthys*, (b) on †*Santanichthys*—Clupeomorpha, and (c) on Clupeomorpha, producing a total of seven chronograms. The primary analysis is considered (c), owing to a higher number of morphological traits suggesting an alepocephaliform-ostariophysan affinity for †*Santanichthys*, as compared to a clupeomorph affinity (Table TX6).

In all cases, we used the appropriate matrix (revised or hybrid) and a representative selection of 75 exons, determined by their proportion of parsimony informative sites and partitioned by codon position, as input to BEAST (40,392 bp in total). A generalized time reversible (GTR) model was applied to the molecular dataset, and a Lewis Mk model was applied to the morphological dataset. An optimized relaxed clock (ORC) was applied to all partitions. Each fossil taxon was calibrated using a tip age prior corresponding to its youngest fossil appearance, including two revised from Liu et al. 14 node age priors were applied with a uniform distribution each. 13 were derived from a range of literature ages for the corresponding node (Supplementary Table TX3), while a fourth (*Chirocentrus* + †*Diplomystus*) was applied to account for the oldest fossil record of †*Diplomystus*. A fifteenth node age prior, corresponding to Clupeocephala, was applied to the topology rooted on †*Santanichthys*.

In order to constrain BEAST to produce the same topology as RAxML-NG, we first performed a provisional calibration of each phylogram using the same set of node-dates, and the *chronos* function in the R package phyTools (90). We then provided each provisional (ultrametric) “chronogram” as a starting topology to BEAST and disabled seven operators that control topology changes during branch length estimation. This allowed us to use a sequence-aware, tip-dating approach to accurately calibrate our query trees without allowing for any topological changes due to methodological (Bayesian vs maximum likelihood) or sampling (75 vs 1,133 loci) effects. We ran BEAST for 10,000,000 iterations per run. After tree estimation, TreeAnnotator v2.7.7 (40) was used to produce a maximum clade credibility (MCC) tree for each of the seven runs, using mean node heights and a burn-in fraction of 0.10.

### S4. Calibration justification

1. Tip-date: †*Acronichthys*. This genus is known from the late-Maastrichtian Pisces Point Locality (Scollard Formation, Alberta, Canada), dated to be 66.9 Ma old by Liu et al. (12). We retain this date from Liu et al.
2. Tip-date: †*Chanoides*. This genus is known from the Monte Bolca Lagerstätte (Region of Veneto, Italy), for which we use the Ypresian lower bound of 48.1 Ma (25). We retain this date from Liu et al.
3. Tip-date: †*Diplomystus*. This genus is best known for the Green River form, †*Diplomystus dentatus*, dated to roughly 48.1 Ma (92). However, a younger taxon †*Diplomystus shengliensis* has also been identified as a true member of this genus using parsimony analyses (68, 73). The youngest †*Diplomystus* are therefore from the Shahejie Formation (Shandong, People’s Republic of China), for which we use the Bartonian lower bound of 37.7 Ma. **Modified from Liu et al.**
4. Tip-date: †*Hypsidoris*. This genus contains two species, the younger of which, †*Hypsidoris oregonensis*, is known from the Clarno Formation (Oregon, United States of America; 76), for which we use the Bartonian lower bound of 37.7 Ma. We retain this date from Liu et al.
5. Tip-date: †*Jianghanichthys*. This genus holds two species, the younger of which, †*Jianghanichthys sanshuiensis*, is known from the Buxin Formation (Guangdong, People’s Republic of China; 82), for which we use the Ypresian lower bound of 48.1 Ma. We retain this date from Liu et al.
6. Tip-date: †*Nardonoides*. This genus is known from the Nardò Formation (Apulia, Italy; 25), for which we use the Campanian lower bound of 72.2 Ma. We retain this date from Liu et al.
7. Tip-date: †*Rubiesichthys*. This genus is known from the Las Hoyas Lagerstätte / Las Huerguinas Formation (Castilla–La Mancha, Spain; 89), for which we use the Barremian lower bound of 121.3 Ma. The age of this fossil is not consistent among the trees presented by Liu et al. (12, Figs S13–S16 vs. S18). **Modified from Liu et al.**
8. Tip-date: †*Santanichthys*. This genus is known from the Santana Locality / Romualdo Formation (Ceará, Brazil, 30), for which we use the Aptian lower bound of 113.2 Ma. We retain this date from Liu et al.
9. Tip-date: †*Sooinichthys*. This genus is known from the Tlayúa Quarry (Puebla, Mexico), for which we use the Albian lower bound of 100.5 Ma (20). **Not included by Liu et al.**
10. Node-date: crown Otomorpha. MRCA of *Denticeps*, *Danio*, and †*Santanichthys*. Uniform distribution from 230.5 – 229.5 Ma, taken from the age recovered by (31).
11. Node-date: crown Clupeoidei. MRCA of *Jenkinsia* and *Alosa*. Uniform distribution from 94.9 Ma (32) to 105.8 Ma (61).
12. Node-date: MRCA of *Chirocentrus* and †*Diplomystus*. Implemented to accommodate the recent rootward extension of the †*Diplomystus* fossil record. Two marine species (†*D. birdi* and †*D. dubertreti*) not considered by Liu et al. (12) have since been recovered as Cretaceous members of this genus using maximum parsimony analyes (85, 87). The older of these two, †*D. birdi*, is known from the Sannine Formation (Keserwan-Jbeil Governorate, Lebanon), for which we use the age of 93.9 Ma. Uniform distribution from 93.9 Ma (youngest age of †*D. birdi*) to 94.9 Ma (the age recovered by (32) for Clupeoidei).
13. Node-date: crown Alepocephaliformes. MRCA of *Alepocephalus* and *Holtbyrnia*. Uniform distribution from 24.3 Ma (34) to 53.3 Ma (31).
14. Node-date: crown Ostariophysi. MRCA of *Chanos* and *Danio*. Uniform distribution from 160.6 Ma (32) to 210.0 Ma (34).
15. Node-date: crown Anotophysa. MRCA of *Gonorynchus* and *Chanos*. Uniform distribution from 150.0 Ma (7) to 183.3 Ma (34).
16. Node-date: crown Otophysa. MRCA of *Gymnotus* and *Danio*. Uniform distribution from 146.8 Ma (32) to 213.3 Ma (7).
17. Node-date: crown Cypriniformes. MRCA of *Moxostoma* and *Danio*. Uniform distribution from 95.9 Ma (34) to 158.9 Ma (7).
18. Node-date: crown Characiphysae. MRCA of *Gymnotus* and *Silurus*. Uniform distribution from 127.3 Ma (32) to 183.5 Ma (7).
19. Node-date: crown Cithariniformes. MRCA of *Citharinus* and *Distichodus*. Uniform distribution from 76.2 Ma (61) to 108.6 Ma (31).
20. Node-date: crown Gymnotiformes. MRCA of *Sternopygus* and *Gymnotus*. Uniform distribution from 71.0 Ma (7) to 89.4 Ma (31).
21. Node-date: crown Characiformes. MRCA of *Crenuchus* and *Brycon*. Uniform distribution from 89.9 Ma (32) to 132.6 Ma (31).
22. Node-date: crown Siluriformes. MRCA of *Ancistrus* and *Ariopsis*. Uniform distribution from 94.9 Ma (32) to 120.4 Ma (31).
23. Node-date: crown Siluroidei. MRCA of *Silurus* and *Ariopsis.* Uniform distribution from 70.0 Ma (32) to 86.2 Ma (7).

### S5. Ancestral habitat reconstruction

We analyzed habitat shifts in ancestral otophysans using the R package BioGeoBEARS (53), which incorporates model-fitting into biogeographic analyses. For maximum simplicity, we considered only two habitat states, freshwater and marine, with several tips present in both (euryhalinity; n = 13 or 14). We tested 6 biogeographic models: DEC, DIVALIKE, and BAYAREALIKE, each with and without the *J* parameter, which permits “jump” dispersal (i.e., allows for parent and daughter nodes to inhabit non-overlapping habitats). For each BioGeoBEARS trial, we calculated model fits using log likelihood and AIC criterion, as well as weights of *D, E,* and *J* parameters for each. The best fitting model was recorded for each of 66 combinations of topology and fossil sampling, and ancestral habitat reconstruction was performed using both the best-fit model (BAYAREALIKE+J in 65 out of 66 cases) as well as the model selected by Liu et al. (DEC; Supplementary Table TX4; the best fit model in 1 out of 66 cases).

A total of 132 ancestral habitat reconstructions were performed (Supplementary Figures SX02–13). Our primary hypothesis uses the single best supported tree from our RAxML-NG analysis, rooted on Clupeiformes, with all fossils included, the best-fitting BioGeoBEARS model (BAYAREALIKE+J), and †*Santanichthys* coded as marine. To confirm the sensitivity of this result to different sources of uncertainty, we replicated this analysis with †*Santanichthys* coded as euryhaline, consistent with the Codó Formation (considered lacustrine by Lindoso et al, 54), and again using the DEC model, following Liu et al. (12). To account for the inclusion of †*Sooinichthys varii*, which was not available for consideration at the time of Liu et al.’s analyses, we also performed reconstructions without this tip. To additionally account for two tenuously-placed fossil taxa with high missing data, †*Acronichthys maccagnoi* and †*Nardonoides chardoni*, we performed additional reconstructions lacking these two tips. We ran trials with each combination of †*Santanichthys* habitat coding (n=2), fossil tip inclusion scheme (n=3), and biogeographic model (n=2), for a total of 12 trials on our primary phylogeny.

Because this topology resolves †*Santanichthys diasii* in an ambiguous placement with respect to Clupeiformes and Alepocephaliformes–Ostariophysi, we repeated this set of twelve ancestral reconstructions with alternative rooting schemes (either on †*Santanichthys diasii*, or on Clupeiformes + †*Santanichthys diasii*; 24 additional trials total). In order to account for further instability in fossil placement, we also calibrated the second-, third-, fourth-, and fifth-best supported topologies recovered from our RAxML-NG analysis, in all cases rooted on Clupeiformes. The same twelve parameter combinations were applied to each topology (48 additional trials).

To further evaluate the impact of morphological character interpretation on ancestral habitat reconstruction, we performed the same set of twelve BioGeoBEARS trials using each of the three “hybrid” matrix chronograms (36 additional trials). Finally, we performed the same set of twelve ancestral habitat reconstructions on a topology constrained to that of Liu et al. (12 additional trials).

### S6. Habitat coding justification

#### Fossil Taxa

†*Acronichthys*: **Fresh**. Known from the Scollard Formation in Canada, a freshwater habitat following Liu et al. (12).

†*Chanoides*: **Marine**. Known from the Monte Bolca Lagerstätte in Italy, a shallow marine ecosystem (25).

†*Diplomystus*: **Euryhaline**. While the type species (†*D. dentatus*) is known from the freshwater Green River Formation in the United States of America, additional species are known from the marine Sannine and Sahel Alma Formations in Lebanon. Parsimony-based analyses (85) have demonstrated these forms to be congeneric. Following these findings, we considered this a euryhaline genus. **Modified from Liu et al.** (**12**), who consider this genus freshwater in all analyses.

†*Hypsidoris*: **Fresh**. Both species are known only from freshwater localities in the United States of America (Green River and Clarno Formations; 76).

†*Jianghanichthys*: **Fresh**. Both species are known only from freshwater localities in the People’s Republic of China (Buxin and Yangxi Formations; 82).

†*Nardonoides*: **Marine**. Known from the Nardò Formation in Italy, a shallow marine ecosystem (25).

†*Rubiesichthys*: **Euryhaline**. Known from the Las Hoyas Lagerstätte / Las Huerguinas Formation in Spain, considered to be lacustrine mudstone deposits, and also the marine deposits of El Montsec (89). **Modified from Liu et al.**, who consider this genus freshwater in all analyses.

†*Santanichthys*: **Marine, or Euryhaline**. Type material collected from the Santana Locality / Romualdo Formation in Brazil, considered to be coastal marine (30). In keeping with this, and the interpretation put forward by Liu et al., we perform half of our ancestral habitat reconstructions (66 / 132 trials) under the assumption that this species is stenohaline marine. Additional specimens purportedly occur in the Maruim Member of the Riachuelo Formation (considered marine, continental slope by Filleul and Maisey, 30) and the Codó Formation (considered lacustrine by Lindoso et al, 54), both of Brazil. These additional reports are described in less detail but indicate that †*Santanichthys* could plausibly have been euryhaline, instead of stenohaline marine. We perform the other half of our analyses under this alternative consideration.

†*Sooinichthys*: **Marine**. Known from the Tlayúa Quarry, a lagoonal Lagerstätte considered marine by Alvarado-Ortega et al (20).

#### Clupeiforms

*Alosa*: **Euryhaline**. Species considered [freshwater] or [freshwater, brackish] or [freshwater, brackish, marine] or [brackish] or [brackish, marine] by (2) and/or (3). Some taxa are anadromous, and as such we consider this a euryhaline genus.

*Brevoortia*: **Euryhaline**. Species considered [freshwater, brackish, marine] or [brackish, marine] or [marine] by (2) and/or (3). We consider this a euryhaline genus.

*Chirocentrus*: **Marine**. All species marine following (2) and (3). *C. dorab* is considered [brackish, marine] by both sources. Because these species are primarily marine and never freshwater, we consider this a marine stenohaline genus. Liu et al. code this genus as marine in Figures 3 and S15, but euryhaline in Figure S16.

*Clupea*: **Euryhaline**. Species considered [freshwater, brackish, marine] or [brackish, marine] by both sources. We consider this a euryhaline genus.

*Clupeonella*: **Euryhaline**. Species considered [freshwater] or [brackish, marine] by both sources. We consider this a euryhaline genus.

*Coilia*: **Euryhaline**. Species considered [freshwater] or [freshwater, brackish] or [freshwater, brackish, marine] or [brackish, marine] by both sources. Some taxa are anadromous. We consider this a euryhaline genus.

*Corica*: **Fresh**. All species freshwater following (2) and (3). *C. laciniata* is considered [freshwater, brackish] by both sources, and *C. soborna* is considered [freshwater, brackish, marine] by (3). Because these species are primarily freshwater and only dubiously occur in marine waters, we consider this a freshwater stenohaline genus.

*Denticeps*: **Fresh**. Monotypic, sole species freshwater following (2) and (3).

*Dorosoma*: **Euryhaline**. Species considered [freshwater] or [freshwater, brackish] or [freshwater, brackish, marine] by (2) and/or (3). Some taxa are anadromous, and as such we consider this a euryhaline genus. **Modified from Liu et al.** (**12**), who code this genus as freshwater in Figures 3 and S19, marine in Figures S15 and S20, and euryhaline in Figure S16.

*Dussumieria*: **Marine**. All species marine following (2) and (3).

*Encrasicholina*: **Marine**. All species marine following (2) and (3). *E. purpurea* is considered [brackish, marine] by both sources. Because these species are primarily marine and never freshwater, we consider this a marine stenohaline genus.

*Engraulis*: **Marine**. All species marine following (2) and (3). *E. encrasicolus* is considered [brackish, marine] by both sources. Because these species are primarily marine and never freshwater, we consider this a marine stenohaline genus.

*Etrumeus*: **Marine**. All species marine following (2) and (3).

*Herklotsichthys*: **Euryhaline**. Species considered [freshwater, brackish, marine] or [brackish, marine] or [marine] by (2) and/or (3). Some taxa are anadromous, and as such we consider this a euryhaline genus.

*Ilisha*: **Marine**. This genus is potentially polyphyletic and contains freshwater, euryhaline, and marine lineages. We examine only *I. melastoma*, which is considered [brackish, marine] by both (2) and (3). Because this species is primarily marine and never freshwater, we consider this a marine stenohaline genus as represented within the present study.

*Jenkinsia*: **Marine**. All species marine following (2) and (3).

*Spratelloides*: **Marine**. All species marine following (2) and (3). Two species (*S. delicatulus* and *S. robustus*) are considered [brackish, marine] by one or both sources. Because these species are primarily marine and never freshwater, we consider this a marine stenohaline genus.

*Sundasalanx*: **Fresh**. All species freshwater following (2) and (3).

*Tenualosa*: **Euryhaline**. Species considered [freshwater] or [freshwater, brackish, marine] by (2) and/or (3). Some taxa are anadromous, and as such we consider this a euryhaline genus.

*Thryssa*: **Euryhaline**. Many species considered [brackish, marine] or [marine] by (2) and/or (3), while others are considered [freshwater] by one or both. We consider this a euryhaline genus. **Modified from Liu et al.** (**12**), who code this genus as marine in Figures 3, S15, S19, and S20, and euryhaline in Figure S16.

#### Alepocephaliforms

*Alepocephalus*: **Marine**. All species marine following (2) and (3).

*Bajacalifornia*: **Marine**. All species marine following (2) and (3).

*Holtbyrnia*: **Marine**. All species marine following (2) and (3).

*Normichthys*: **Marine**. All species marine following (2) and (3).

#### Anotophysans

*Chanos*: **Euryhaline**. Monotypic, sole species considered [freshwater, brackish, marine] by both (2) and (3), with a catadromous life history. We consider this a euryhaline genus. **Modified from Liu et al.** (**12**), who consider this genus marine in Figures 3, S15, S19, and S20, and euryhaline in Figure S16.

*Gonorynchus*: **Marine**. All species marine following (2) and (3). *G. greyi* is considered [freshwater, brackish, marine] by (2), but [brackish, marine] by (3). Because these species are primarily marine and only dubiously occur in fresh waters, we consider this a marine stenohaline genus.

*Parakneria*: **Fresh**. All species freshwater following (2) and (3).

*Phractolaemus*: **Fresh**. All species freshwater following (2) and (3).

#### Otophysans

*Ageneiosus* / *Trachycorystes*: **Fresh**. All species freshwater following (2) and (3).

*Ameiurus*: **Fresh**. All species freshwater following (2) and (3).

*Ancistrus*: **Fresh**. All species freshwater following (2) and (3).

*Apareiodon*: **Fresh**. All species freshwater following (2) and (3).

*Apteronotus*: **Fresh**. All species freshwater following (2) and (3).

*Ariopsis*: **Euryhaline**. Species are considered [freshwater, brackish] or [freshwater, brackish, marine] or [brackish, marine] or [marine] by (2) and (3). Because these species may be found regularly in both freshwater and marine habitats, we consider this a euryhaline genus. **Modified from Liu et al.** (**12**), who consider this genus marine in Figures 3, S15, S19, and S20, and brackish in Figure S16.

*Astyanax*: **Fresh**. All species freshwater following (2) and (3). **Modified from Liu et al.** (**12**), who consider this genus freshwater in Figures 3, S15, S19, and S20, but euryhaline in Figure S16.

*Auchenoglanis*: **Fresh**. All species freshwater following (2) and (3).

*Brachyhypopomus*: **Fresh**. All species freshwater following (2) and (3). *B. janeiroensis* is considered [freshwater, brackish] by (2), but not (3). Because this species is primarily freshwater and never marine, we consider this a freshwater stenohaline genus.

*Brycinus*: **Fresh**. All species freshwater following (2) and (3). **Modified from Liu et al.** (**12**), who consider this genus freshwater in Figures 3, S15, S19, and S20, but euryhaline in Figure S16.

*Brycon*: **Fresh**. All species freshwater following (2) and (3).

*Bryconaethiops*: **Fresh**. All species freshwater following (2) and (3).

*Channallabes*: **Fresh**. Monotypic, sole species freshwater following (2) and (3).

*Citharinus*: **Fresh**. All species freshwater following (2) and (3).

*Clarias*: **Fresh**. All species freshwater following (2) and (3). *C. gariepinus* is considered [freshwater, brackish] by (2) but not (3). Because these species are primarily freshwater and never marine, we consider this a freshwater stenohaline genus.

*Colossoma*: **Fresh**. Monotypic, sole species freshwater following (2) and (3).

*Corydoras*: **Fresh**. All species freshwater following (2) and (3).

*Cranoglanis*: **Fresh**. All species freshwater following (2) and (3).

*Crenuchus*: **Fresh**. All species freshwater following (2) and (3).

*Ctenolucius*: **Fresh**. All species freshwater following (2) and (3).

*Cyprinella*: **Fresh**. All species freshwater following (2) and (3).

*Cyprinus*: **Fresh**. All species freshwater following (2) and (3). Some species (*C. acutidorsalis, C. carpio, C. chilia, C. melanes,* and *C. rubrofuscus*) are considered [freshwater, brackish] by one or both sources. Because these species are primarily freshwater and never marine, we consider this a freshwater stenohaline genus.

*Danio*: **Fresh**. All species freshwater following (2) and (3).

*Diplomystes*: **Fresh**. All species freshwater following (2) and (3).

*Distichodus*: **Fresh**. All species freshwater following (2) and (3).

*Gogo*: **Fresh**. All species freshwater following (2) and (3).

*Gymnotus*: **Fresh**. All species freshwater following (2) and (3).

*Hepsetus*: **Fresh**. All species freshwater following (2) and (3).

*Hoplias*: **Fresh**. All species freshwater following (2) and (3).

*Labeo*: **Fresh**. All species freshwater following (2) and (3). Some species (*L. calbasu, L. catla,* and *L. rohita*) are considered [freshwater, brackish] by one or both sources. Because these species are primarily freshwater and never marine, we consider this a freshwater stenohaline genus.

*Leptobotia*: **Fresh**. All species freshwater following (2) and (3).

*Leuciscus*: **Fresh**. All species freshwater following (2) and (3). Some species (*L. aspius*, *L. burdigalensis*, *L. idus, L. leuciscus, L. oxianus, L. vorax,* and *L. waleckii*) are considered [freshwater, brackish] by one or both sources. Because these species are primarily freshwater and never marine, we consider this a freshwater stenohaline genus.

*Malapterurus*: **Fresh**. All species freshwater following (2) and (3).

*Megaleporinus*: **Fresh**. All species freshwater following (2) and (3).

*Misgurnus*: **Fresh**. All species freshwater following (2) and (3). *M. fossilis* is considered [freshwater, brackish] by both sources. Because this species is primarily freshwater and never marine, we consider this a freshwater stenohaline genus.

*Moxostoma*: **Fresh**. All species freshwater following (2) and (3).

*Myxocyprinus*: **Fresh**. Monotypic, sole species freshwater following (2) and (3).

*Nannaethiops*: **Fresh**. All species freshwater following (2) and (3).

*Pangasius*: **Fresh**. All species freshwater following (2) and (3). *P. mekongensis* is considered [freshwater, brackish] by both sources. A second species (*P. myanmar*) is considered [freshwater, brackish, marine] by (2), but [freshwater] by (3). A third species (*P. krempfi*) is considered [freshwater, brackish, marine] by both, and is anadromous. This is a derived condition limited to *P. krempfi*, and we consider it unrepresentative of the genus (molecular data compiled from *P. djambal* and *P. pangasius*, both considered [freshwater] by both sources). We consider this a freshwater stenohaline genus. **Modified from Liu et al.**, who consider this genus freshwater in Figures 3, S15, S19, and S20, but euryhaline in Figure S16.

*Plotosus*: **Marine**. All species marine following (2) and/or (3). One species (*P. abbreviatus*) is considered [brackish] by (2) but [brackish, marine] by (3). A second species (*P. papuensis*) is considered [freshwater] by both sources, a derived condition which we consider unrepresentative of the genus. Some additional species (*P. canius, P. limbatus,* and *P. nkunga*) are considered [freshwater, brackish, marine] by both sources. Because these species are almost all marine and less regularly occur in freshwaters, we consider this a marine stenohaline genus.

*Prochilodus*: **Fresh**. All species freshwater following (2) and (3). *P. vimboides* is considered [freshwater, brackish] by (2), but not (3). Because these species are primarily freshwater and never marine, we consider this a freshwater stenohaline genus.

*Puntioplites*: **Fresh**. All species freshwater following (2) and (3).

*Pygocentrus*: **Fresh**. All species freshwater following (2) and (3).

*Silurus*: **Fresh**. All species freshwater following (2) and (3). Two species (*S. soldatovi* and *S. triostegus*) are considered [freshwater, brackish] by (2), but not (3). Because these species are primarily freshwater and never marine, we consider this a freshwater stenohaline genus.

*Sternopygus*: **Fresh**. All species freshwater following (2) and (3).

*Synodontis*: **Fresh**. All species freshwater following (2) and (3).

*Tor*: **Fresh**. All species freshwater following (2) and (3).

*Trichomycterus*: **Fresh**. All species freshwater following (2) and (3). *T. pradensis* is considered [freshwater, brackish] by (2), but not (3). Because these species is primarily freshwater and never marine, we consider this a freshwater stenohaline genus.

*Triplophysa*: **Fresh**. All species freshwater following (2) and (3).

*Xenocharax*: **Fresh**. All species freshwater following (2) and (3).

*Zacco*: **Fresh**. All species freshwater following (2) and (3).

### S7. Morphological matrix justification

We herein revise the 63-trait morphological matrix of Liu et al. (12) based on the authors’ interpretations of available fossil material for †*Acronichthys maccagnoi*, †*Chanoides macropoma*, †*Nardonoides chardoni*, and †*Santanichthys diasii*. The authors also code an additional species, the recently-named †*Sooinichthys varii* from the Albian Tlayúa Quarry of Puebla, Mexico (20). This involves the interpretations provided in the literature for each taxon, as well as their comparison of images (photographs and CT scans) illustrating the relevant structures on the fossils. Moreover, in the case of †*Santanichthys diasii*, the reference fossils were examined by one of us (D.M.) and allowed the reinterpretation of some character states (a comprehensive review is currently being drafted).

For each character, following a restatement of its name and number following the codings of (12) and (23), we provide the state for each fossil taxa. When relevant we explain any differences in coding compared with Liu et al. (12), and comment when relevant compared with the descriptions by former authors.

#### Weberian Apparatus: Characters 1–27

See also Fig. SX26–27 for figuring of †*Santanichthys diasii* cranial anatomy.

**Character 1.** Claustrum (0) absent; (1) present [(23) characters 65–67].

Coding for †*Acronichthys maccagnoi*: “?”

Coding for †*Chanoides macropoma* and †*Nardonoides chardoni*: “1” and “1”

Coding for †*Sooinichthys varii*: “1”

Coding for †*Santanichthys diasii*: we agree with Filleul & Maisey (30). There is no trace of a claustrum. There is no trace of foreshortening in the first four centra and moreover no trace of modifications that could configure the presence of Weberian ossicles. Despite this original description, Liu et al. (12), coded this state ambiguously. “0” (and not “?”)

**Character 2.** scaphium (0) absent; (1) present [(23) characters 65-66, 68].

Coding for †*Acronichthys maccagnoi*: the bony piece interpreted as a scaphium in Liu et al. (12) is not in the right position for this interpretation (it is related to the third vertebra, not the first). We do not agree with the reviewer’s suggestion on how to code this character and make them change their initial coding. In the species description, the tripus and scaphium are said to be observed in some fossils (see Supplementary Information of their paper, lines 524–526), with reference to Fig. S5C, which shows a segmentation of the extant *Citharinus* WA, and to Figs. 2A and 2B, which show a segmentation of the anterior vertebral region of specimen 003. In the latter, the position of the bone in question to the third vertebra is evident. We agree with the author’s initial coding: “?” (and not “1”).

Coding for †*Chanoides macropoma* and †*Nardonoides chardoni*: “1” and “0”

Coding for †*Sooinichthys varii*: “1”

Coding for †*Santanichthys diasii*: Filleul & Maisey (30) described a scaphium with a “ventral process that fits within a pit in the anterior border of the first centrum. It also has a shell-like concave anterior edge extending anteriorly beyond the end of the basioccipital, and an ascending process projecting posteriorly and dorsally”. The bone interpreted and figured as such (their Fig. 6) is a dislocated (cichloid) scale, the ventral process is a process of the neural arch 1, and the foramen is a spot on the scale. No exemplar displays what can be interpreted as a scaphium, but some exemplar was damaged in that area so we prefer not to code this character. “?” (and not “1” in Liu et al.).

**Character 3.** Intercalarium (0) absent; (1) present, even if reduced in size [(23) characters 69-71].

Coding for †*Acronichthys maccagnoi*: We found no evidence in the paper of an intercalarium being present. The description of the species does not explicitly state that it is present. Elsewhere, in an osteological comparison, the presence of an intercalarium is told “inferred solely from the presence of an articular pit on the second vertebral centrum” (Supp. l. 777), with reference to S7A&B. However, this structure is not labelled, and we could not identify any indication of an intercalarium articulation. Moreover, the two vertebrae attributed to †*Acronichthys maccagnoi* (one articulated and one not) differ significantly and may not belong to the same fish. “?”

Coding for †*Chanoides macropoma* and †*Nardonoides chardoni*: “1” and “0”

Coding for †*Sooinichthys varii*: “1”

Coding for †*Santanichthys diasii*: Liu et al. coded “1” following the description of Filleul & Maisey (30). According to the original description the “The intercalarium of †*Santanichthys diasii* has a pedicel ventrally that articulates with the second vertebra. The bone is similar to the scaphium in shape, but its anterior edge is pointed”. Filleul and Maisey (30) illustrated the intercalarium (their Fig. 6). The so-called intercalarium described is not associated to any vertebral centra. We interpretate it as a fragment of a skull roof bone fragment, probably a part of a broken dermopterotic. “0” (and not “1”)

**Character 4.** Tripus: (0) absent; (1) present [modified from (23) characters 85–86].

Coding for †*Acronichthys maccagnoi*: Identifying a tripus is problematic since the bone shown in Liu et al. (12, Fig. 2) is reconstructed from two fragments in Geomagic (specimen 003, Liu et al. (12, Fig. 1D and the zoom in 2B); in this reconstruction, the supposed tripus is an impair piece placed between two supposed suspensorium bones (whereas the tripus is a paired piece). If we understand correctly the figures, the interpretation of the holotype specimen (1506) is different but equally problematic: in the photo of the specimen (12, Fig. 1A), we see that the right side is exposed and the left side is in the matrix, yet the CT scan reveals only the right lateral structures (S2) and not those on the left side: we only have supposed right suspensorium and tripus. The absence of the left counterparts is not discussed, even though they are expected. “?” (and not “1”)

Coding for †*Chanoides macropoma* and †*Nardonoides chardoni*: (see character 5) “?” and “?” (and not “1” and “1”)

Coding for †*Sooinichthys varii*: “1”

Coding for †*Santanichthys diasii*: Liu et al. coded “1” following the description of Filleul & Maisey (30). The tripus in (30) is the proximal edge of anterior ribs (their figure 5). There is no trace of tripus in any specimen of †*Santanichthys diasii*. Vertebral centrum 2 (V2) is not preserved, except the associated rib which was identified as a tripus. “0” (and not “1”).

**Character 5.** Tripus: (0 two separate pieces; (1) one single ossicle with the paraphyseal part and transformator part partially fused; (2) fused into one single ossicle [modified from (23) characters 85–86].

Coding for †*Acronichthys maccagnoi*: Non applicable (see character 4). “-” (and not “1”)

Coding for †*Chanoides macropoma* and †*Nardonoides chardoni*: In contrast to Liu et al. (12) understanding of their interpretation, Mayrinck et al. (25) questioned the homology of the bone displaying the shape and location of the tripus. The coherent coding is “?” and “?” (and not “0” and “0”)

Coding for †*Sooinichthys varii*: “1”

Coding for †*Santanichthys diasii*: Since there is no tripus in †*Santanichthys diasii*, the character is not applicable. “-” (and not “0”)

**Character 6.** Parapophysis of first centrum (0) absent; (1) present (even if modified) [autogenous in most primitive teleosts and *Chanos*) or absent on first centrum (cithariniforms) or absent on second centrum (siluriforms). In other otophysans, the parapophysis is present as a lateral process fused to the centrum]. This is considered the same as the lateral process of Cypriniformes which may be present or absent in characiphysans and cypriniphysans [modified from (23) characters 82].

Coding for †*Acronichthys maccagnoi*: “?”

Coding for †*Chanoides macropoma* and †*Nardonoides chardoni*: the descriptions of †*Chanoides macropoma* and †*Nardonoides chardoni* show that no structure can be interpreted as a parapophysis (or transverse process): “0” and “0” (and not “1” and “?”)

Coding for †*Sooinichthys varii*: “?” [There is a small protruding structure visible on V1 (20), but its interpretation cannot be determined]

Coding for †*Santanichthys diasii*: Liu et al. coded “?” since Filleul & Maisey (30) did not addressed any comment concerning the parapophysis on the first centrum. In all exemplars observed with the first centrum preserved, it presents a ventral projection associated to a rib – clearly a parapophysis. “1” and not “?”

**Character 7.** Parapophysis of second centrum: (0) absent; (1) present and rod-like; (2) present and modified to form the lateral process (= transverse process).

Coding for †*Acronichthys maccagnoi*: “?”

Coding for †*Chanoides macropoma* and †*Nardonoides chardoni*: 2 and “0” (and not 2 and “?”)

Coding for †*Sooinichthys varii*: Like for character 6/ V1, there is a small protruding structure visible on V2, but its interpretation cannot be determined. “?”

Coding for †*Santanichthys diasii*: The same as for character 6. Liu et al. coded “?”. †*Santanichthys diasii* presents ribs associated to the first four centra. Photo below. Please, it is very important to highlight that all exemplars are preserved in a mass mortality and the largest specimen is at most 15 mm. Please, pay attention to the rib associated to a ventral projection in the ventral surface of the second centrum. “1” (and not “?”)

**Character 8 and Character 9.** Character 8: parapophysis and rib of fourth centrum: (0 rib elongate, not fused to parapophysis and having similar morphology to those of the following centra; (1) present but greatly shortened or modified [(23) characters 89-90]. Character 9: shortened rib of fourth centrum: (0 small size, no contribution from rib 2; (1) extends vertically, fused with rib 2 and forms a rib complex; (2) extends laterally, fused with ribs, and parapophysis of third or second centrum [modified from (23) characters 89–91].

Coding for †*Acronichthys maccagnoi*: A large and stout rib is present in †*Acronichthys maccagnoi*, as evidenced by the holotype specimen and fossil description (Fig. S2). This rib may articulate with the short parapophysis, which appears to develop antero-laterally on the fourth centrum (Liu et al. (12): S5A). “?” and “-” (and not “1” and “1” for characters 8 and 9 respectively)

Coding for †*Chanoides macropoma* and †*Nardonoides chardoni*: “1” and “-” for character 8 (and not “1” and ); “?” and “-” for character 9 (and not “0” and )

Coding for †*Sooinichthys varii*: “1” and “1”

Coding for †*Santanichthys diasii*: Ribs are articulates with the fourth centrum and the state “0” of character 8 is easily identified. There is no described state in character 9 that applies to †*Santanichthys diasii*. “0” and “-” (and not “?” & “?”).

**Character 10.** Os suspensorium (shortened rib of C4 fused with parapophysis and bearing a median process): (0) absent; (1) present [(23) character 88].

Coding for †*Acronichthys maccagnoi*: the presence of a suspensory bone is not proved (paired and dependent on V4, see also character 8 and 9) and a pair of true rib articulates on V4. “?” (and not “1”)

Coding for †*Chanoides macropoma* and †*Nardonoides chardoni*: “1” and “0”

Coding for †*Sooinichthys varii*: It is present but not in a fused configuration. “0”

Coding for †*Santanichthys diasii*: “0” (and not “?”)

**Character 11.** Left and right inner arm of os suspensoria (when present): (0 meeting in midline; (1) not meeting in midline. The left and right os suspensoria of catfish do not meet in the midline [modified from (23) character 88].

Coding for †*Acronichthys maccagnoi*: see character 10. “-” (and not “0”).

Coding for †*Chanoides macropoma* and †*Nardonoides chardoni*: “0” and “?”

Coding for †*Sooinichthys varii*: “-”

Coding for †*Santanichthys diasii*: “-” (and not “?”)

**Character 12.** “Transverse process” of the fourth centrum: (0 not broadly expanded in the horizontal plane; (1) broadly expanded in the horizontal plane and fused to complex centrum [(23) characters 89-91].

Coding for †*Acronichthys maccagnoi*: not applicable. “-” (and not “0”)

Coding for †*Chanoides macropoma* and †*Nardonoides chardoni*: “-” and “-” (and not “0” and “0”)

Coding for †*Sooinichthys varii*: “-”

Coding for †*Santanichthys diasii*: “-” (and not “0”)

**Character 13.** Length of first centrum compared to fifth centrum: (0 first centrum of similar length to fifth centrum; (1) shorter than fifth centrum [modified from (23) characters 78-79].

Coding for †*Acronichthys maccagnoi*: “1”.

Coding for †*Chanoides macropoma* and †*Nardonoides chardoni*: “0” and “0”

Coding for †*Sooinichthys varii*: The first centrum is slightly shortened at its base, but not significantly. “0”

Coding for †*Santanichthys diasii*: All centra are equal in size. “0” (and not “1”)

**Character 14.** Second centrum: (0 same length or longer than first centrum; (1) shorter in length than first centrum [modified from (23) characters 78-79].

Coding for †*Acronichthys maccagnoi*: All the figures show that the dimensions of centra 1 and 2 are pretty similar, including in their length. “0” (and not “1”)

Coding for †*Chanoides macropoma* and †*Nardonoides chardoni*: “0” and “0”

Coding for †*Sooinichthys varii*: “0”

Coding for †*Santanichthys diasii*: All centra are equal in size. There is any pattern of foreshorten in the first occipital centra in †*Santanichthys diasii*. “0” (and not “1”)

**Character 15.** Third centrum: (0 same length or longer than fifth centrum; (1) shorter in length than fifth centrum [modified from (23) characters 78–79].

Coding for †*Acronichthys maccagnoi*: “0”

Coding for †*Chanoides macropoma* and †*Nardonoides chardoni*: “1” and “1”

Coding for †*Sooinichthys varii*: “0”

Coding for †*Santanichthys diasii*: “0”

**Character 16.** Fourth centrum: (0 same length or longer than fifth centrum; (1) shorter in length than fifth centrum [modified from (23) characters 78-79].

Coding for †*Acronichthys maccagnoi*: “0”

Coding for †*Chanoides macropoma* and †*Nardonoides chardoni*: “0” and “0”

Coding for †*Sooinichthys varii*: “0”

Coding for †*Santanichthys diasii*: “0”

**Character 17.** Second and third centra: (0 separate from one another; (1) fused together [Fink and Fink (27:18) discussed this feature].

Coding for †*Acronichthys maccagnoi*: “0”

Coding for †*Chanoides macropoma* and †*Nardonoides chardoni*: “0” and “0”

Coding for †*Sooinichthys varii*: “0”

Coding for †*Santanichthys diasii*: “0”

**Character 18.** Third and fourth centra: (0 separate from one another; (1) exhibiting fusion of the elements but retaining clear distinctions between the centra such that each centrum is clearly identifiable; (2) fused together into a solid mass such that the individual centra are not delimited [modified from (23) character 80].

Coding for †*Acronichthys maccagnoi*: “0”.

Coding for †*Chanoides macropoma* and †*Nardonoides chardoni*: “0” and “0”

Coding for †*Sooinichthys varii*: “0”

Coding for †*Santanichthys diasii*: “0”

**Character 19.** Fifth centrum (0 free from anterior centra or Weberian apparatus; (1) incorporated into the Weberian apparatus [Fink and Fink (27: p. 308) discussed this feature].

Coding for †*Acronichthys maccagnoi*: “0”.

Coding for †*Chanoides macropoma* and †*Nardonoides chardoni*: “0” and “0”

Coding for †*Sooinichthys varii*: “0”

Coding for †*Santanichthys diasii*: “0”

**Character 20.** Supraneural anterior to neural arch of first vertebra (0 present; (1) absent [(23) characters 58].

Coding for †*Acronichthys maccagnoi*: “1”

Coding for †*Chanoides macropoma* and †*Nardonoides chardoni*: “1” and “0”

Coding for †*Sooinichthys varii*: “1”

Coding for †*Santanichthys diasii*: “1”

**Character 21.** Supraneural anterior to neural arch of second vertebra (0 present; (1) absent. [(23) characters 59 and 61].

Coding for †*Acronichthys maccagnoi*: “1”

Coding for †*Chanoides macropoma* and †*Nardonoides chardoni*: “0” and “0”

Coding for †*Sooinichthys varii*: “0”

Coding for †*Santanichthys diasii*: “1”

**Character 22.** Anterior supraneural(s) expanded ventrally to form a synchondral joint with the neural arches of the third and fourth vertebrae (the presence of a neural complex): (0 not expanded, similar to other supraneurals; (1) expanded, but do not form joint with neural arches; (2) expanded, and form neural complex [modified from (23) character 60 based on the ontogeny and development study by Hoffman and Britz (124)].

Coding for †*Acronichthys maccagnoi*: From what can be seen on the holotype (Liu et al. (12): fig. 2A&B, S2A&B), the supposed neural complex appears to occupy the space where the neural arch and spine of the third centrum would be. We find no evidence to support the ‘supraneural complex’ interpretation; the presence of such a complex remains to be demonstrated. “?” (and not “2”)

Coding for †*Chanoides macropoma* and †*Nardonoides chardoni*: “2” and “2” (and not “2” and “1”)

Coding for †*Sooinichthys varii*: “2”

Coding for †*Santanichthys diasii*: “2”

**Character 23.** Supraneurals contributing to neural complex: (0 supraneural 2 and 3; (1) supraneural 3 only; (2) no supraneural contributes to the neural complex [modified from (23) character 61]

Coding for †*Acronichthys maccagnoi*: “?”

Coding for †*Chanoides macropoma* and †*Nardonoides chardoni*: “0” and “0” (and not “-” and “-”)

Coding for †*Sooinichthys varii*: “0”

Coding for †*Santanichthys diasii*: †*Santanichthys diasii* and †*Sooinichthys varii* share the same condition for supraneural 2 and 3. “0” (and not “?”).

**Character 24.** Articulation between anterior supraneural and posterior margin of cranium (posterior process of supraoccipital and suture lines between supraoccipital and exoccipital): (0 absent, the anterior supraneural does not reach the cranium; (1) anterior supraneural expands close to the cranium, but does not suture with it; (2) strong and extensive articulation present [(23) character 62].

Coding for †*Acronichthys maccagnoi*. “?”

Coding for †*Chanoides macropoma* and †*Nardonoides chardoni*: “0” and “0”

Coding for †*Sooinichthys varii*: “1”

Coding for †*Santanichthys diasii*: There is an expanded supraneural 2, but it is not possible to ascertain any relationship with the cranium. “0” (Liu et al. coded “0”).

**Character 25.** Anterior neural arches dorsomedially expanded to about one another and the posterior margin of the exoccipital to form a roof over the neural canal: (0) absent; (1) present. [(23) character 63].

Coding for †*Acronichthys maccagnoi*: due to the poor preservation, we found no way to estimate this character state in †*Acronichthys maccagnoi* from the information available in Liu et al. (12). “?” (and not “1”)

Coding for †*Chanoides macropoma* and †*Nardonoides chardoni*: “1” and “1”

Coding for †*Sooinichthys varii*: “1”

Coding for †*Santanichthys diasii*: “1”

**Character 26.** Anterior neural arch greatly enlarged with extensive, tight joint with exoccipitals: (0 absent, only a restricted articulation mediated through cartilage exists between exoccipitals and anterior neural arch; (1) present [(23) character 65].

Coding for †*Acronichthys maccagnoi*: “?”

Coding for †*Chanoides macropoma* and †*Nardonoides chardoni*: “0” and “0”

Coding for †*Sooinichthys varii*: “0”

Coding for †*Santanichthys diasii*: “0”

**Character 27.** Neural arches of third and fourth centra: (0 separate from one another, may or may not be fused to centra; (1) fused to one another and to the complex centrum [(23) character 76].

Coding for †*Acronichthys maccagnoi*: “0”

Coding for †*Chanoides macropoma* and †*Nardonoides chardoni*: “0” and “0”

Coding for †*Sooinichthys varii*: “0”

Coding for †*Santanichthys diasii*: “0”

#### Caudal Fin: Traits 28–41

See also Fig. SX25 for figuring of †*Santanichthys diasii* caudal anatomy.

**Character 28.** Compound centrum: (0 absent, first preural and first ural centra are separate elements; (1) present, the first preural and first (or first and second) ural centra are fused into a single unit [modified from (23) character 110].

Coding for †*Acronichthys maccagnoi*: “1”

Coding for †*Chanoides macropoma* and †*Nardonoides chardoni*: “1” and “1”

Coding for †*Sooinichthys varii*: “1” Coding for †*Santanichthys diasii*: “1”

**Character 29.** Compound centrum (fusion of two elements usually identified as the first preural and first ural centra, but suggested by some authors to also include u2): (0 separate from uroneurals; (1) fused with uroneural to form a urostyle [modified from (23) character 110].

Coding for †*Acronichthys maccagnoi*: For this character only, we were not able to agree with each other on a same interpretation of the iconography. We conclude that the data available do not allow to conclude: “?” (and not “1”).

Coding for †*Chanoides macropoma* and †*Nardonoides chardoni*: “1” and “1”

Coding for †*Sooinichthys varii*: “1”

Coding for †*Santanichthys diasii*: “1”

**Character 30.** Neural spine on first preural centrum: (0 present; (1) absent. Coding for †*Acronichthys maccagnoi*: “1”

Coding for †*Chanoides macropoma* and †*Nardonoides chardoni*: Despite the correct reference and citation of the character state ‘neural spine present’ from Mayrinck et al. (25), the character state was coded as ‘absent’ in the corrected matrix published by Liu et al. (12) “0” and “0” (and not “1” and “1”)

Coding for †*Sooinichthys varii*: “1”

Coding for †*Santanichthys diasii*: “0” (and not “1”; see also character below)

**Character 31.** Haemal arch (of parhypural) on compound centrum or 1st preural centrum if compound centrum not formed: (0 autogenous; (1) arch fused with centrum [modified from (23) character 112].

Coding for †*Acronichthys maccagnoi*: “?”

Coding for †*Chanoides macropoma* and †*Nardonoides chardoni*: “1” and “0” (and not “0” and “0”)

Coding for †*Sooinichthys varii*: “1”

Coding for †*Santanichthys diasii*: “1” (see also Fig. SX26–27, displays an interpretated photograph of the interpretative drawing displayed in (30: Fig. 9).

**Character 32.** Haemal arch on second preural centrum: (0 autogenous; (1) arch fused with centrum [modified from (23) character 112].

Coding for †*Acronichthys maccagnoi*: The resolution is sufficient in the iconography provided by Liu et al. (12) to cast doubt on this fusion since it really seems a separate element on their 3D model (their Fig. 1F, G). “?” (and not “1”).

Coding for †*Chanoides macropoma* and †*Nardonoides chardoni*: “0” and “0”

Coding for †*Sooinichthys varii*: “1”

Coding for †*Santanichthys diasii*: “1”

**Character 33.** Haemal arch on third (or more anterior) preural centrum: (0 autogenous; (1) arch fused with centrum [(23) character 111].

Coding for †*Acronichthys maccagnoi*: For us, the resolution is sufficient in the iconography to cast doubt on this fusion; it really seems a separate element on the 3D model (1F, G). “?” (and not “1”).

Coding for †*Chanoides macropoma* and †*Nardonoides chardoni*: “0” and1 (and not “1” and “1”)

Coding for †*Sooinichthys varii*: “0”

Coding for †*Santanichthys diasii*: “1”

**Character 34.** Epural number: (0 three or more; (1) two; (2) one; (3) none [modified from (23) character 115].

Coding for †*Acronichthys maccagnoi*: The interpretation is questioned given the poor state of preservation of the distal part of the inner caudal skeleton. Moreover some of the supposed caudal ray fragments may correspond to fragments of epural bones (Liu et al. (12): fig. 1F, G). “?”

Coding for †*Chanoides macropoma* and †*Nardonoides chardoni*: “0” and “0”

Coding for †*Sooinichthys varii*: “0”

Coding for †*Santanichthys diasii*: “?”

**Character 35.** Uroneural number: (0 a urostyle may form through fusion with centrum, but no separate uroneural; (1) two uroneurals (second one remains free); (2) three or more uroneurals.

Coding for †*Acronichthys maccagnoi*: “?”

Coding for †*Chanoides macropoma* and †*Nardonoides chardoni*: “2” and “2”

Coding for †*Sooinichthys varii*: “2”

Coding for †*Santanichthys diasii*: according to our interpretation there are at least 3 uroneurals (see figure above). “2” (and not “?”)

**Character 36.** Parhypural and hypural 1: (0 contact or fuse to terminal centrum separately; (1) phy and hy1 articulate with each other proximally and then suture to or articulate with the terminal centrum.

Coding for †*Acronichthys maccagnoi*: “0”.

Coding for †*Chanoides macropoma* and †*Nardonoides chardoni*: “0” and “0”

Coding for †*Sooinichthys varii*: “0”

Coding for †*Santanichthys diasii*: “0”

**Character 37.** Hypural 1 and 2 fusion: (0 no fusion between hypurals one and two; (1) hypurals one and two partially fused distally or fully fused together.

Coding for †*Acronichthys maccagnoi*: It is not possible to ascertain such degree of fusion between the hypural 1 and 2 of even if it exists (Liu et al. (12): fig. 1F, G). “?”

Coding for †*Chanoides macropoma* and †*Nardonoides chardoni*: “0” and “0”

Coding for †*Sooinichthys varii*: “0”

Coding for †*Santanichthys diasii*: “0”

**Character 38.** First hypural: (0 autogenous, even if it meets the compound centrum the two are not fused; (1) fused to compound centrum [modified from (23) characters 112, and 113].

Coding for †*Acronichthys maccagnoi*: “0”.

Coding for †*Chanoides macropoma* and †*Nardonoides chardoni*: “0” and “0”

Coding for †*Sooinichthys varii*: “1”

Coding for †*Santanichthys diasii*: the review of the material allows us to distinguish and autogenous first hypural (see figure above). “0” (and not “1”).

**Character 39.** Second hypural: (0 autogenous, even if it meets the compound centrum the two are not fused; (1) fused to compound centrum [(23) character 114].

Coding for †*Acronichthys maccagnoi*: “0”.

Coding for †*Chanoides macropoma* and †*Nardonoides chardoni*: “0” and “0” (and not “1” and “0”)

Coding for †*Sooinichthys varii*: “1”

Coding for †*Santanichthys diasii*: the review of the material allows us to distinguish and autogenous second hypural (see figure above). “0” (and not “1”)

**Character 40.** Hypurapophysis: (0) absent; (1) present.

Coding for †*Acronichthys maccagnoi*: “?”

Coding for †*Chanoides macropoma* and †*Nardonoides chardoni*: “0” and “0”

Coding for †*Sooinichthys varii*: “0”

Coding for †*Santanichthys diasii*: The proximal edge of the haemal elements is preserved and there are no hypurapophysis (see figure above). “0” (and not “?”).

**Character 41.** Caudal fin ray formula: (0 i,9,8,i or more; (1) i,9,9, i [most catfish with many reductions]; (2) i,8,8, i, or fewer [modified from (23) character 108].

Coding for †*Acronichthys maccagnoi*: “?”

Coding for †*Chanoides macropoma* and †*Nardonoides chardoni*: “0” and “0” (and not “?” and “?”)

Coding for †*Sooinichthys varii*: “0”

Coding for †*Santanichthys diasii*: No caudal fin is completely preserved, making it impossible to accurately infer the formula. “?”

#### Jaws: 42–49

See also Fig. SX26–27 for figuring of †*Santanichthys diasii* cranial anatomy.

**Character 42.** Dorsomedial (= ascending) process of premaxilla: (0 lateral rather than along midline (left and right do not meet in midline); (1) along midline (left and right meet in midline); (2) absent [modified from Fink & Fink (23) character 37].

Coding for †*Acronichthys maccagnoi*: Despite the low resolution it is possible to observe the anterior outline of the premaxillary preserved. “2”

Coding for †*Chanoides macropoma* and †*Nardonoides chardoni*: “1” and “2”

Coding for †*Sooinichthys varii*: “2”

Coding for †*Santanichthys diasii*: “2”

**Character 43.** Premaxillary teeth: (0 present; (1) absent [modified from (23) character].

Coding for †*Acronichthys maccagnoi*: “?”

Coding for †*Chanoides macropoma* and †*Nardonoides chardoni*: “1” and “1”

Coding for †*Sooinichthys varii*: “1”

Coding for †*Santanichthys diasii*: “1”

**Character 44.** Maxilla: (0 forms part of jaw margin; (1) excluded from jaw margin [Fink and Fink (27: p. 308) noted the form of the maxilla in catfishes compared to the presumed primitive form].

Coding for †*Acronichthys maccagnoi*: As Liu et al. (12) identify it as ‘tentatively maxilla’, it seems complicated to code this character state; from the iconography, we are also unable to recognise the bones of the upper jaw with certainty: “?” (and not “0”)

Coding for †*Chanoides macropoma* and †*Nardonoides chardoni*: “0” and “0”

Coding for †*Sooinichthys varii*: “0”

Coding for †*Santanichthys diasii*: “0”

**Character 45.** Dentary coronoid process: (0 prominent, clearly higher than symphysis, 2.5 times or more the height of the symphysis; (1) low, similar in height or not much higher than symphysis.

Coding for †*Acronichthys maccagnoi*: The dentary is said to be one of the best-preserved jaw bones, and it appears as a low bar bone with no sign of a coronoid process. However, we cannot distinguish if the dorsal edge is complete or not: we agree with Liu et al.’ caution. “?”

Coding for †*Chanoides macropoma* and †*Nardonoides chardoni*: “0” and “0”

Coding for †*Sooinichthys varii*: “1”

Coding for †*Santanichthys diasii*: “0”

**Character 46.** Dentary symphysis: (0 smooth, no sutures between left and right elements; (1) interdigitating sutures uniting left and right dentaries.

Coding for †*Acronichthys maccagnoi*: The quality of preservation seems insufficient to us (anterior end of the dentary unknown) to code this character. “?” (and not “0”).

Coding for †*Chanoides macropoma* and †*Nardonoides chardoni*: “?” and “?” (and not “0” and “0”)

Coding for †*Sooinichthys varii*: “0”

Coding for †*Santanichthys diasii*: No specimen is well preserved enough to allow any inference to be made about the symphysis. “?” (and not “0”)

**Character 47.** Dentary teeth: (0 present; (1) absent [modified from (23) character 42].

Coding for †*Acronichthys maccagnoi*: “?”

Coding for †*Chanoides macropoma* and †*Nardonoides chardoni*: “1” and “1”

Coding for †*Sooinichthys varii*: “0”

Coding for †*Santanichthys diasii*: “1”

**Character 48.** Anguloarticular and retroarticular: (0 separate elements; (1) retroarticular fused to anguloarticular.

Coding for †*Acronichthys maccagnoi*: According to Liu et al.: fig. 1H, part of the purple bone (labelled RR) (posterior-ventral) may be a retroarticular bone, as it is in the correct position and has the correct shape. However, according to the label, it seems that all purple bone is interpreted as RR. In any case, Liu et al. (12) do not discuss their interpretation of the highly fragmentary remains of the lower jaw which cast doubt on character state coding: “?”

Coding for †*Chanoides macropoma* and †*Nardonoides chardoni*: “0” and “0”

Coding for †*Sooinichthys varii*: “0”

Coding for †*Santanichthys diasii*: We were able to observe the character in one specimen that clearly displays state “0”. “0” (and not “?”).

**Character 49.** Kinethmoid: (0) absent; (1) present [(23) character 1].

Coding for †*Acronichthys maccagnoi*: The preservation of the ethmoid region is insufficient for assessing this character trait. Furthermore, the article does not discuss its presence or absence: “?” (and not “0”).

Coding for †*Chanoides macropoma* and †*Nardonoides chardoni*: “1” and “1”

Coding for †*Sooinichthys varii*: “0”

Coding for †*Santanichthys diasii*: “0”

#### Other characters: 50–63

**Character 50.** Basisphenoid: (0 present; (1) absent [(23) character 7].

Coding for †*Acronichthys maccagnoi*: “?”

Coding for †*Chanoides macropoma* and †*Nardonoides chardoni*: “1” and “1” (and not “?” and “?”)

Coding for †*Sooinichthys varii*: “1”

Coding for †*Santanichthys diasii*: “1” (and not “?”).

**Character 51.** Pharyngobranchial toothplate: (0 more than one toothplate present; (1) one toothplate; (2) absent [modified from (23) characters 47, 48, 50, and 51].

Coding for †*Acronichthys maccagnoi*: The diagnosis indicates ‘*more than one pair of pharyngeal bones moderately developed, bearing minute dentition’*. However, the description indicates “*The fifth ceratobranchial (Fig 1D,H) is slender and triangular; the left and right pair is preserved in TMP 2019.022.0003 (Fig 1D,H). They appear to bear dentition, possibly a single row of minute teeth. An additional pharyngeal bone next to the fifth ceratobranchial in this specimen is probably one of the infrabranchials*”. It is unfortunate that the teeth are not shown, but above all, the coding of the character seems impossible to us given the very limited knowledge of pharyngeal bones in †*Acronichthys maccagnoi*. “?” (and not “0”).

Coding for †*Chanoides macropoma* and †*Nardonoides chardoni*: “?” and “?”

Coding for †*Sooinichthys varii*: “?”

Coding for †*Santanichthys diasii*: “?”

**Character 52.** Pharyngeal teeth on fifth ceratobranchial: (0) absent; (1) denticles or minute teeth present; (2) teeth present with some or all enlarged, ankylosed to pharyngeal bone [modified from (23) characters 49, 52, and 53].

Coding for †*Acronichthys maccagnoi*: The presence is not shown (the character state cannot be discussed) but teeth on Cb5 are explicitly described in Liu et al. (12). “1”

Coding for †*Chanoides macropoma* and †*Nardonoides chardoni*: “1” and “1” (and not “2” and “?”)

Coding for †*Sooinichthys varii*: In our description “The branchial arches are exposed partially, but a portion of the right opercle is broken in IGM 14351 (20: fig. 4), showing the five elongated ceratobranchials and the triangular tooth plate of the fifth ceratobranchial. No teeth are visible on any of these bones”. “0”

Coding for †*Santanichthys diasii*: Most exemplars present the branchials elements preserved with a single row of minute teeth. “1” (and not “?”)

**Character 53.** Parietals: (0 two present forming at least one quarter of the skull roof (or 1/3 length of frontal); (1) two present, reduced in size; (2) fused together to form a single element co-ossified with the supraoccipital [modified from (23) characters 10 and 11].

Coding for †*Acronichthys maccagnoi*: “0”.

Coding for †*Chanoides macropoma* and †*Nardonoides chardoni*: “1” and “0”

Coding for †*Sooinichthys varii*: “1”

Coding for †*Santanichthys diasii*: “1”

**Character 54.** Intercalar: (0 present; (1) absent [(23) character 13].

Coding for †*Acronichthys maccagnoi*: “?”

Coding for †*Chanoides macropoma* and †*Nardonoides chardoni*: “?” and “?”

Coding for †*Sooinichthys varii*: “?”

Coding for †*Santanichthys diasii*: “?”

**Character 55.** Supraorbital bone: (0 present; (1) absent [(23) character 19].

Coding for †*Acronichthys maccagnoi*: Interpreting the bone fragment (S2) as a supraorbital bone is not well supported anatomically whenever its presence is probable. We suggest not asserting its presence on this weak base. “?” (and not “0”).

Coding for †*Chanoides macropoma* and †*Nardonoides chardoni*: The presence of a supraorbital bone is noted by Mayrinck et al. (2015). As with †*Acronichthys maccagnoi*, this is also highly probable, but the anatomical evidence is weak. So we agree with Liu et al. (12) and code “?” and “?”

Coding for †*Sooinichthys varii*: “0”

Coding for †*Santanichthys diasii*: “0”

**Character 56.** Metapterygoid-quadrate fenestra: (0 present; (1) absent [modified from (23) character 30].

Coding for †*Acronichthys maccagnoi*: This feature is not described in detail, and we find it very difficult to confirm the presence of this window when the contours of the metapterygoid are badly damaged/partially visible, as far as can be judged from the virtual reconstruction. “?” (and not “0”)

Coding for †*Chanoides macropoma* and †*Nardonoides chardoni*: “1” and “1”

Coding for †*Sooinichthys varii*: “1”

Coding for †*Santanichthys diasii*: There is no metapterygoid-quadrate fenestra. This character is directly related to the shape of the metapterygoid since generally in Otophysi the metapterygoid is X-shaped and the concave posterior tip of the bone creates a fenestra with the ventral edge of the hyomandibula [see Fink & Fink, (23, 27)]. In †*Santanichthys diasii* (see Figure SX16), there is no fenestra between these two bones.

**Character 57.** Symplectic and the postero-ventral portion of quadrate: (0 both present and separate; (1) absent or fused indistinguishably with postero-ventral portion reduced [(23) character 33].

Coding for †*Acronichthys maccagnoi*: “0”.

Coding for †*Chanoides macropoma* and †*Nardonoides chardoni*: “0” and “0”

Coding for †*Sooinichthys varii*: “0”

Coding for †*Santanichthys diasii*: “0”

**Character 58.** Preopercle and interopercle: (0 present, interopercle lies under horizontal/ventral limb of preopercle; (1) shortened antero-posteriorly such that the preopercle lacks a horizontal/ventral limb and the interopercle is a short triangular bone [modified from (23) character 34].

Coding for †*Acronichthys maccagnoi*: “0”.

Coding for †*Chanoides macropoma* and †*Nardonoides chardoni*: “0” and “0”

Coding for †*Sooinichthys varii*: “0”

Coding for †*Santanichthys diasii*: “0”

**Character 59.** Subopercle: (0 present; (1) absent [(23) character 35].

Coding for †*Acronichthys maccagnoi*: “0”

Coding for †*Chanoides macropoma* and †*Nardonoides chardoni*: “0” and “?”

Coding for †*Sooinichthys varii*: “0”

Coding for †*Santanichthys diasii*: “0”

**Character 60.** Opercle: (0 approximately rectangular or oval forming a large lateral covering over the branchial chamber; (1) triangular and reduced in size [(23) character 36].

Coding for †*Acronichthys maccagnoi*: OK: “0”.

Coding for †*Chanoides macropoma* and †*Nardonoides chardoni*: “0” and “?”

Coding for †*Sooinichthys varii*: “0”

Coding for †*Santanichthys diasii*: “0”

**Character 61.** Cranial fontanelle: (0 one or two present; (1) absent.

Coding for †*Acronichthys maccagnoi*: “0”.

Coding for †*Chanoides macropoma* and †*Nardonoides chardoni*: “1” and “1”

Coding for †*Sooinichthys varii*: “0”

Coding for †*Santanichthys diasii*: “0”

**Character 62.** Large and globular lagenar chambers that project well laterally to the cranial condyle: (0) absent; (1) present [modified from (23) character 15].

Coding for †*Acronichthys maccagnoi*: “?”

Coding for †*Chanoides macropoma* and †*Nardonoides chardoni*: “?” and “?”

Coding for †*Sooinichthys varii*: “?”

Coding for †*Santanichthys diasii*: “1”

**Character 63.** Dorsal fin: (0 present; (1) absent [(23) character 104].

Coding for †*Acronichthys maccagnoi*: “0”.

Coding for †*Chanoides macropoma* and †*Nardonoides chardoni*: “0” and “?”

Coding for †*Sooinichthys varii*: “0”

Coding for †*Santanichthys diasii*: “0”

**Fig SX01.**
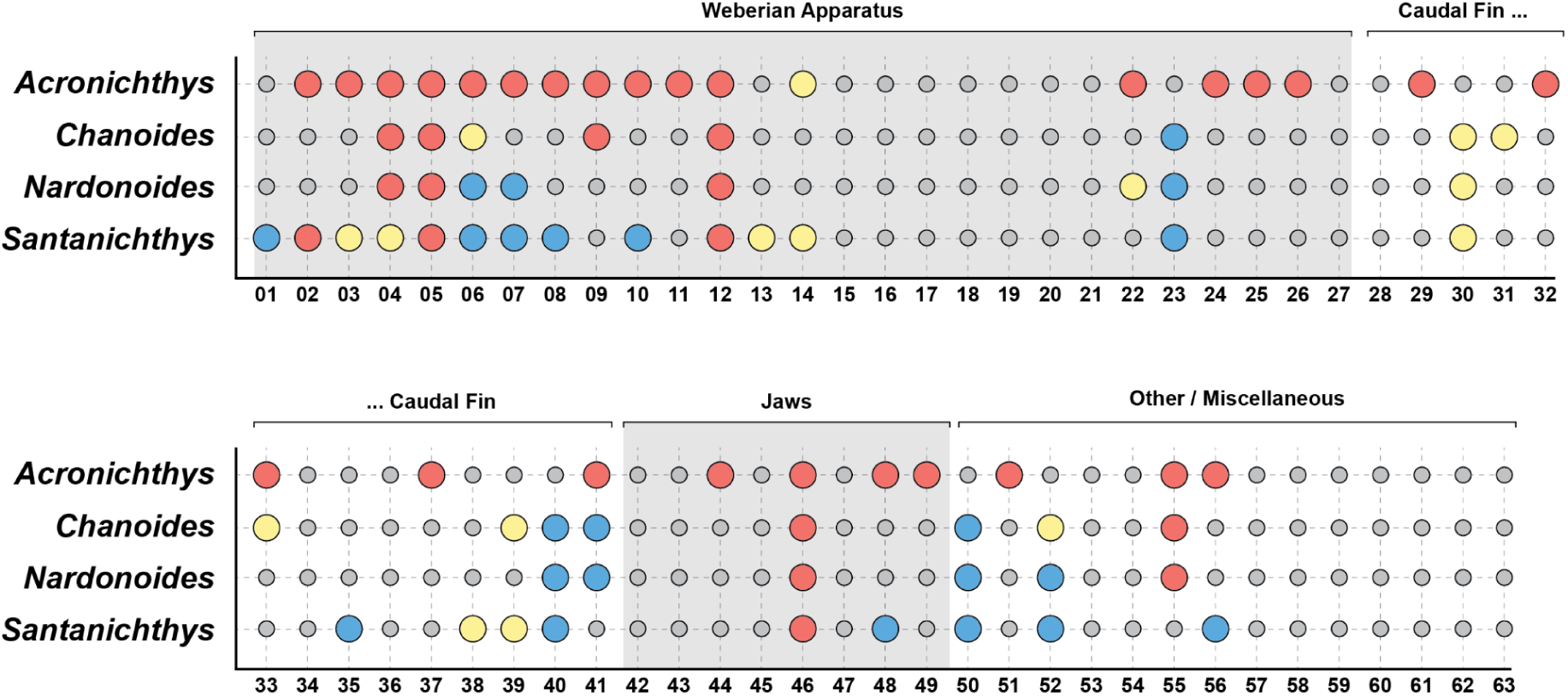
Revisions made to the Liu et al. morphological matrix. Small gray circles indicate no change was made (or changes between “-” and “?”). Red circles indicate previously-coded characters reconsidered to be ambiguous or inapplicable (n = 42). Blue circles indicate previously-ambiguous or inapplicable characters that are coded here (n = 23). Yellow circles indicate characters that were reassigned (i.e., 0 ⇄ 1; n = 16).

**Fig SX02.**
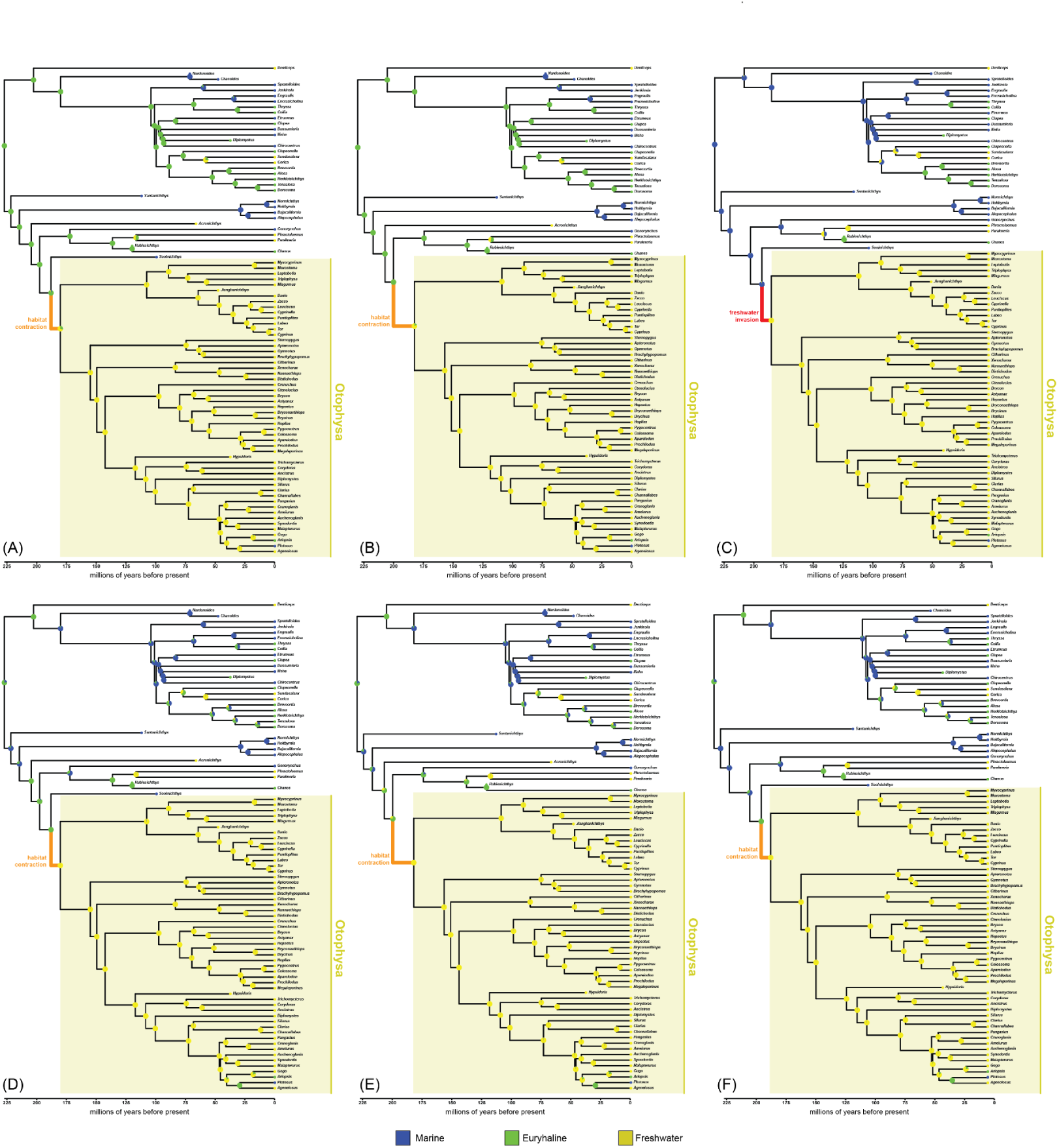
BioGeoBEARS results using best supported topology, rooted on Clupeiformes, with †*Santanichthys* coded as marine, and with different combinations of fossil sampling and model. (**A**) All fossil taxa included, reconstruction under the best fitting model (BAYAREALIKE+J). (**B**) †*Sooinichthys* excluded, reconstruction under the best fitting model (BAYAREALIKE+J). (**C**) †*Acronichthys* and †*Nardonoides* excluded, reconstruction under the best fitting model (BAYAREALIKE+J). (**D**) All fossil taxa included, reconstruction under Liu et al.’s chosen model (DEC). (**E**) †*Sooinichthys* excluded, reconstruction under Liu et al.’s chosen model (DEC). (**F**) †*Acronichthys* and †*Nardonoides* excluded, reconstruction under Liu et al.’s chosen model (DEC).

**Fig SX03.**
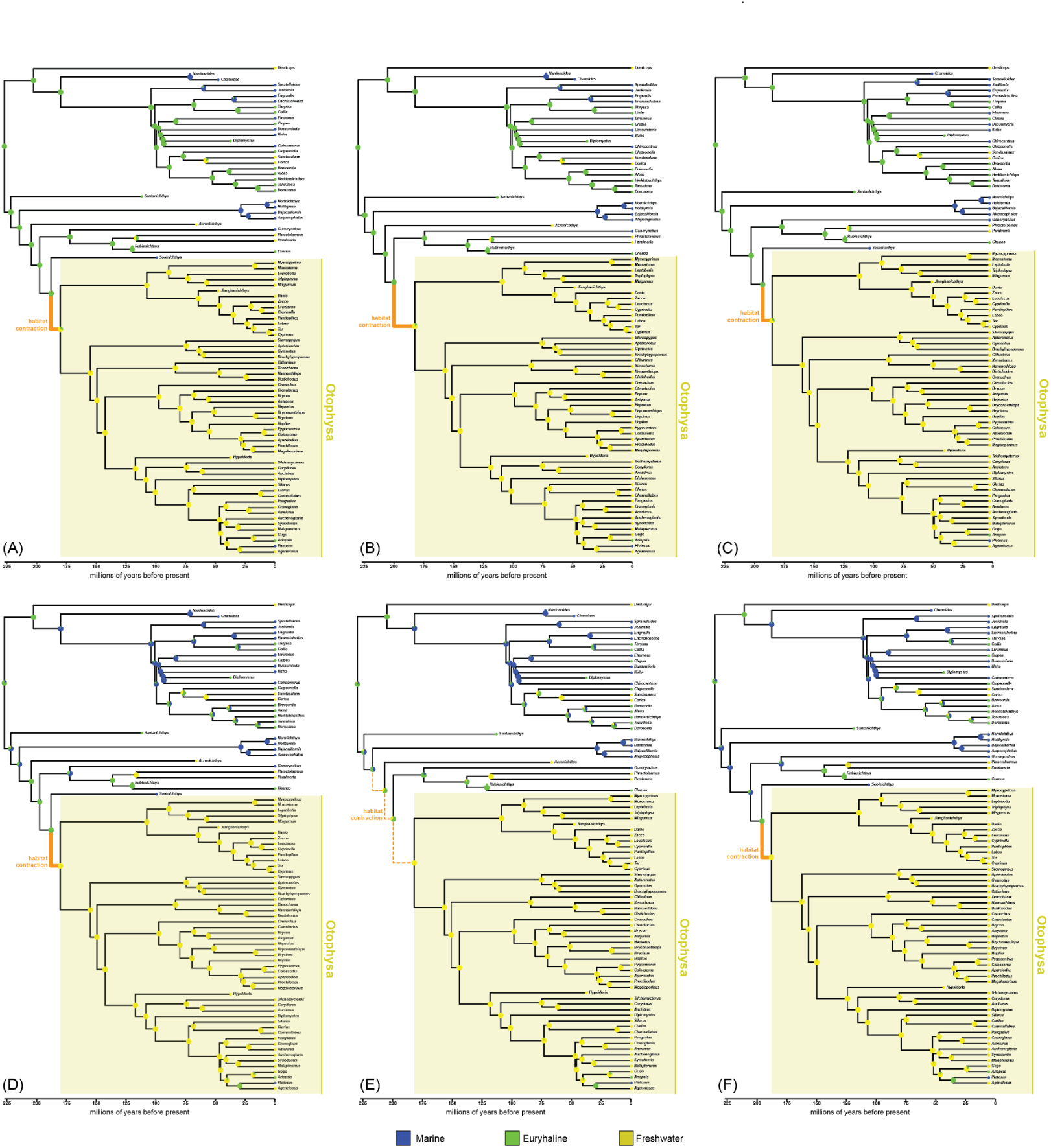
BioGeoBEARS results using best supported topology, rooted on Clupeiformes, with †*Santanichthys* coded as euryhaline, and with different combinations of fossil sampling and model. (**A**) All fossil taxa included, reconstruction under the best fitting model (BAYAREALIKE+J). (**B**) †*Sooinichthys* excluded, reconstruction under the best fitting model (BAYAREALIKE+J). (**C**) †*Acronichthys* and †*Nardonoides* excluded, reconstruction under the best fitting model (BAYAREALIKE+J). (**D**) All fossil taxa included, reconstruction under Liu et al.’s chosen model (DEC). (**E**) †*Sooinichthys* excluded, reconstruction under Liu et al.’s chosen model (DEC). (**F**) †*Acronichthys* and †*Nardonoides* excluded, reconstruction under Liu et al.’s chosen model (DEC).

**Fig SX04.**
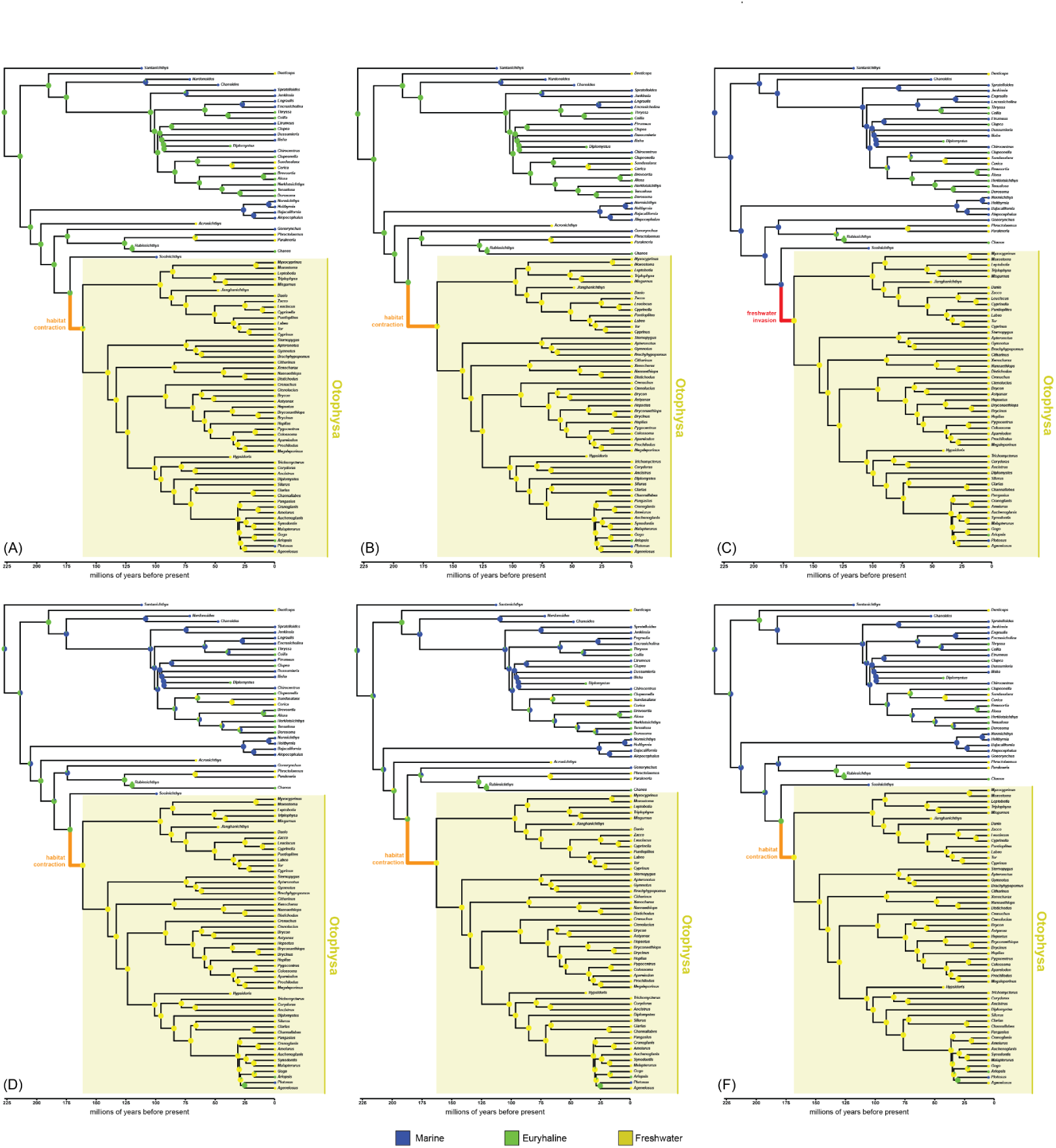
BioGeoBEARS results using best supported topology, rooted on †*Santanichthys*, with †*Santanichthys* coded as marine, and with different combinations of fossil sampling and model. (**A**) All fossil taxa included, reconstruction under the best fitting model (BAYAREALIKE+J). (**B**) †*Sooinichthys* excluded, reconstruction under the best fitting model (BAYAREALIKE+J). (**C**) †*Acronichthys* and †*Nardonoides* excluded, reconstruction under the best fitting model (BAYAREALIKE+J). (**D**) All fossil taxa included, reconstruction under Liu et al.’s chosen model (DEC). (**E**) †*Sooinichthys* excluded, reconstruction under Liu et al.’s chosen model (DEC). (**F**) †*Acronichthys* and †*Nardonoides* excluded, reconstruction under Liu et al.’s chosen model (DEC).

**Fig SX05.**
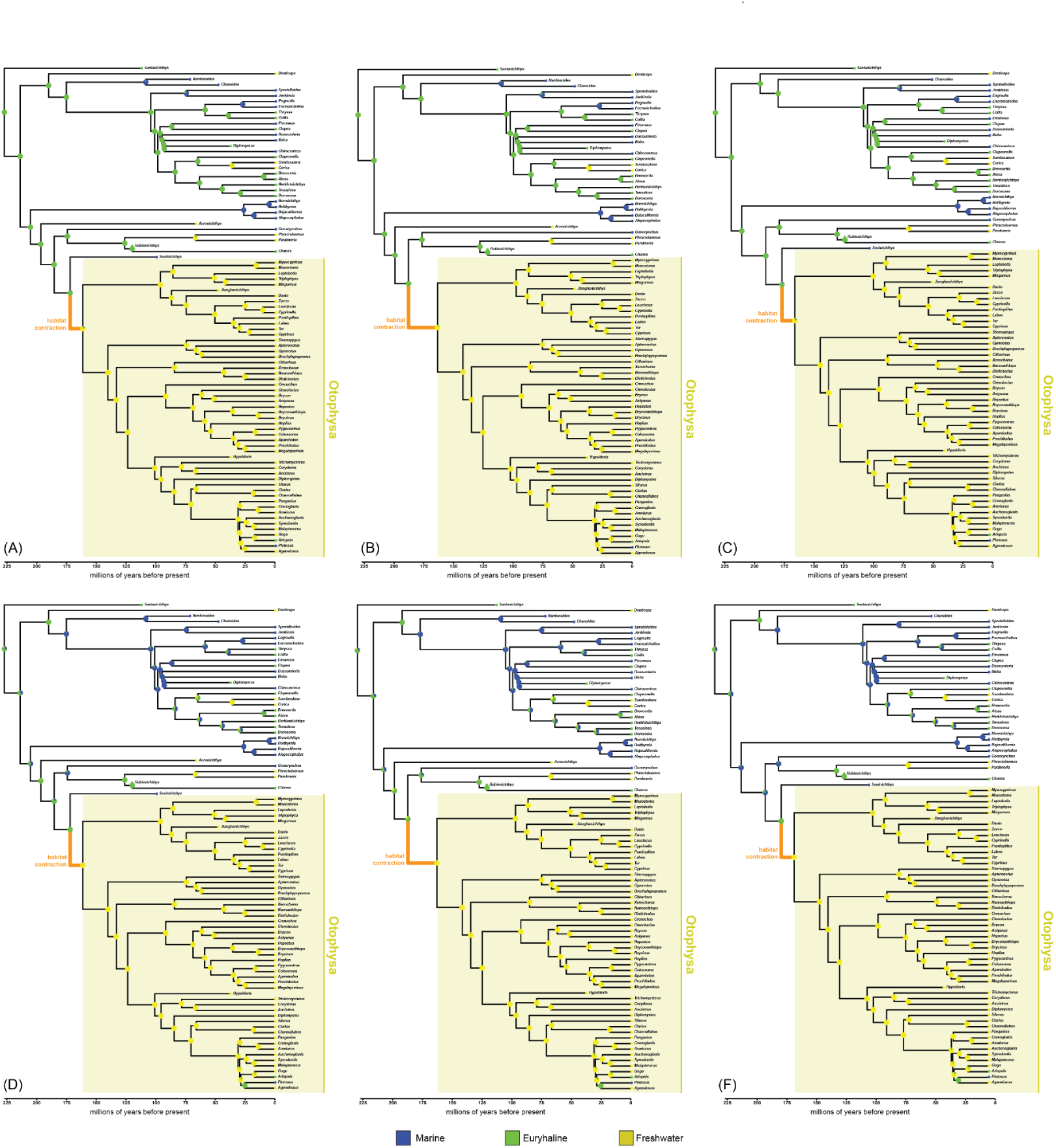
BioGeoBEARS results using best supported topology, rooted on †*Santanichthys*, with †*Santanichthys* coded as euryhaline, and with different combinations of fossil sampling and model. (**A**) All fossil taxa included, reconstruction under the best fitting model (BAYAREALIKE+J). (**B**) †*Sooinichthys* excluded, reconstruction under the best fitting model (BAYAREALIKE+J). (**C**) †*Acronichthys* and †*Nardonoides* excluded, reconstruction under the best fitting model (BAYAREALIKE+J). (**D**) All fossil taxa included, reconstruction under Liu et al.’s chosen model (DEC). (**E**) †*Sooinichthys* excluded, reconstruction under Liu et al.’s chosen model (DEC). (**F**) †*Acronichthys* and †*Nardonoides* excluded, reconstruction under Liu et al.’s chosen model (DEC).

**Fig SX06.**
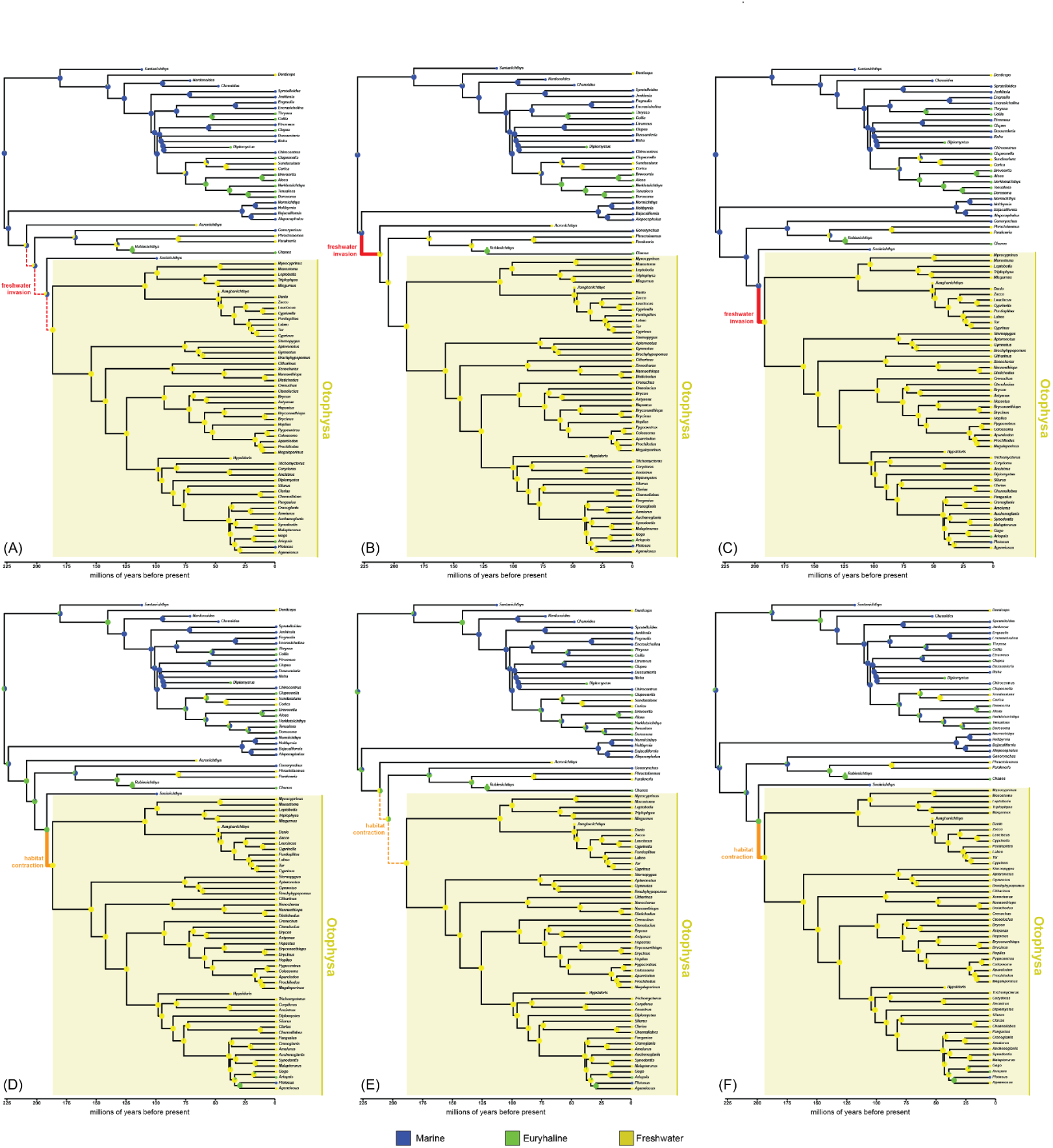
BioGeoBEARS results using best supported topology, rooted on Clupeiformes–†*Santanichthys*, with †*Santanichthys* coded as marine, and with different combinations of fossil sampling and model. (**A**) All fossil taxa included, reconstruction under the best fitting model (BAYAREALIKE+J). (**B**) †*Sooinichthys* excluded, reconstruction under the best fitting model (BAYAREALIKE+J). (**C**) †*Acronichthys* and †*Nardonoides* excluded, reconstruction under the best fitting model (BAYAREALIKE+J). (**D**) All fossil taxa included, reconstruction under Liu et al.’s chosen model (DEC). (**E**) †*Sooinichthys* excluded, reconstruction under Liu et al.’s chosen model (DEC). (**F**) †*Acronichthys* and †*Nardonoides* excluded, reconstruction under Liu et al.’s chosen model (DEC).

**Fig SX07.**
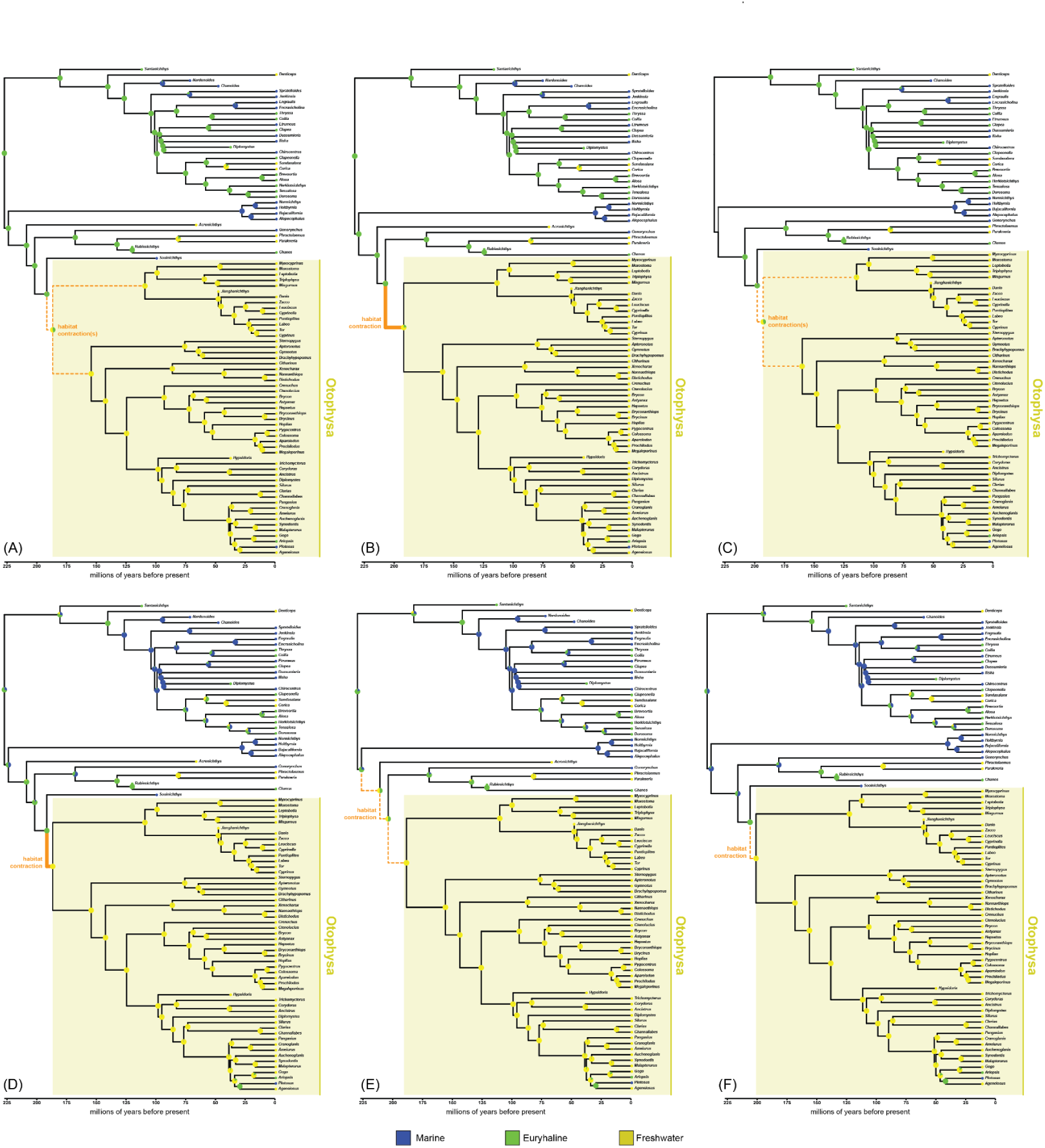
BioGeoBEARS results using best supported topology, rooted on Clupeiformes–†*Santanichthys*, with †*Santanichthys* coded as euryhaline, and with different combinations of fossil sampling and model. (**A**) All fossil taxa included, reconstruction under the best fitting model (BAYAREALIKE+J). (**B**) †*Sooinichthys* excluded, reconstruction under the best fitting model (BAYAREALIKE+J). (**C**) †*Acronichthys* and †*Nardonoides* excluded, reconstruction under the best fitting model (BAYAREALIKE+J). (**D**) All fossil taxa included, reconstruction under Liu et al.’s chosen model (DEC). (**E**) †*Sooinichthys* excluded, reconstruction under Liu et al.’s chosen model (DEC). (**F**) †*Acronichthys* and †*Nardonoides* excluded, reconstruction under Liu et al.’s chosen model (DEC).

**Fig SX08.**
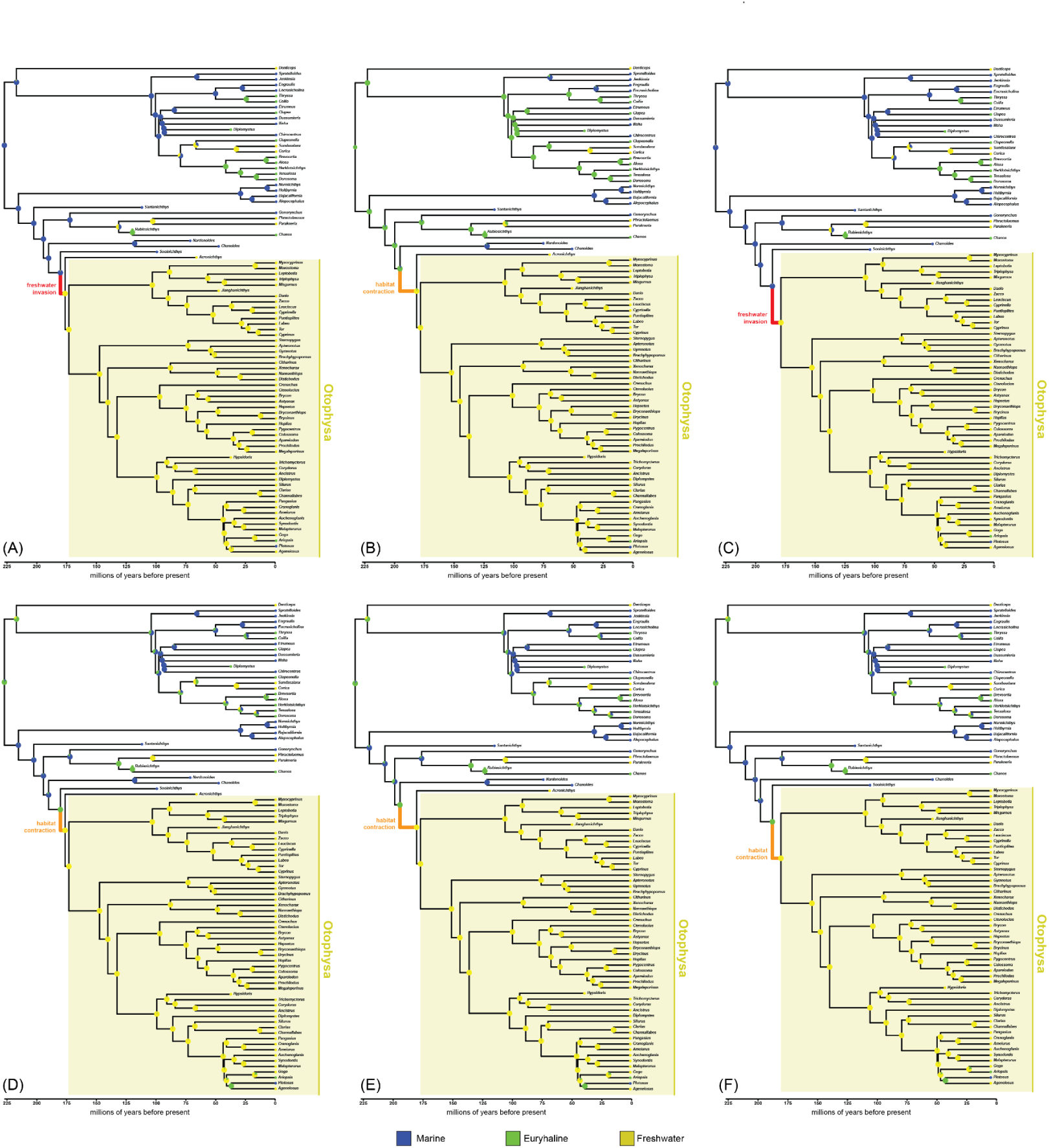
BioGeoBEARS results using second best supported topology, with †*Santanichthys* coded as marine, and with different combinations of fossil sampling and model. (**A**) All fossil taxa included, reconstruction under the best fitting model (BAYAREALIKE+J). (**B**) †*Sooinichthys* excluded, reconstruction under the best fitting model (BAYAREALIKE+J). (**C**) †*Acronichthys* and †*Nardonoides* excluded, reconstruction under the best fitting model (BAYAREALIKE+J). (**D**) All fossil taxa included, reconstruction under Liu et al.’s chosen model (DEC). (**E**) †*Sooinichthys* excluded, reconstruction under Liu et al.’s chosen model (DEC). (**F**) †*Acronichthys* and †*Nardonoides* excluded, reconstruction under Liu et al.’s chosen model (DEC).

**Fig SX09.**
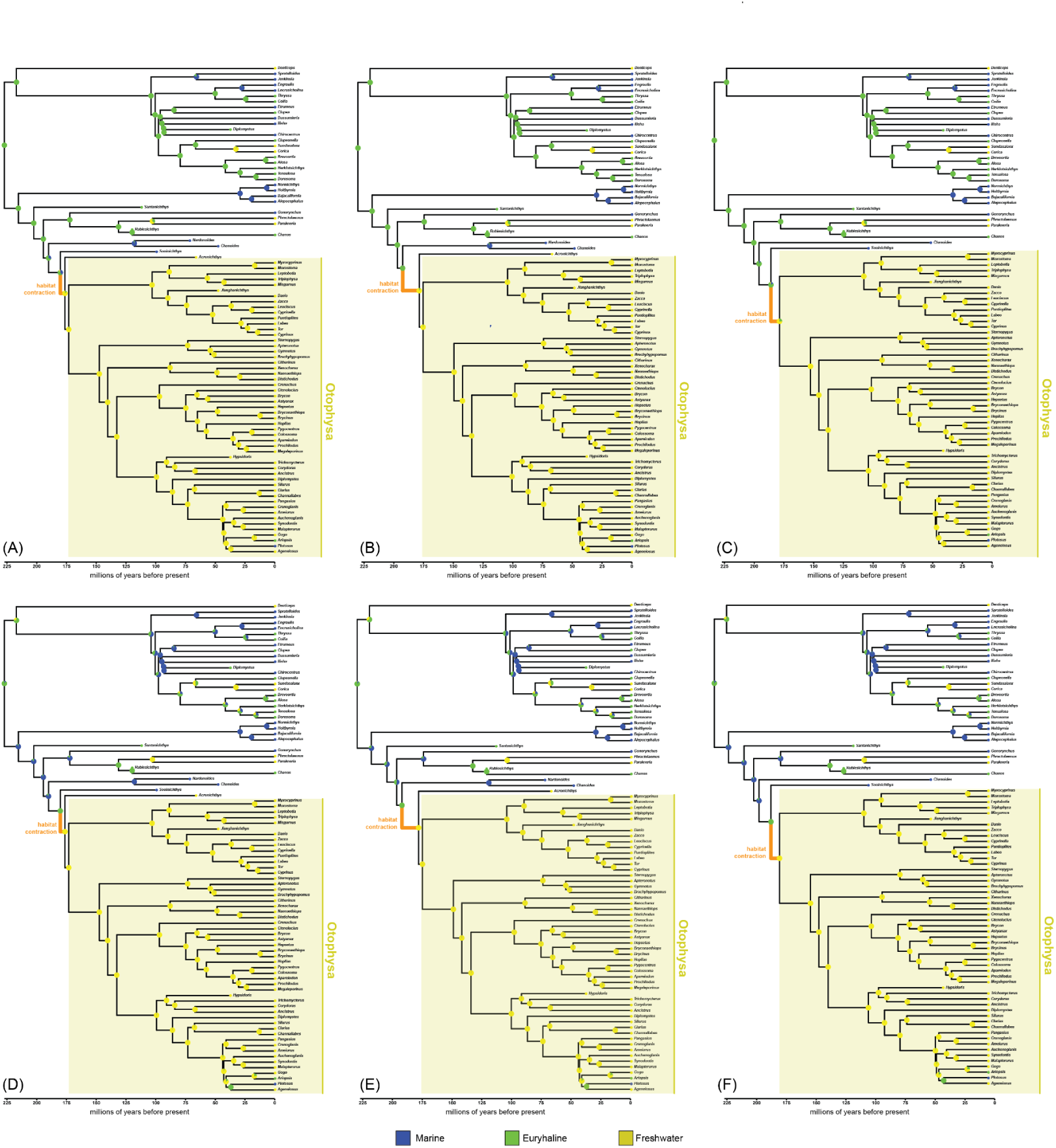
BioGeoBEARS results using second best supported topology, with †*Santanichthys* coded as euryhaline, and with different combinations of fossil sampling and model. (**A**) All fossil taxa included, reconstruction under the best fitting model (BAYAREALIKE+J). (**B**) †*Sooinichthys* excluded, reconstruction under the best fitting model (BAYAREALIKE+J). (**C**) †*Acronichthys* and †*Nardonoides* excluded, reconstruction under the best fitting model (BAYAREALIKE+J). (**D**) All fossil taxa included, reconstruction under Liu et al.’s chosen model (DEC). (**E**) †*Sooinichthys* excluded, reconstruction under Liu et al.’s chosen model (DEC). (**F**) †*Acronichthys* and †*Nardonoides* excluded, reconstruction under Liu et al.’s chosen model (DEC).

**Fig SX10.**
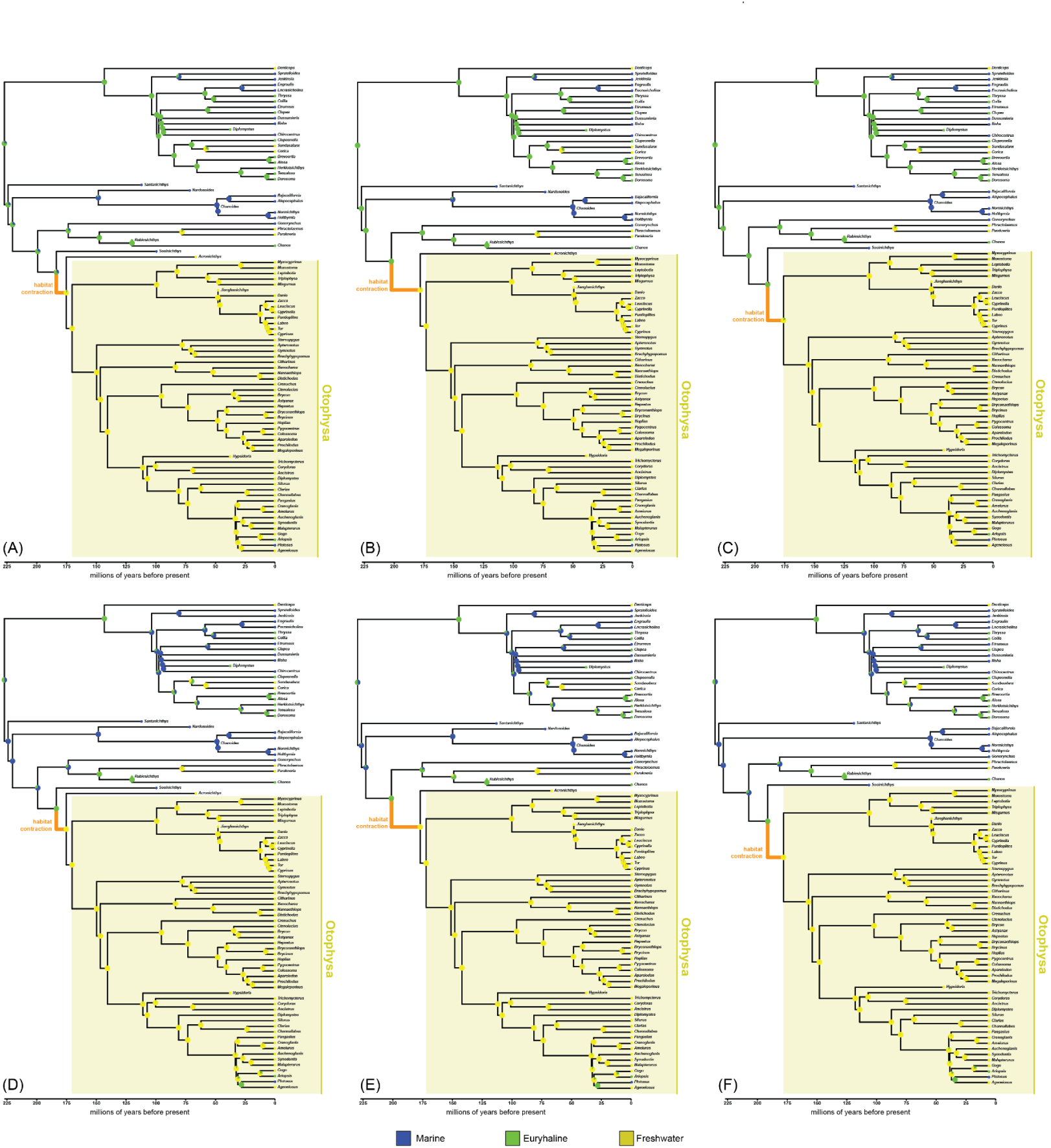
BioGeoBEARS results using third best supported topology, with †*Santanichthys* coded as marine, and with different combinations of fossil sampling and model. (**A**) All fossil taxa included, reconstruction under the best fitting model (BAYAREALIKE+J). (**B**) †*Sooinichthys* excluded, reconstruction under the best fitting model (BAYAREALIKE+J). (**C**) †*Acronichthys* and †*Nardonoides* excluded, reconstruction under the best fitting model (BAYAREALIKE+J). (**D**) All fossil taxa included, reconstruction under Liu et al.’s chosen model (DEC). (**E**) †*Sooinichthys* excluded, reconstruction under Liu et al.’s chosen model (DEC). (**F**) †*Acronichthys* and †*Nardonoides* excluded, reconstruction under Liu et al.’s chosen model (DEC).

**Fig SX11.**
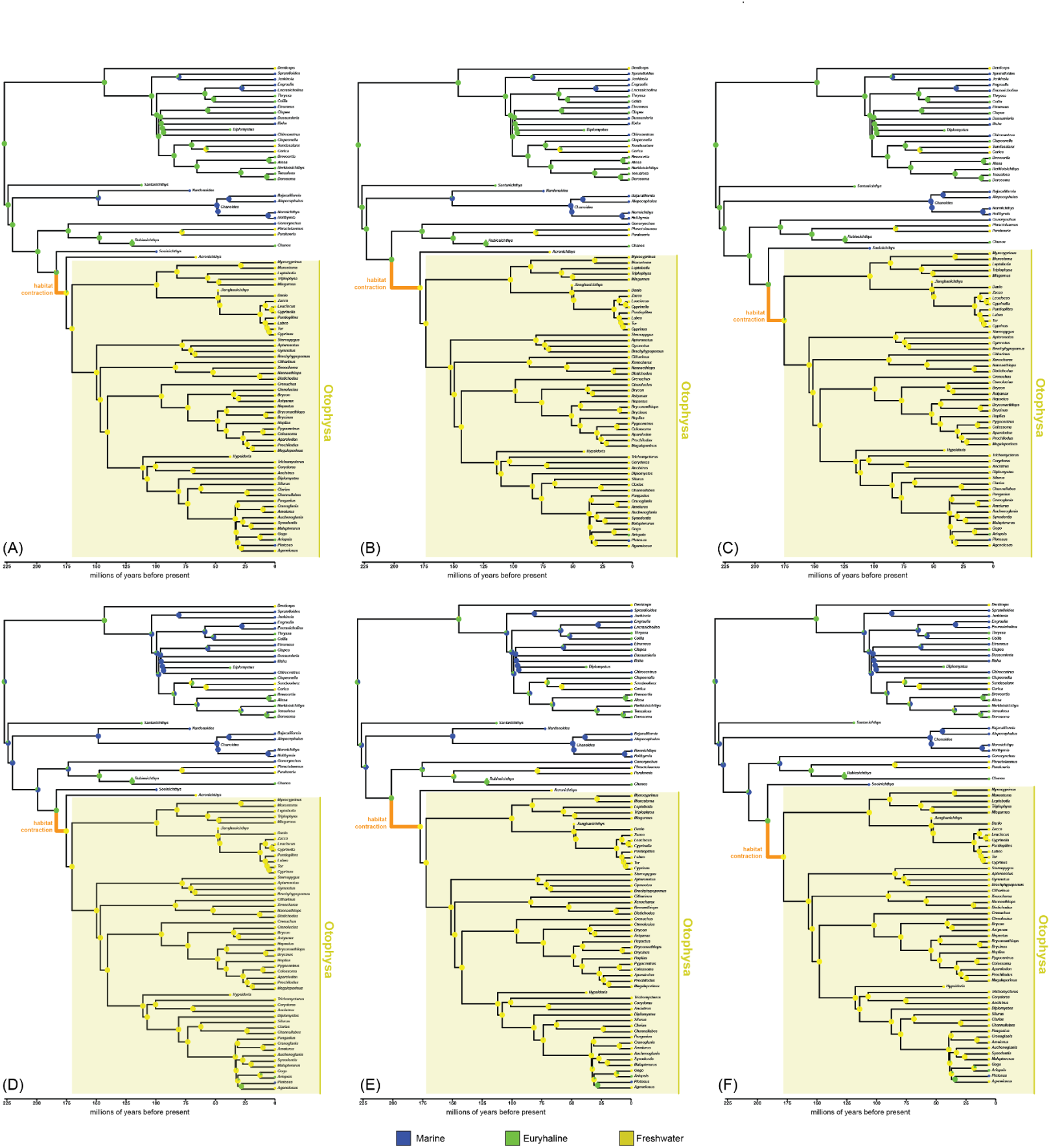
BioGeoBEARS results using third best supported topology, with †*Santanichthys* coded as euryhaline, and with different combinations of fossil sampling and model. (**A**) All fossil taxa included, reconstruction under the best fitting model (BAYAREALIKE+J). (**B**) †*Sooinichthys* excluded, reconstruction under the best fitting model (BAYAREALIKE+J). (**C**) †*Acronichthys* and †*Nardonoides* excluded, reconstruction under the best fitting model (BAYAREALIKE+J). (**D**) All fossil taxa included, reconstruction under Liu et al.’s chosen model (DEC). (**E**) †*Sooinichthys* excluded, reconstruction under Liu et al.’s chosen model (DEC). (**F**) †*Acronichthys* and †*Nardonoides* excluded, reconstruction under Liu et al.’s chosen model (DEC).

**Fig SX12.**
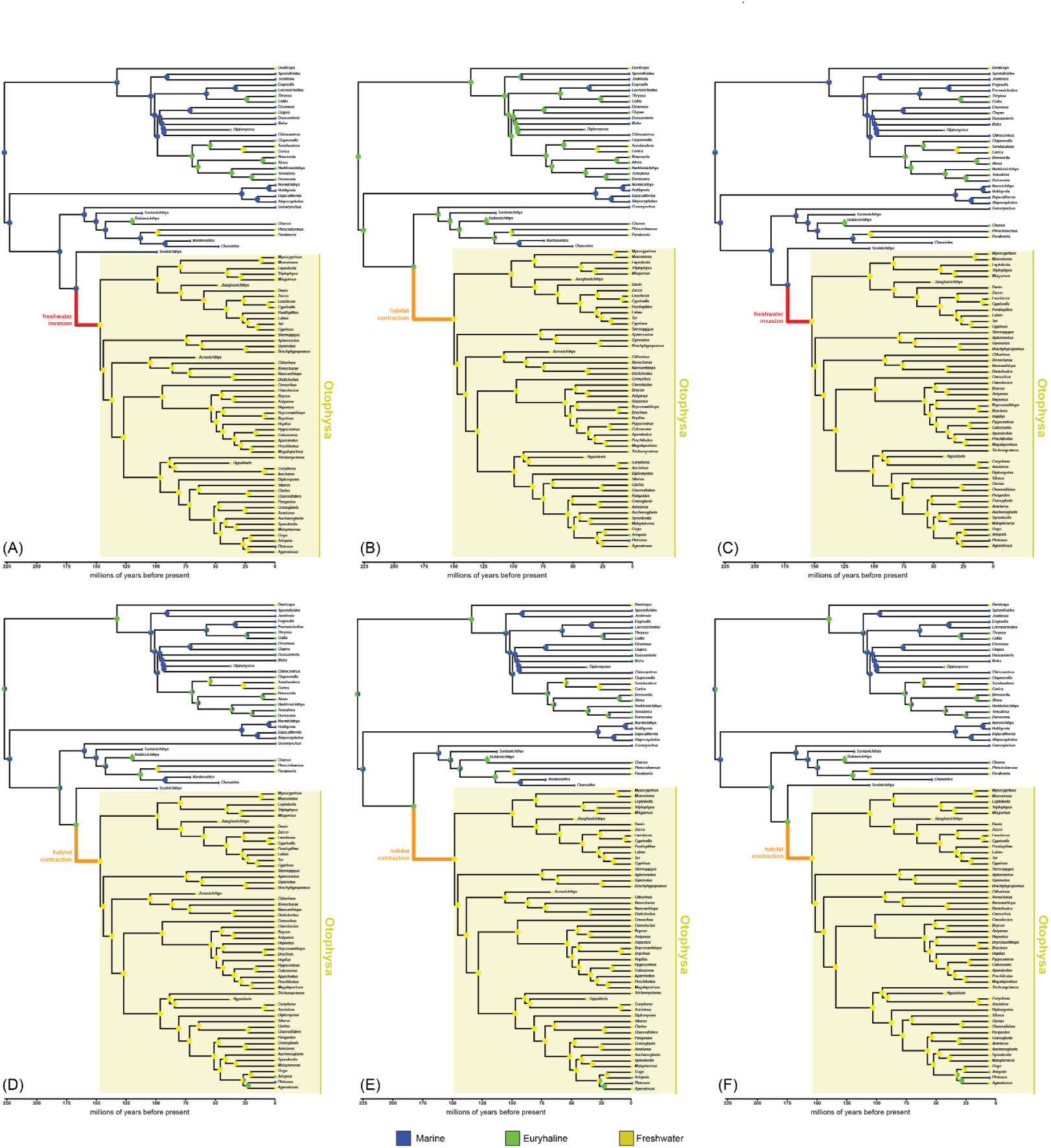
BioGeoBEARS results using fourth best supported topology, with †*Santanichthys* coded as marine, and with different combinations of fossil sampling and model. (**A**) All fossil taxa included, reconstruction under the best fitting model (BAYAREALIKE+J). (**B**) †*Sooinichthys* excluded, reconstruction under the best fitting model (BAYAREALIKE+J). (**C**) †*Acronichthys* and †*Nardonoides* excluded, reconstruction under the best fitting model (BAYAREALIKE+J). (**D**) All fossil taxa included, reconstruction under Liu et al.’s chosen model (DEC). (**E**) †*Sooinichthys* excluded, reconstruction under Liu et al.’s chosen model (DEC). (**F**) †*Acronichthys* and †*Nardonoides* excluded, reconstruction under Liu et al.’s chosen model (DEC).

**Fig SX13.**
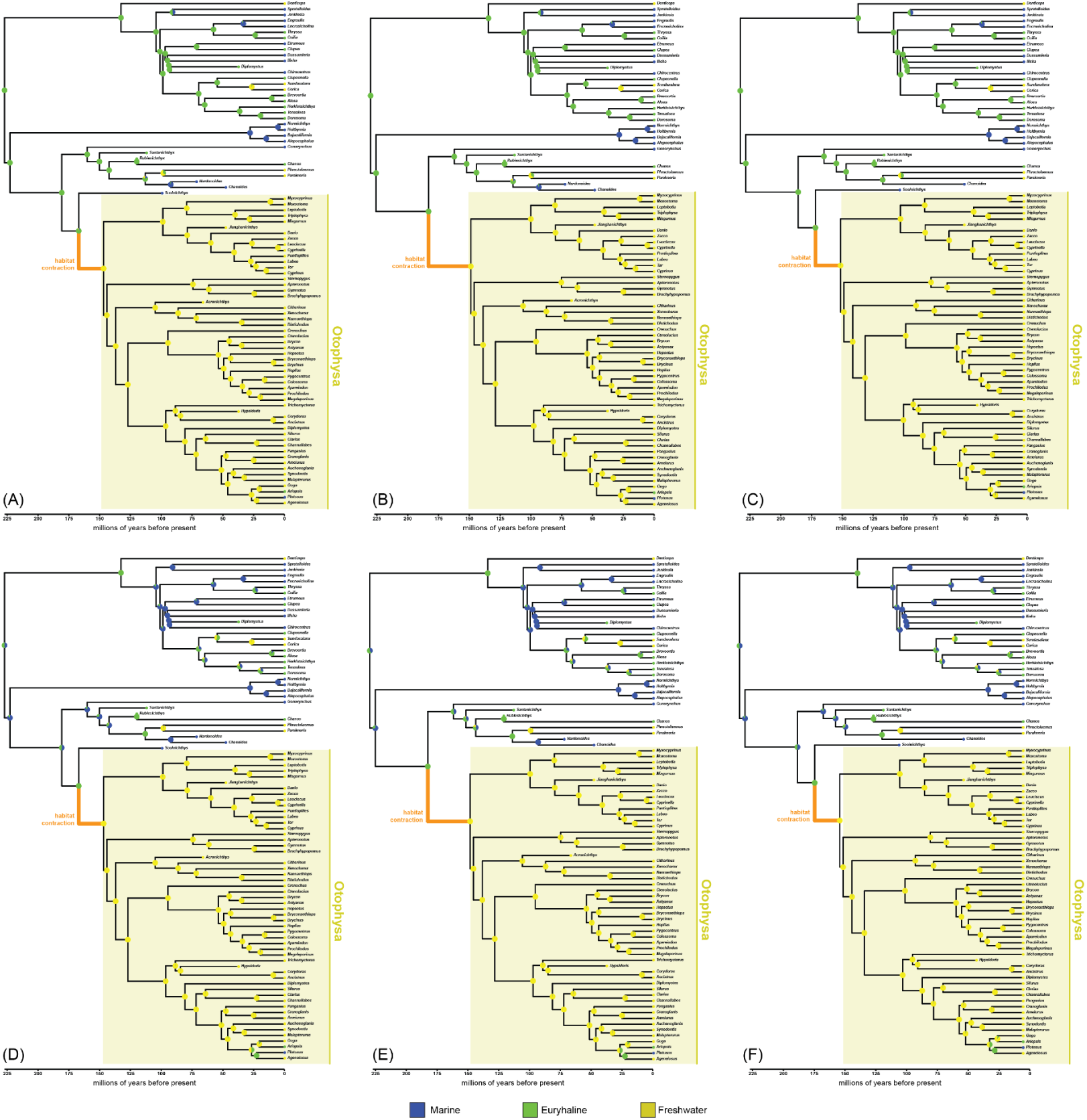
BioGeoBEARS results using fourth best supported topology, with †*Santanichthys* coded as euryhaline, and with different combinations of fossil sampling and model. (**A**) All fossil taxa included, reconstruction under the best fitting model (BAYAREALIKE+J). (**B**) †*Sooinichthys* excluded, reconstruction under the best fitting model (BAYAREALIKE+J). (**C**) †*Acronichthys* and †*Nardonoides* excluded, reconstruction under the best fitting model (BAYAREALIKE+J). (**D**) All fossil taxa included, reconstruction under Liu et al.’s chosen model (DEC). (**E**) †*Sooinichthys* excluded, reconstruction under Liu et al.’s chosen model (DEC). (**F**) †*Acronichthys* and †*Nardonoides* excluded, reconstruction under Liu et al.’s chosen model (DEC).

**Fig SX14.**
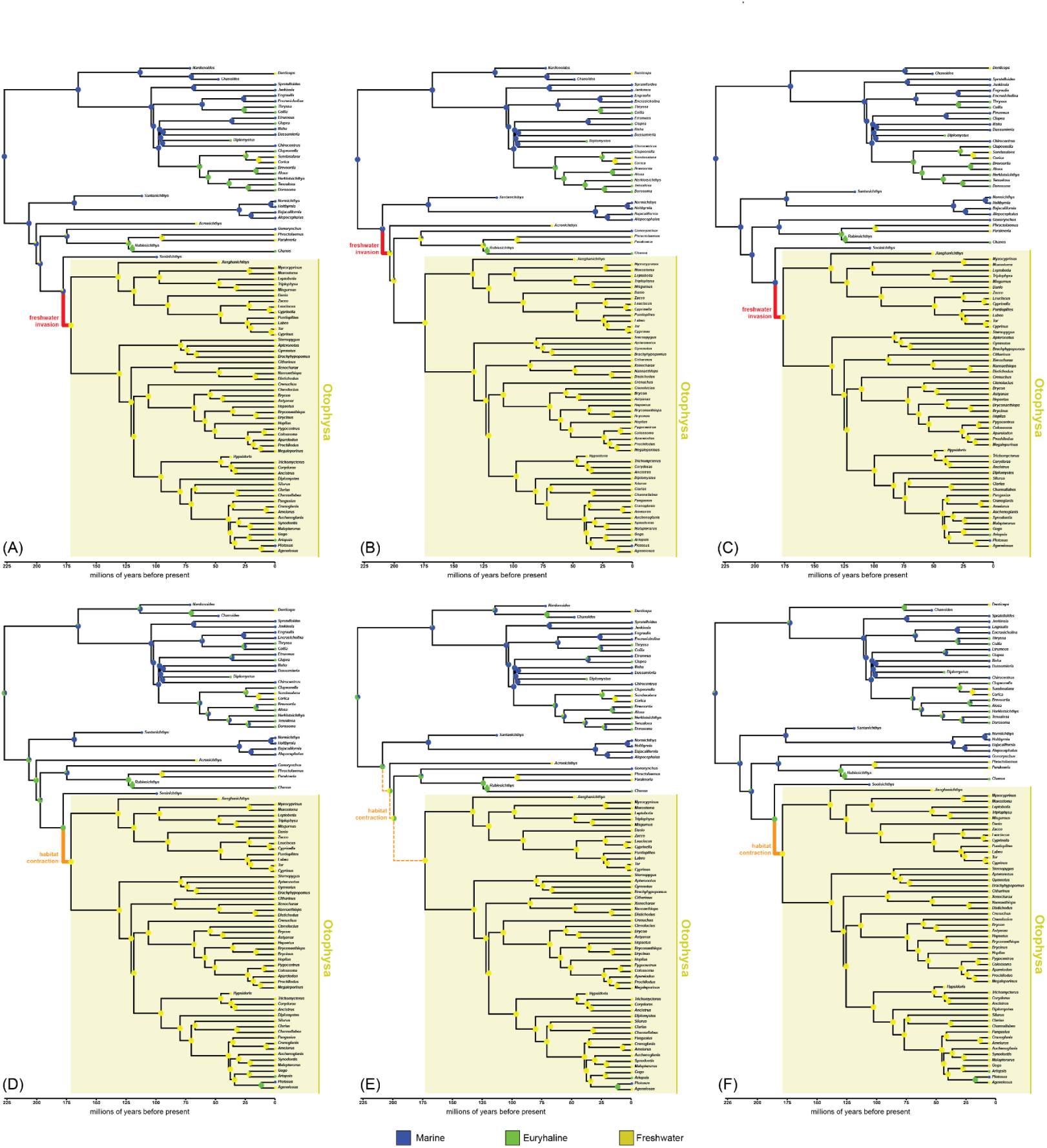
BioGeoBEARS results using fifth best supported topology, with †*Santanichthys* coded as marine, and with different combinations of fossil sampling and model. (**A**) All fossil taxa included, reconstruction under the best fitting model (BAYAREALIKE+J). (**B**) †*Sooinichthys* excluded, reconstruction under the best fitting model (BAYAREALIKE+J). (**C**) †*Acronichthys* and †*Nardonoides* excluded, reconstruction under the best fitting model (BAYAREALIKE+J). (**D**) All fossil taxa included, reconstruction under Liu et al.’s chosen model (DEC). (**E**) †*Sooinichthys* excluded, reconstruction under Liu et al.’s chosen model (DEC). (**F**) †*Acronichthys* and †*Nardonoides* excluded, reconstruction under Liu et al.’s chosen model (DEC).

**Fig SX15.**
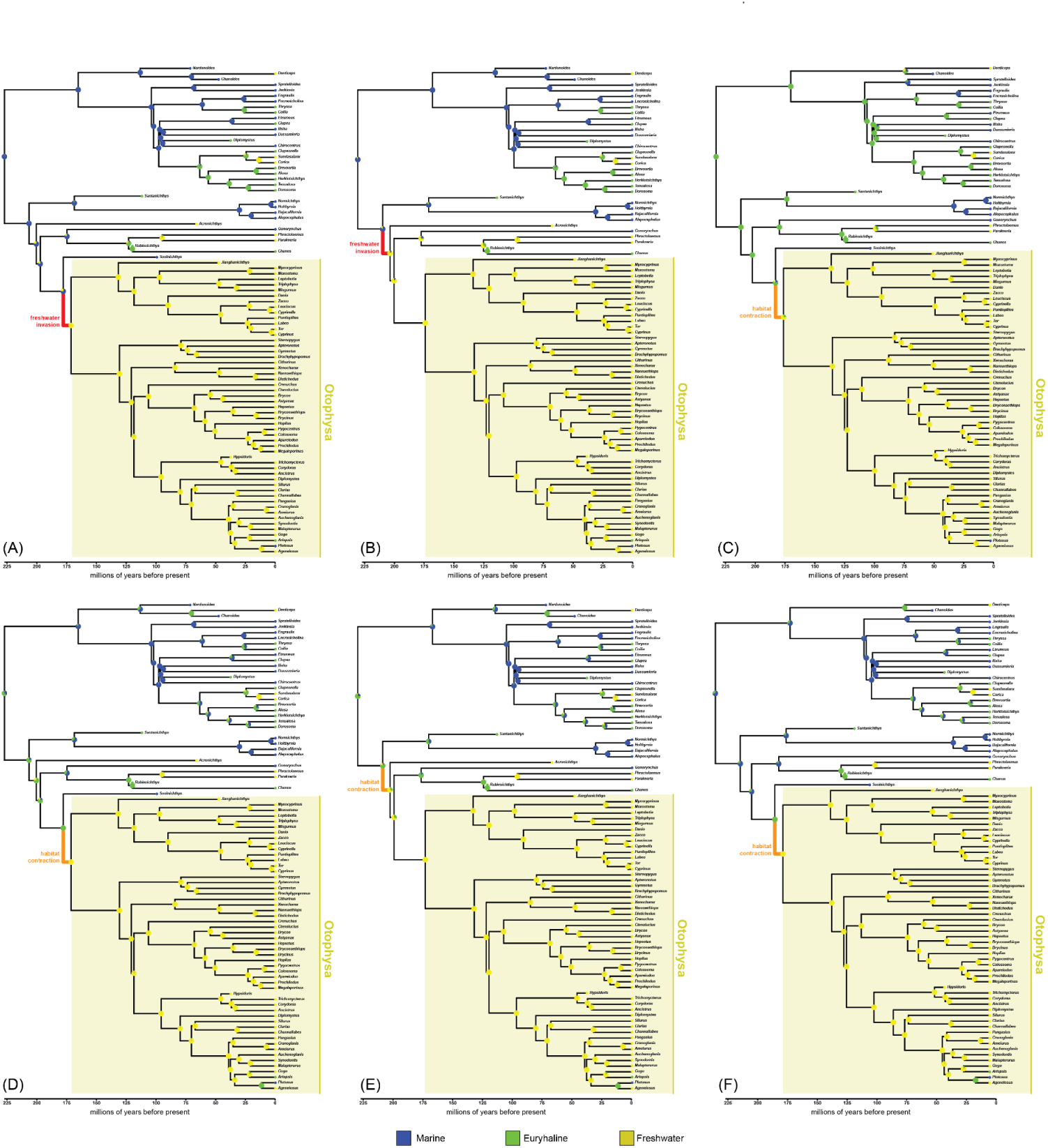
BioGeoBEARS results using fifth best supported topology, with †*Santanichthys* coded as euryhaline, and with different combinations of fossil sampling and model. (**A**) All fossil taxa included, reconstruction under the best fitting model (BAYAREALIKE+J). (**B**) †*Sooinichthys* excluded, reconstruction under the best fitting model (BAYAREALIKE+J). (**C**) †*Acronichthys* and †*Nardonoides* excluded, reconstruction under the best fitting model (DEC, also Liu et al.’s chosen model). **Note:** Across 66 trials, this is the only instance in which (BAYAREALIKE+J) is not the best fitting model. (**D**) All fossil taxa included, reconstruction under Liu et al.’s chosen model (DEC). (**E**) †*Sooinichthys* excluded, reconstruction under Liu et al.’s chosen model (DEC). (**F**) †*Acronichthys* and †*Nardonoides* excluded, reconstruction under the second best fitting model (BAYAREALIKE+J).

**Fig SX16.**
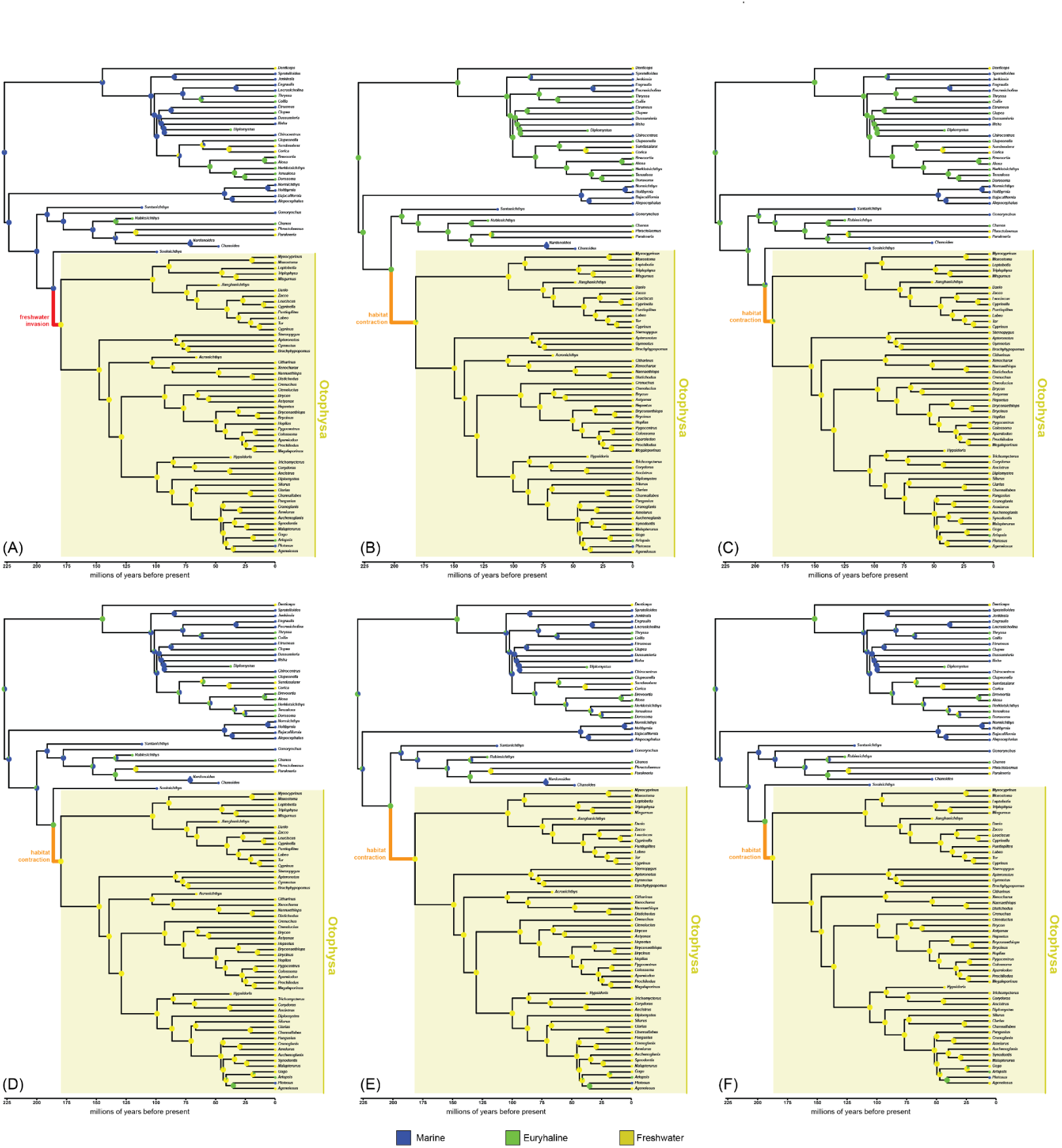
BioGeoBEARS results using the topology recovered with the matrix of Liu et al., revised coding for the Weberian apparatus (traits 1–27), with †*Santanichthys* coded as marine, and with different combinations of fossil sampling and model. (**A**) All fossil taxa included, reconstruction under the best fitting model (BAYAREALIKE+J). (**B**) †*Sooinichthys* excluded, reconstruction under the best fitting model (BAYAREALIKE+J). (**C**) †*Acronichthys* and †*Nardonoides* excluded, reconstruction under the best fitting model (BAYAREALIKE+J). (**D**) All fossil taxa included, reconstruction under Liu et al.’s chosen model (DEC). (**E**) †*Sooinichthys* excluded, reconstruction under Liu et al.’s chosen model (DEC). (**F**) †*Acronichthys* and †*Nardonoides* excluded, reconstruction under Liu et al.’s chosen model (DEC).

**Fig SX17.**
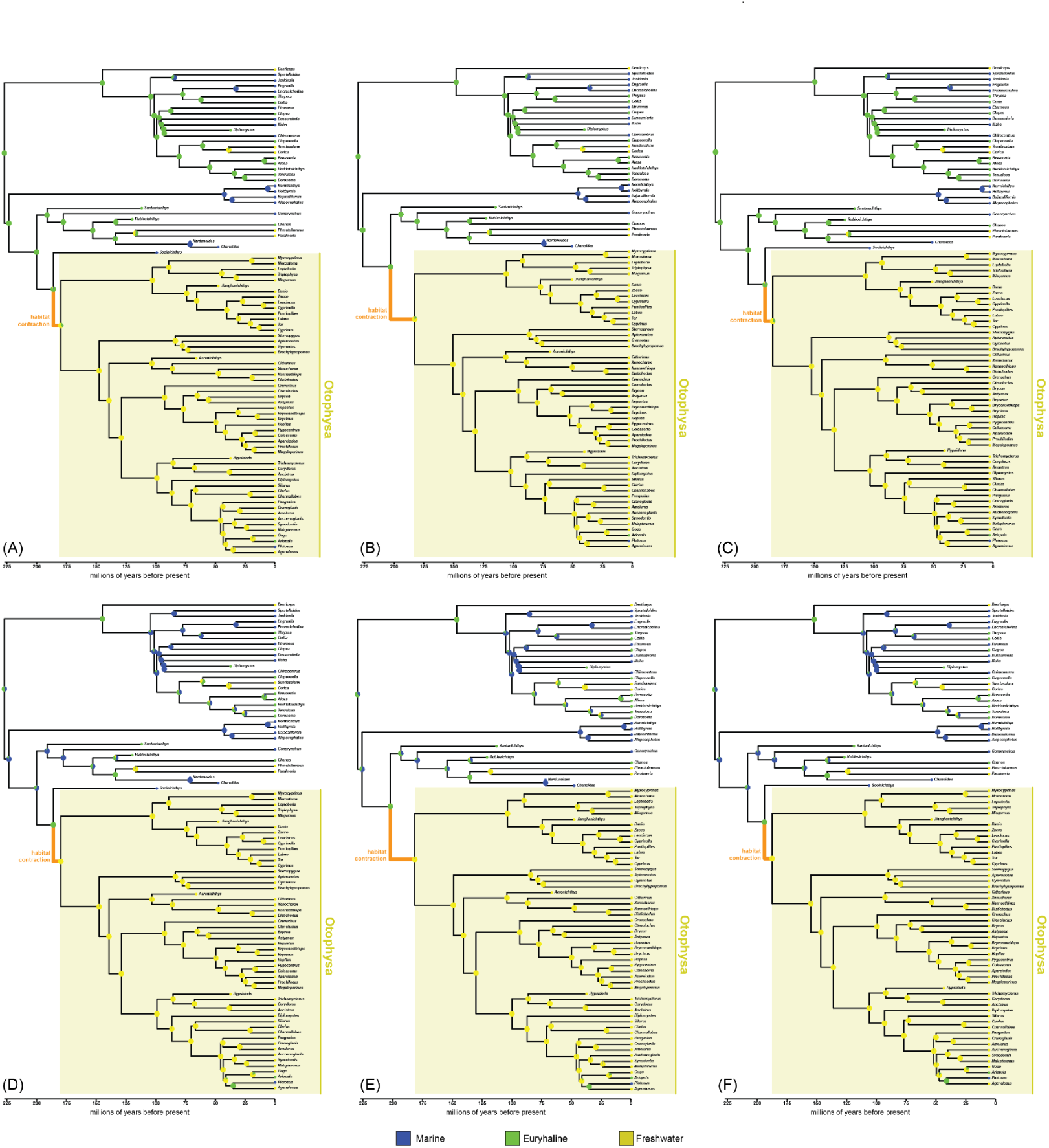
BioGeoBEARS results using the topology recovered with the matrix of Liu et al., revised coding for the Weberian apparatus (traits 1–27), with †*Santanichthys* coded as euryhaline, and with different combinations of fossil sampling and model. (**A**) All fossil taxa included, reconstruction under the best fitting model (BAYAREALIKE+J). (**B**) †*Sooinichthys* excluded, reconstruction under the best fitting model (BAYAREALIKE+J). (**C**) †*Acronichthys* and †*Nardonoides* excluded, reconstruction under the best fitting model (BAYAREALIKE+J). (**D**) All fossil taxa included, reconstruction under Liu et al.’s chosen model (DEC). (**E**) †*Sooinichthys* excluded, reconstruction under Liu et al.’s chosen model (DEC). (**F**) †*Acronichthys* and †*Nardonoides* excluded, reconstruction under Liu et al.’s chosen model (DEC).

**Fig SX18.**
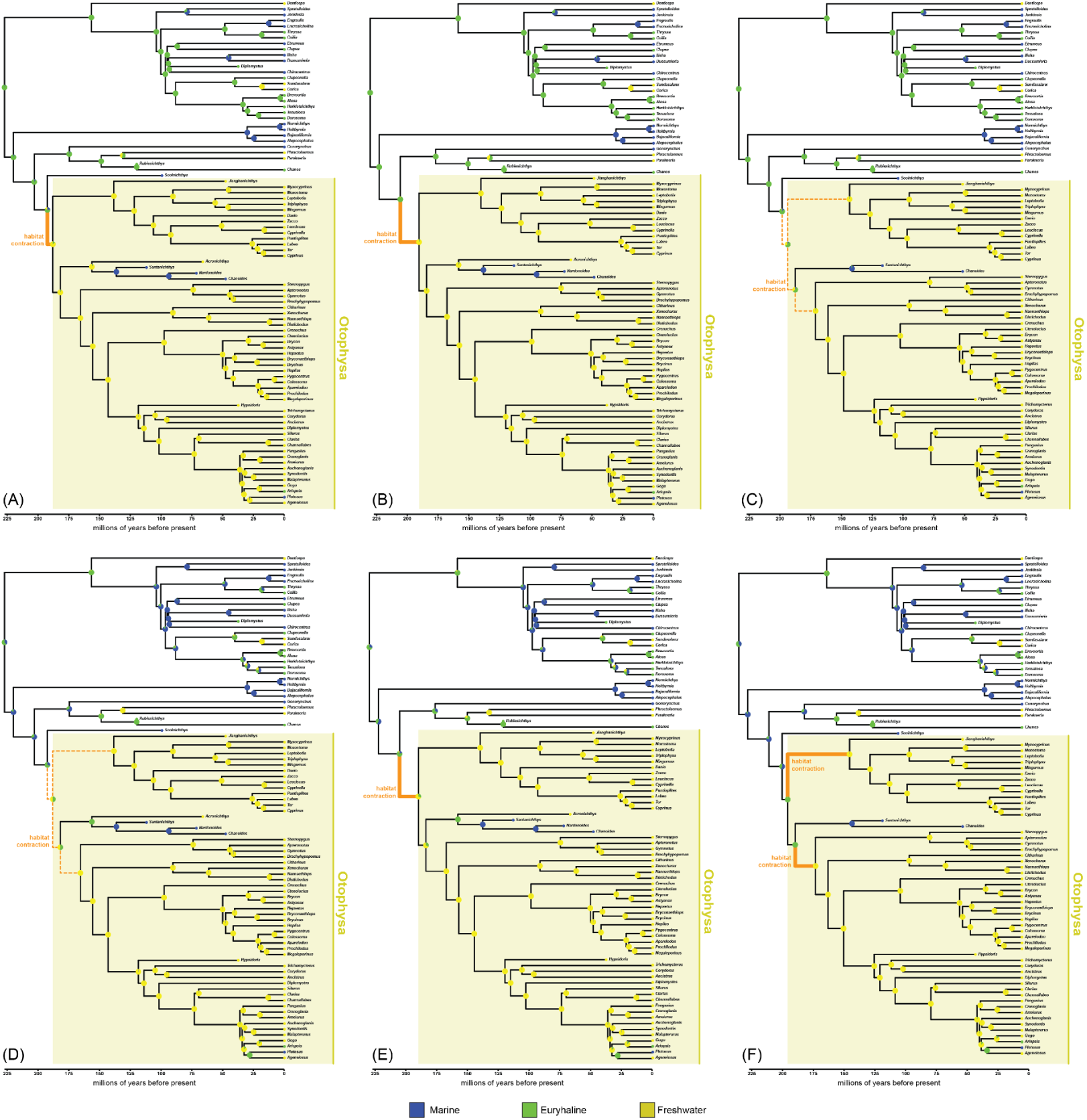
BioGeoBEARS results using the topology recovered with the matrix of Liu et al., revised coding for the caudal region (traits 28–41), with †*Santanichthys* coded as marine, and with different combinations of fossil sampling and model. (**A**) All fossil taxa included, reconstruction under the best fitting model (BAYAREALIKE+J). (**B**) †*Sooinichthys* excluded, reconstruction under the best fitting model (BAYAREALIKE+J). (**C**) †*Acronichthys* and †*Nardonoides* excluded, reconstruction under the best fitting model (BAYAREALIKE+J). (**D**) All fossil taxa included, reconstruction under Liu et al.’s chosen model (DEC). (**E**) †*Sooinichthys* excluded, reconstruction under Liu et al.’s chosen model (DEC). (**F**) †*Acronichthys* and †*Nardonoides* excluded, reconstruction under Liu et al.’s chosen model (DEC).

**Fig SX19.**
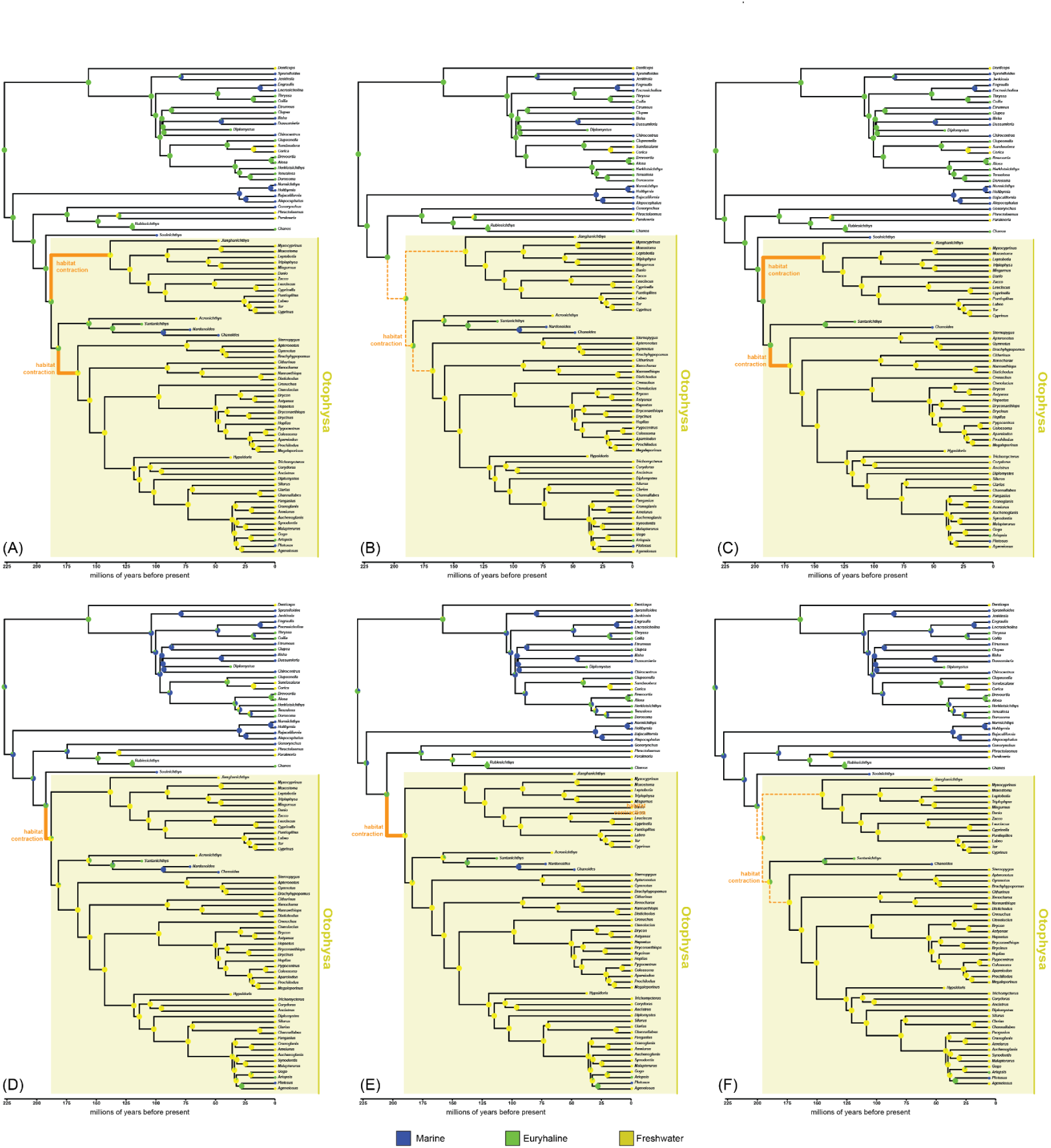
BioGeoBEARS results using the topology recovered with the matrix of Liu et al., revised coding for the caudal region (traits 28–41), with †*Santanichthys* coded as euryhaline, and with different combinations of fossil sampling and model. (**A**) All fossil taxa included, reconstruction under the best fitting model (BAYAREALIKE+J). (**B**) †*Sooinichthys* excluded, reconstruction under the best fitting model (BAYAREALIKE+J). (**C**) †*Acronichthys* and †*Nardonoides* excluded, reconstruction under the best fitting model (BAYAREALIKE+J). (**D**) All fossil taxa included, reconstruction under Liu et al.’s chosen model (DEC). (**E**) †*Sooinichthys* excluded, reconstruction under Liu et al.’s chosen model (DEC). (**F**) †*Acronichthys* and †*Nardonoides* excluded, reconstruction under Liu et al.’s chosen model (DEC).

**Fig SX20.**
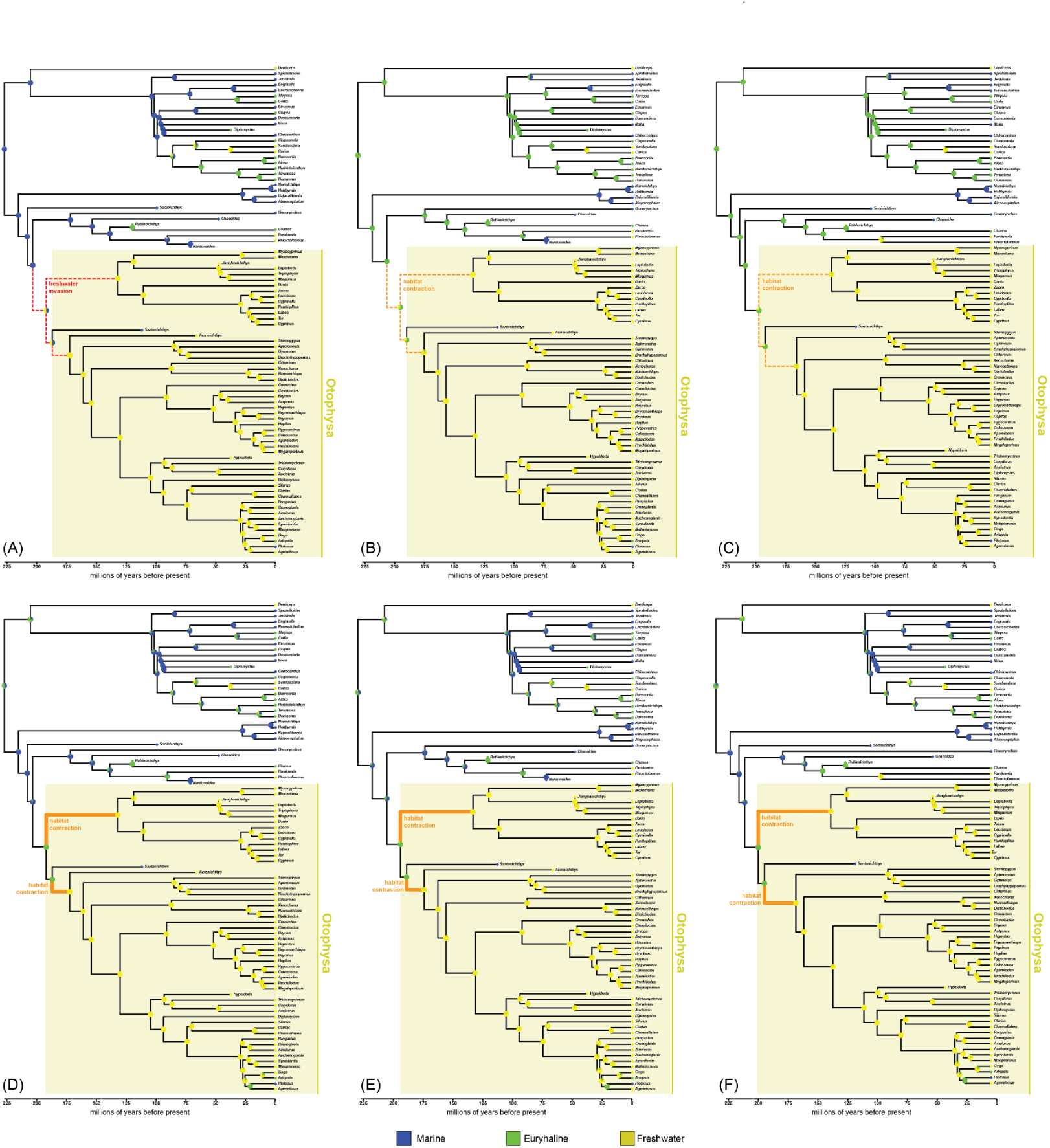
BioGeoBEARS results using the topology recovered with the matrix of Liu et al., revised coding for the jaw (traits 42–49), with †*Santanichthys* coded as marine, and with different combinations of fossil sampling and model. (**A**) All fossil taxa included, reconstruction under the best fitting model (BAYAREALIKE+J). (**B**) †*Sooinichthys* excluded, reconstruction under the best fitting model (BAYAREALIKE+J). (**C**) †*Acronichthys* and †*Nardonoides* excluded, reconstruction under the best fitting model (BAYAREALIKE+J). (**D**) All fossil taxa included, reconstruction under Liu et al.’s chosen model (DEC). (**E**) †*Sooinichthys* excluded, reconstruction under Liu et al.’s chosen model (DEC). (**F**) †*Acronichthys* and †*Nardonoides* excluded, reconstruction under Liu et al.’s chosen model (DEC).

**Fig SX21.**
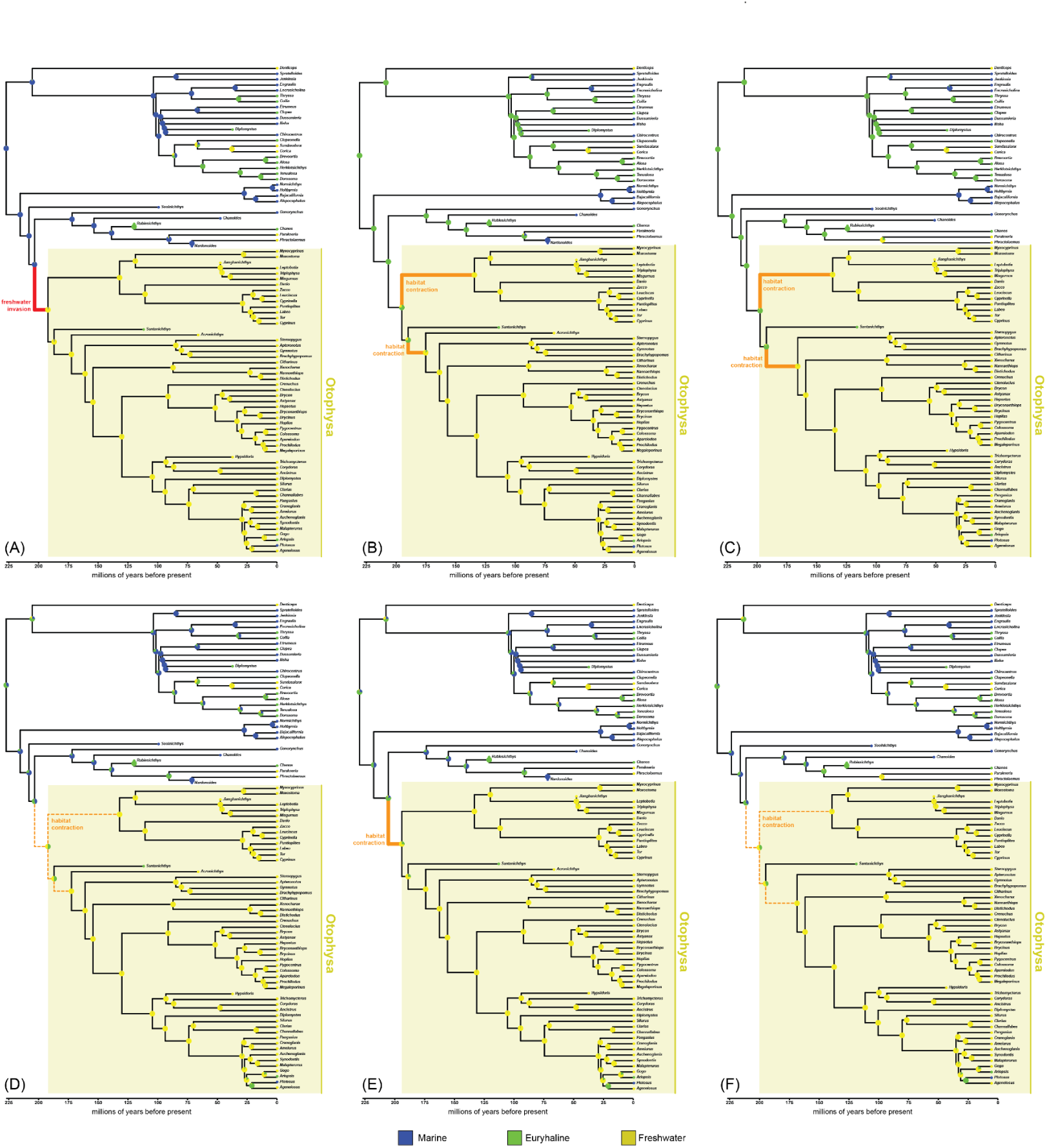
BioGeoBEARS results using the topology recovered with the matrix of Liu et al., revised coding for the jaw (traits 42–49), with †*Santanichthys* coded as euryhaline, and with different combinations of fossil sampling and model. (**A**) All fossil taxa included, reconstruction under the best fitting model (BAYAREALIKE+J). (**B**) †*Sooinichthys* excluded, reconstruction under the best fitting model (BAYAREALIKE+J). (**C**) †*Acronichthys* and †*Nardonoides* excluded, reconstruction under the best fitting model (BAYAREALIKE+J). (**D**) All fossil taxa included, reconstruction under Liu et al.’s chosen model (DEC). (**E**) †*Sooinichthys* excluded, reconstruction under Liu et al.’s chosen model (DEC). (**F**) †*Acronichthys* and †*Nardonoides* excluded, reconstruction under Liu et al.’s chosen model (DEC).

**Fig SX22.**
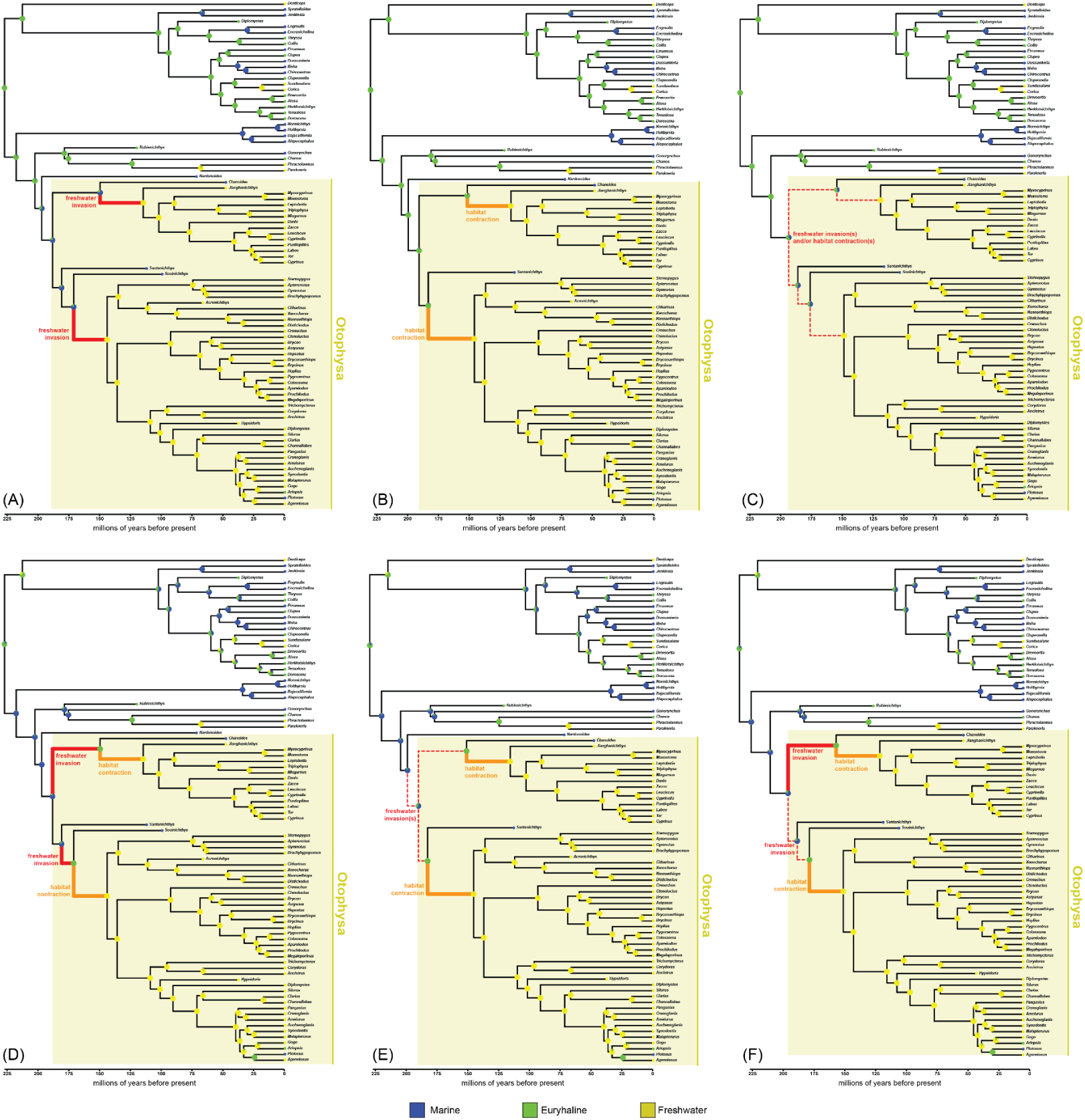
BioGeoBEARS results using the fossil placements recovered by Liu et al., with †*Santanichthys* coded as marine, and with different combinations of fossil sampling and model. (**A**) All fossil taxa included, reconstruction under the best fitting model (BAYAREALIKE+J). (**B**) †*Sooinichthys* excluded, reconstruction under the best fitting model (BAYAREALIKE+J). (**C**) †*Acronichthys* and †*Nardonoides* excluded, reconstruction under the best fitting model (BAYAREALIKE+J). (**D**) All fossil taxa included, reconstruction under Liu et al.’s chosen model (DEC). (**E**) †*Sooinichthys* excluded, reconstruction under Liu et al.’s chosen model (DEC). (**F**) †*Acronichthys* and †*Nardonoides* excluded, reconstruction under Liu et al.’s chosen model (DEC).

**Fig SX23.**
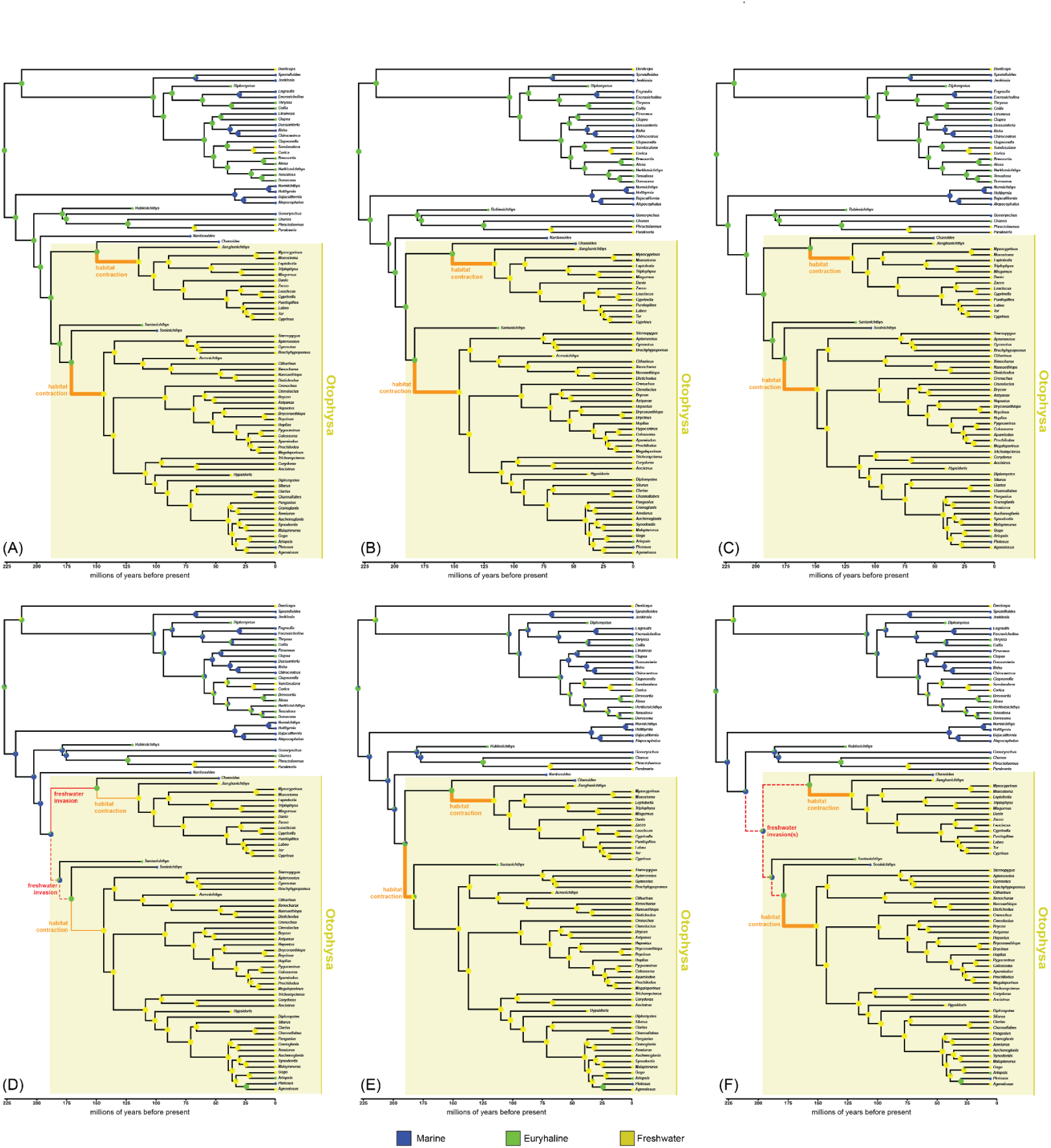
BioGeoBEARS results using the fossil placements recovered by Liu et al., with †*Santanichthys* coded as euryhaline, and with different combinations of fossil sampling and model. (**A**) All fossil taxa included, reconstruction under the best fitting model (BAYAREALIKE+J). (**B**) †*Sooinichthys* excluded, reconstruction under the best fitting model (BAYAREALIKE+J). (**C**) †*Acronichthys* and †*Nardonoides* excluded, reconstruction under the best fitting model (BAYAREALIKE+J). (**D**) All fossil taxa included, reconstruction under Liu et al.’s chosen model (DEC). (**E**) †*Sooinichthys* excluded, reconstruction under Liu et al.’s chosen model (DEC). (**F**) †*Acronichthys* and †*Nardonoides* excluded, reconstruction under Liu et al.’s chosen model (DEC).

**Fig SX24.**
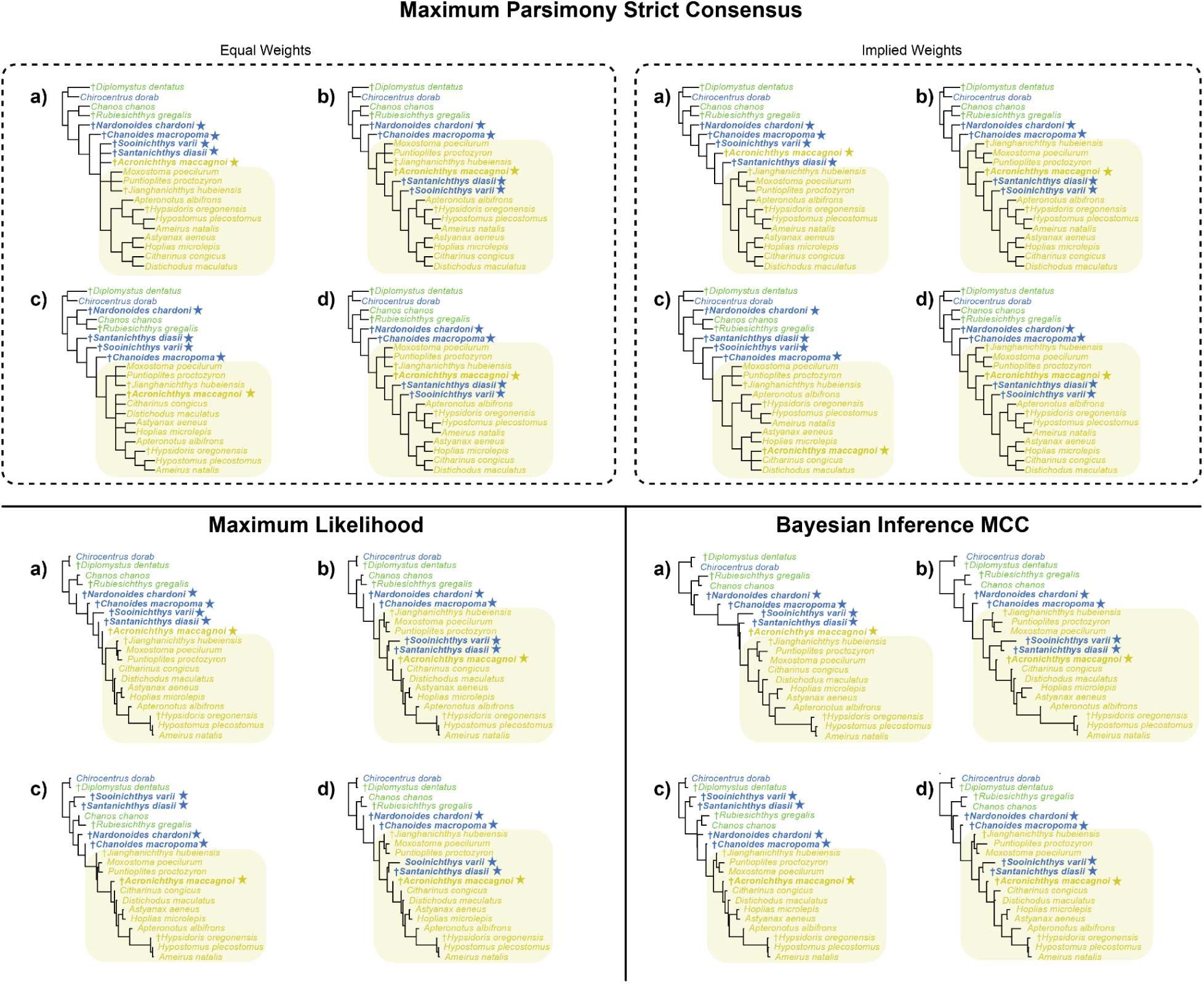
Morphological Trees estimated with combination matrices. Topologies estimated under 4 analytical frameworks: Equal weights maximum parsimony, implied weights maximum parsimony, maximum likelihood, and bayesian inference. Maximum parsimony trees are all strict consensus trees, with implied weights representing the topology shared by the most values of the concavity constant (*k*). Bayesian inference tree summarized in a maximum clade consensus tree. Taxa names that are colored were recoded here.

**Fig SX25.**
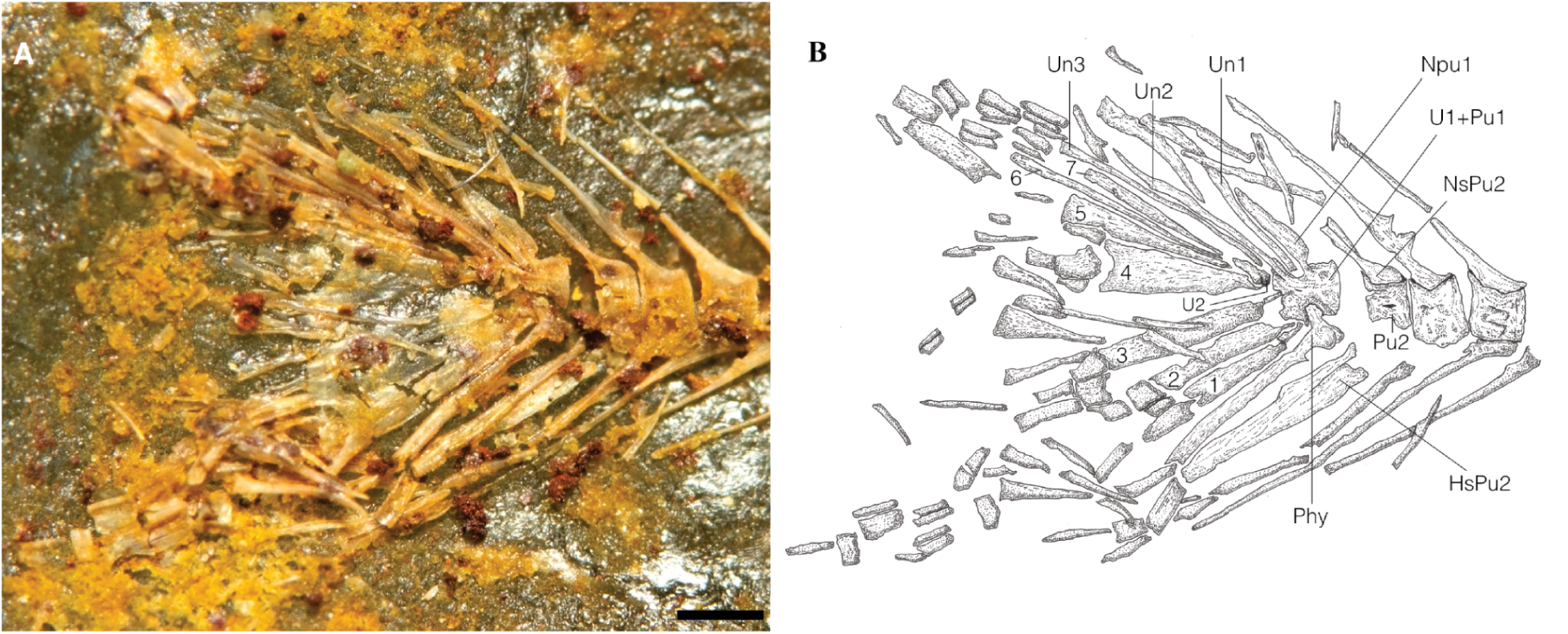
†*Santanichthys diasii*. A, detailed view of the caudal-fin skeletal structure. B, interpretative drawing of the caudal-fin skeletal anatomy. Exemplar AMNH 13981-X, illustrated by Filleul & Maisey (30, fig. 9) as AMNH 20068. **Abbreviations**: 1–7, hypurals 1–7; HsPu2, haemal spine of preural centrum 2; NPu1, neural arch and spine of preural centrum 1; NsPu2, neural spine of preural centrum 2; Phy, parhypural; Pu1, Pu2, preural centrum 1, 2; U1, U2, ural centrum 1, 2; Un1, Un2, Un3 uroneural 1, 2, 3. Scale bar represents 2 mm.

**Fig SX26.**
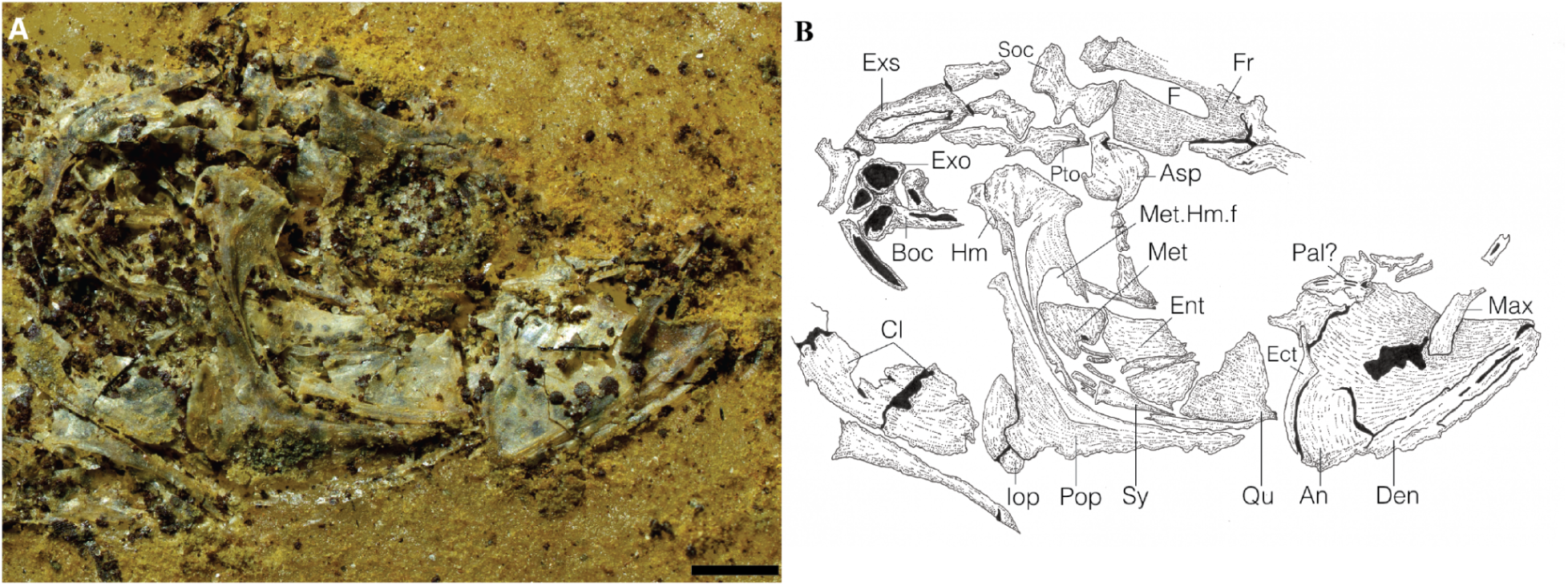
†*Santanichthys diasii*. A, detailed view of the head. B, interpretative drawing of the head skeletal anatomy. Exemplar AMNH 13981-P, illustrated by Filleul & Maisey (30, fig. 3) as AMNH 20062. **Abbreviations**: An, angulo-articular; Asp, autosphenotic; Boc, basioccipital; Cl, cleithrum; Den, dentary; Ect, ectopterygoid; Ent, entopterygoid; Exo, exoccipital; Exs, extrascapular; F, fontanelle; Fr, frontal; Iop, interopercle; Hm, hyomandibula; Max, maxilla; Met, metapterygoid; Met.Hm.f, metaptergoid-hyomandibula fenestra; Pal, palatine; Pop, preopercle; Pto, pterotic; Qu, quadrate; Sy, sympletic. Scale bar represents 2 mm.

**Fig SX27.**
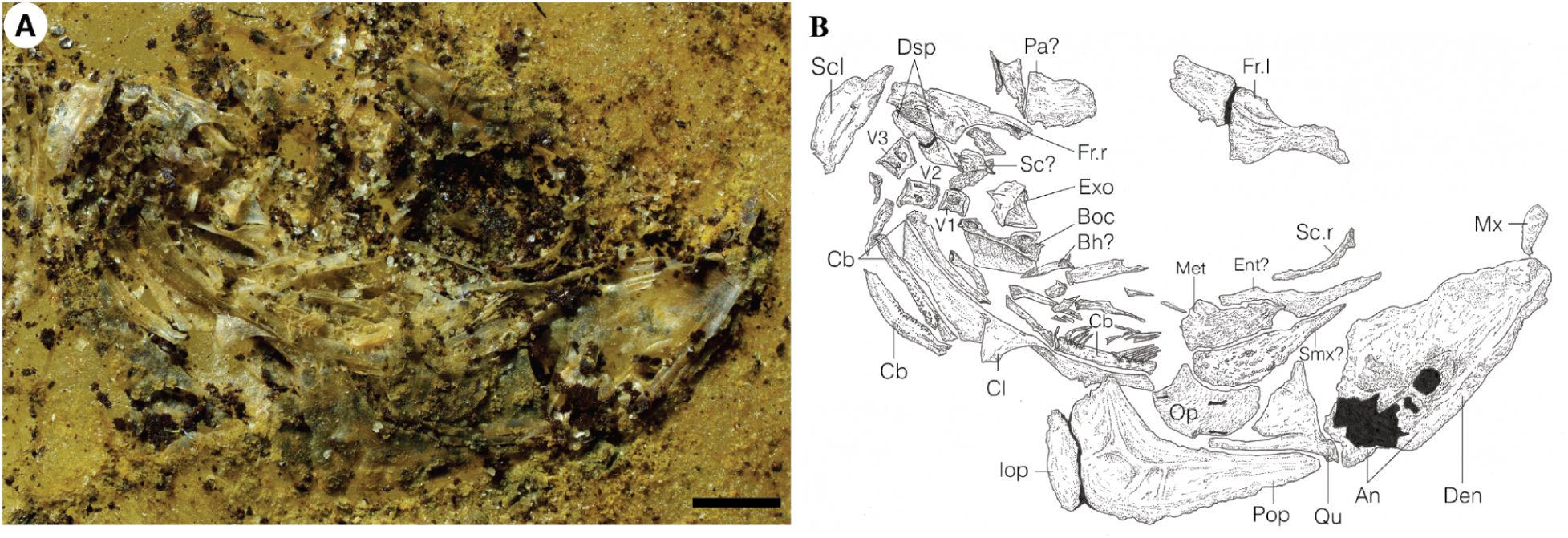
†*Santanichthys diasii*. A, detailed view of the head. B, interpretative drawing of the head skeletal anatomy. Exemplar AMNH 13981-R, illustrated by Filleul & Maisey (30, fig. 6) as AMNH 20064. **Abbreviations**: An, angulo-articular; Bh, basihyal; Boc, basioccipital; Cb, ceratobranchial; Cl, cleithrum; Den, dentary; Dsp, dermosphenotic; Ent, entopterygoid; Exo, exoccipital; Fr.l, left frontal; Fr.r, right frontal; Iop, interopercle; Max, maxilla; Met, metapterygoid; Op, opercle; Pa, parietal; Pop, preopercle; Qu, quadrate; Sca, scaphium; Sc.r; sclerotic ring; V1, V2, V3, vertebra one, two, three. Scale bar represents 2 mm.

**Table TX1. Revised morphological matrix of sampled extant and fossil otophysan fishes**. Unclear or undiagnosable traits are marked with “?” and inapplicable traits are marked with “-”. See supplementary text for detailed analysis of recoding, and (Supplementary Figure SX01) for graphical depiction of changes in coding.

**Table TX2. Genomic data used in the present study, with habitat coding.** Composite taxa are indicated with a yellow highlight. All species with UCE data are taken from the alignment of (1). Species with exonic data originate from published data (NCBI or SRA accession number provided, n=57) or from newly generated target capture data (indicated with bold text and asterisk; tissue or specimen voucher provided, n=22). Museum abbreviation codes follow (91)

**Table TX3. Model fit results for BioGeoBEARS**. BAYAREALIKE+J was recovered as the best fitting biogeographic model in all 18 rounds of model fitting. Despite being the model selected by Liu et al., DEC was never recovered as the best fitting biogeographic model, and is ranked the worst-fitting model in 5 out of 6 trials using the topology recovered by Liu et al.

**Table TX5. Transition probabilities according to BioGeoBEARS, measured as joint probabilities of the node states for crown Otophysa, Characiphysae, and Cypriniphysae.** (**A**) A single freshwater invasion precedes the otophysan crown, and is retained in both Cypriniphysae and Characiphysae. (**B**) A single freshwater invasion precedes the otophysan crown, followed by a return to marine habitats in one daughter lineage. (**C**) A euryhaline ancestor remains euryhaline in both daughter lineages, which independently become freshwater endemics. (**D**) A euryhaline ancestor remains euryhaline in only one daughter lineage, and undergoes a habitat contraction in the other. (**E**) A euryhaline ancestor undergoes two parallel habitat contractions. (**F**) A euryhaline ancestor undergoes habitat segregation, producing one freshwater and one marine daughter lineage. (**G**) A marine ancestor undergoes two parallel freshwater invasions. All other transition schemes never returned a probability ≥0.0001.

**Table TX6. Characters supporting potential affinities of †*Santanichthys diasii*.** Columns CLU and OST correspond to observed character states in Clupeomorpha and Ostariophysi, respectively; column †*San* corresponds to character state recorded in †*Santanichthys diasii* following the present study. Final column states the extant-lineage affinity indicated by each character; **CLU** indicates that the †*Santanichthys* state is only recorded in clupeomorphs (n = 6), and **OST** indicates that the †*Santanichthys* state is only recorded in ostariophysians (n = 17). Characters coded as “**-**” are those in which the †*Santanichthys* state is ambiguous, inapplicable, or observed in both living clupeomorphs and ostariophysians.

**Table TX7. Maximum Parsimony results.**

Number of best trees, number of steps, consistency index and retention index measured for every weighting scheme under maximum parsimony. For *k* = 1 of the unconstrained run, two values are given as there were trees of different lengths that had the best score using that concavity constant.

**Table TX8. Maximum Likelihood results.**

Log likelihoods and AU score for unconstrained and constrained ML trees inferred using both our revised matrix and the original Liu et al. matrix. Using our revised matrix, our topology is strongly favored and Liu et al. is rejected, while using the Liu matrix, their topology is strongly favored, ours fails to be rejected.

**Table TX9. Bayesian Inference results**

Marginal likelihoods and Bayes factor comparisons between trees inferred using our revised matrix under Bayesian inference. Three runs were conducted, one unconstrained, one constrained to match the topology of Liu et al., and one to match our unconstrained topology to reduce the tree space of the search.

