## Supplementary figures and images for "A single freshwater invasion gave rise to the world’s dominant freshwater fish group"

### Figure SX01

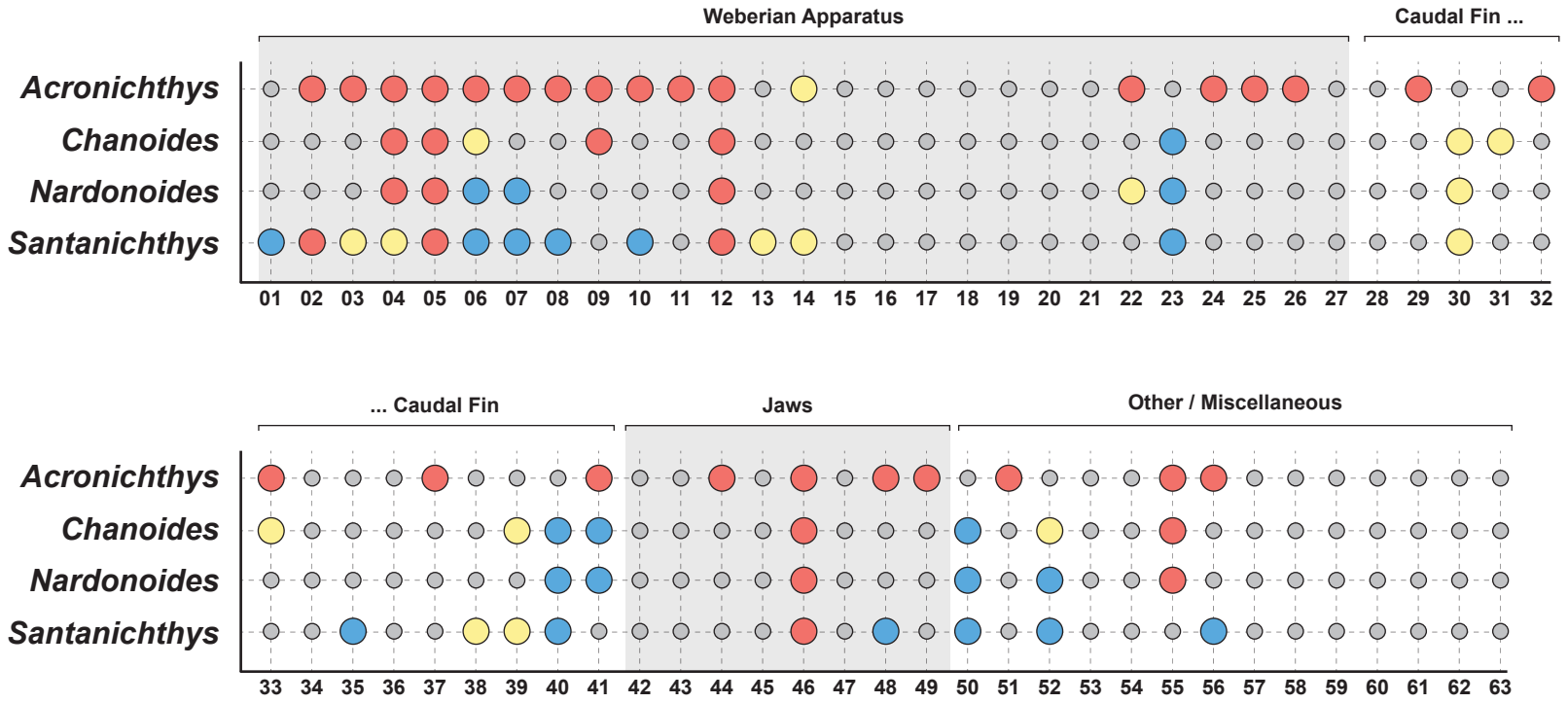

### Figure SX02

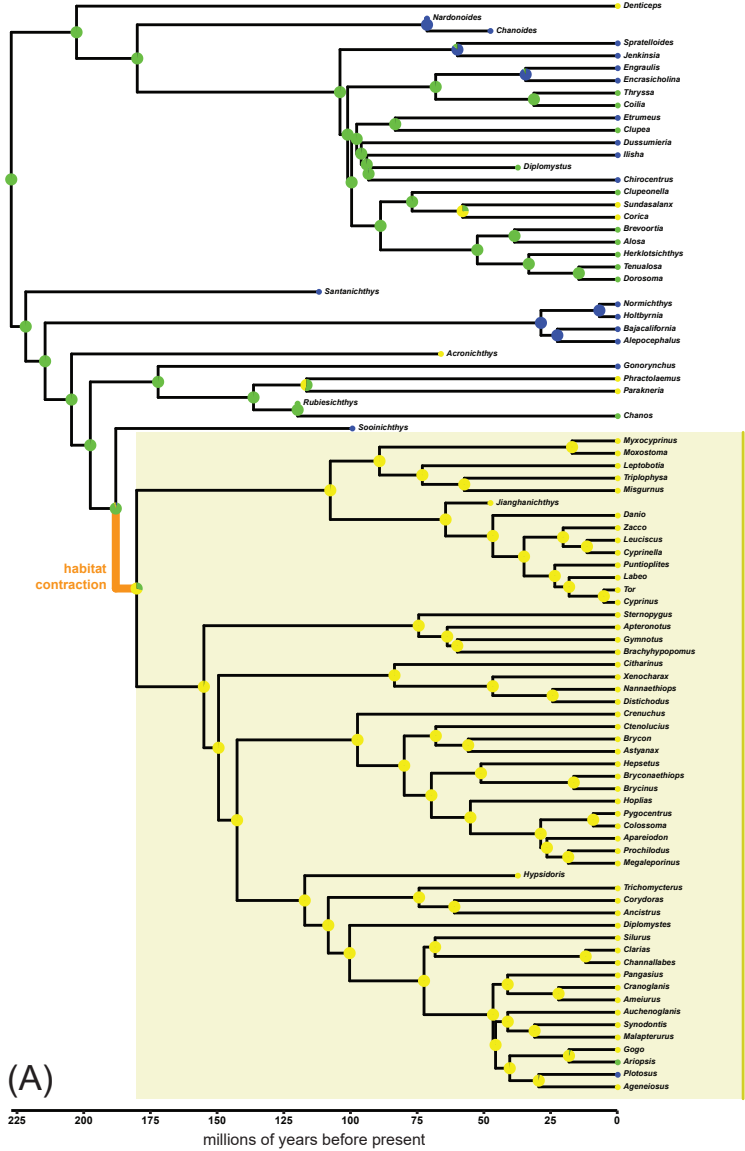

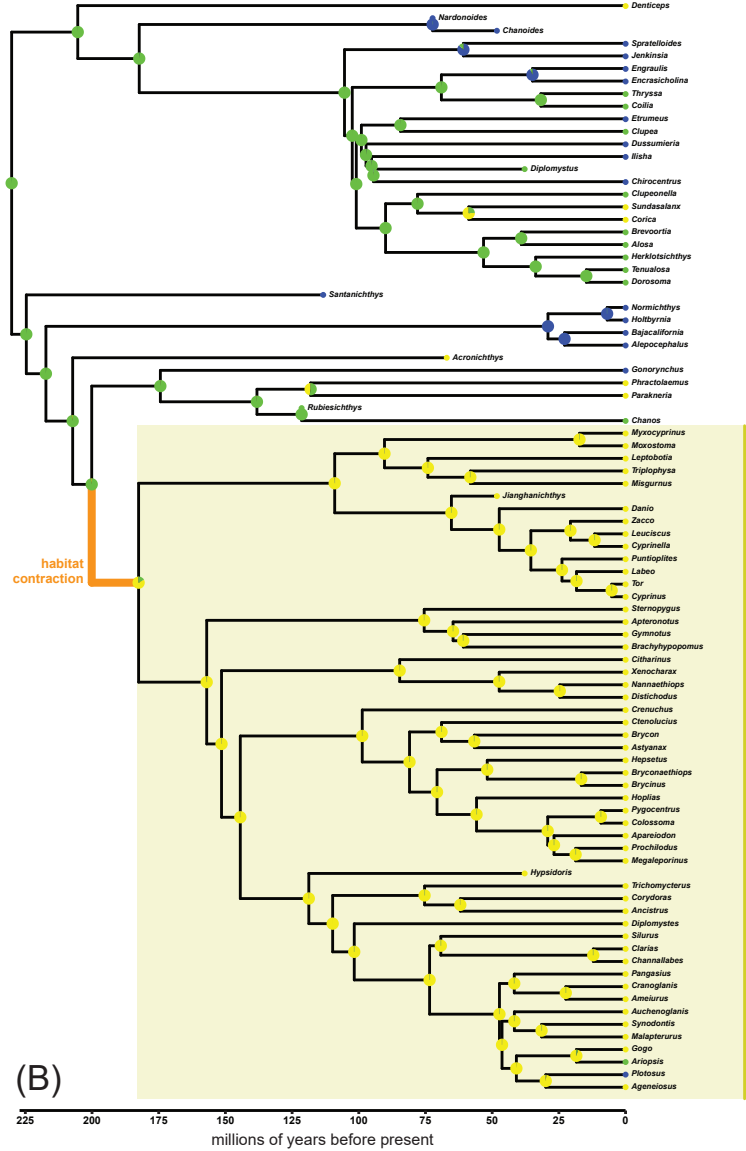

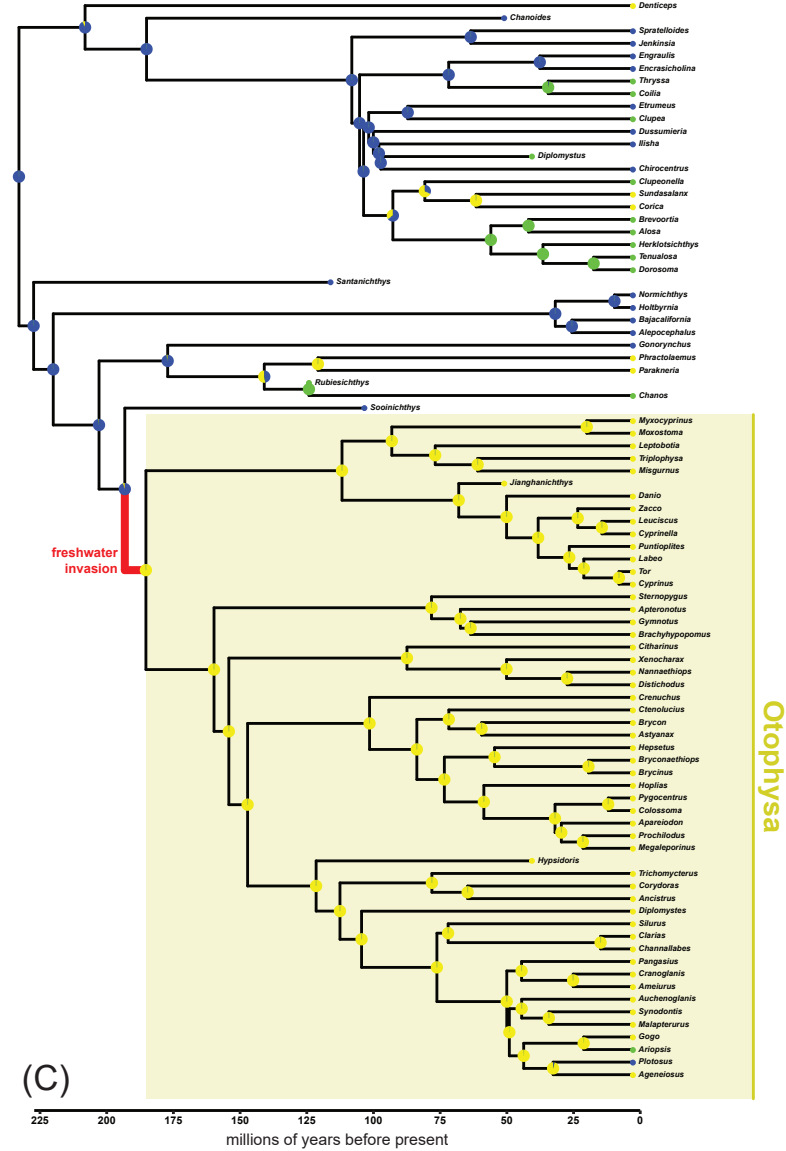

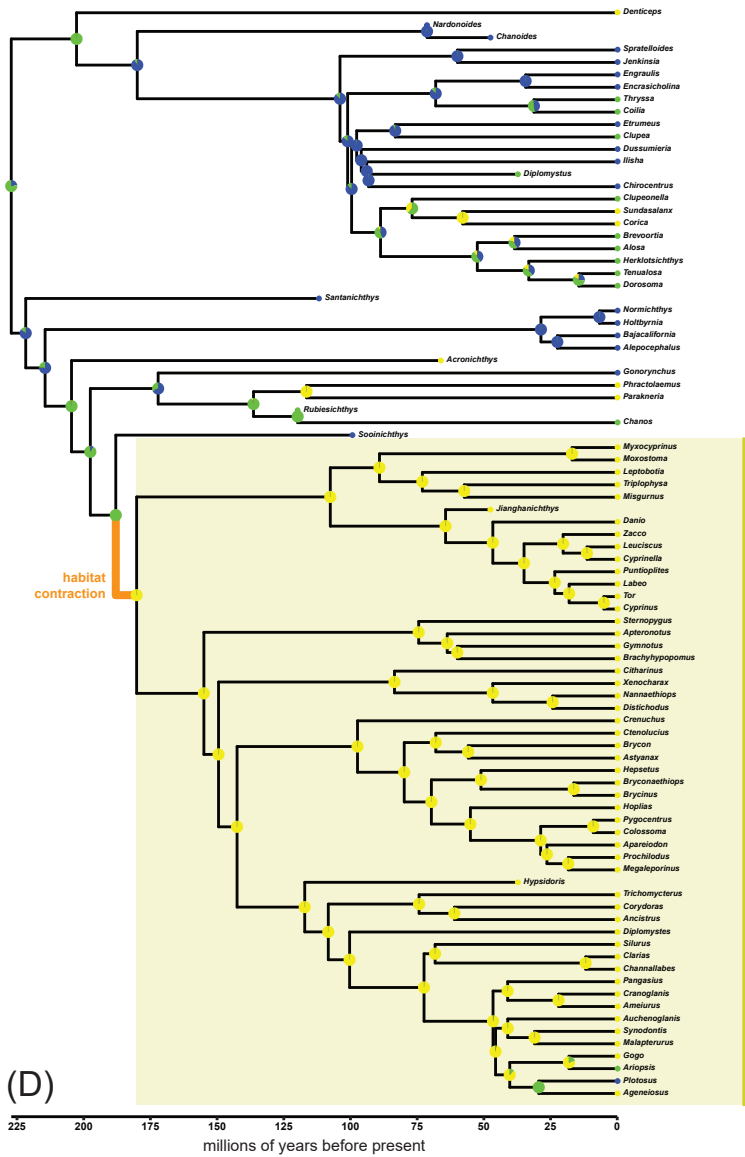

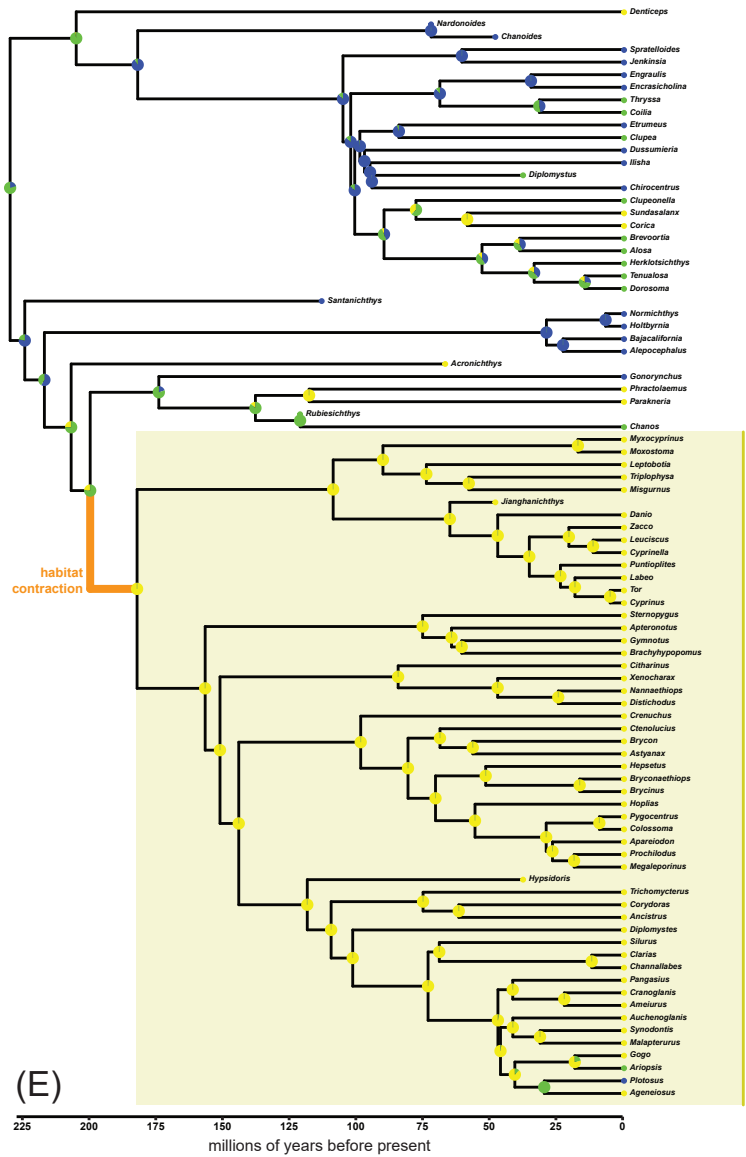

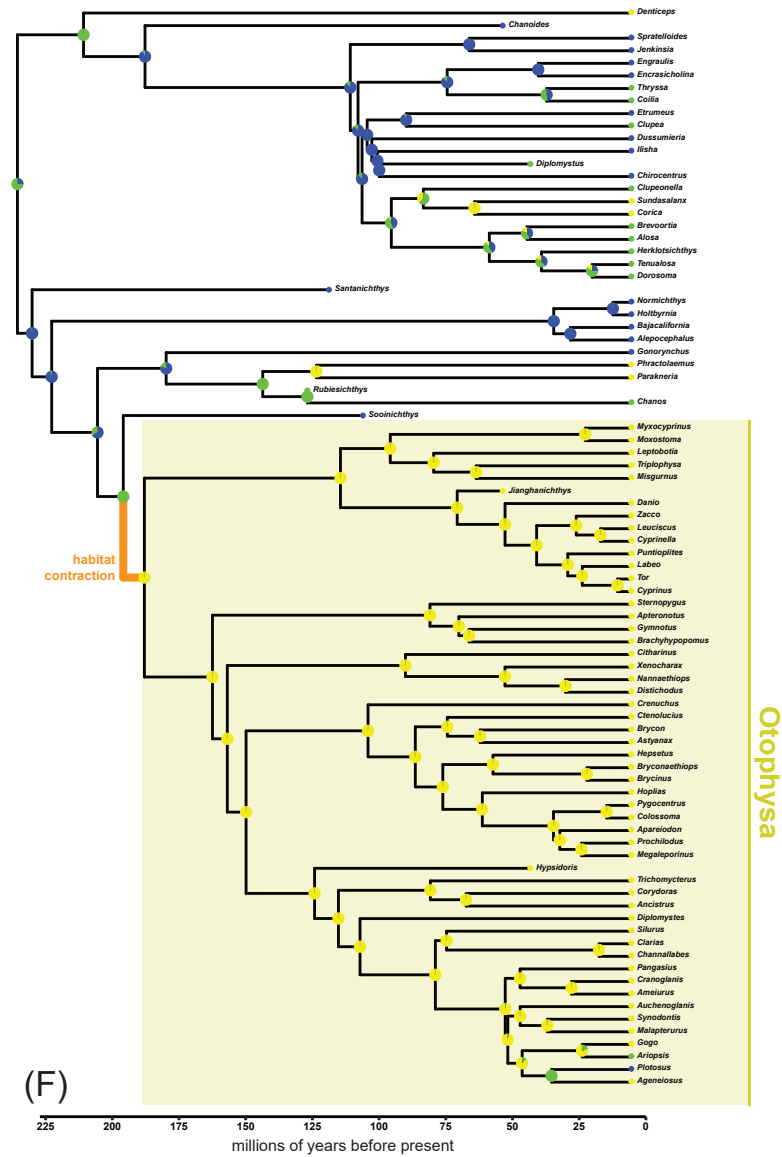

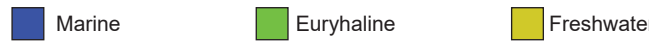

### Figure SX03

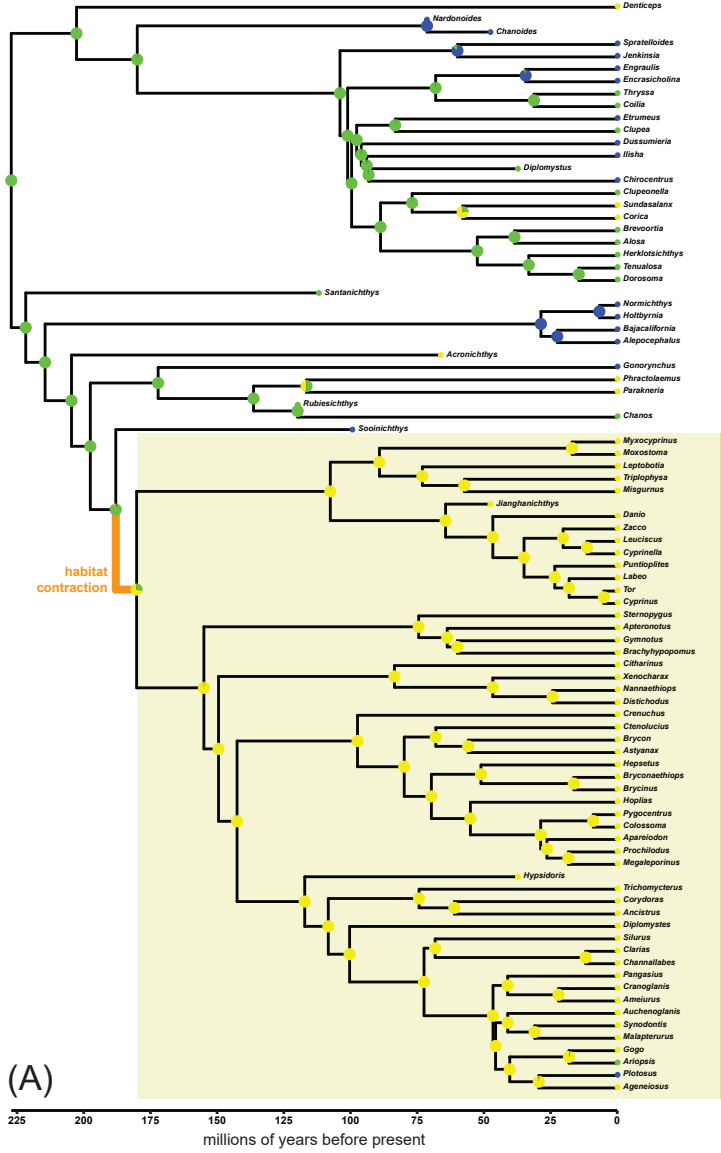

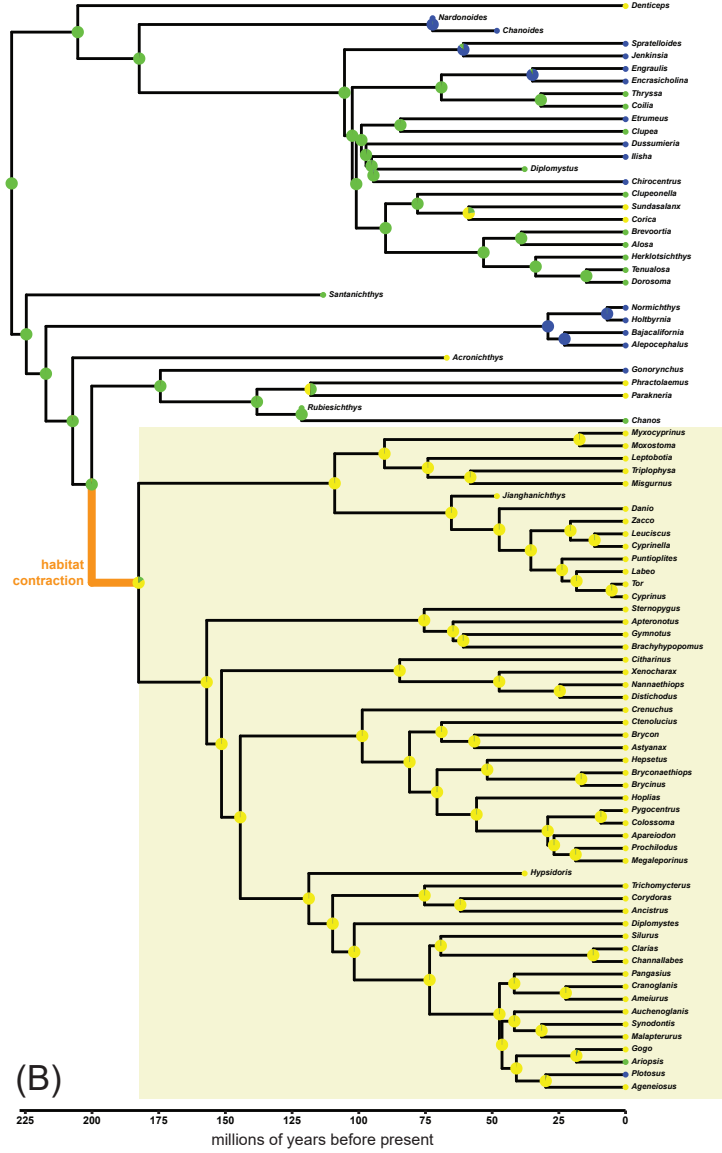

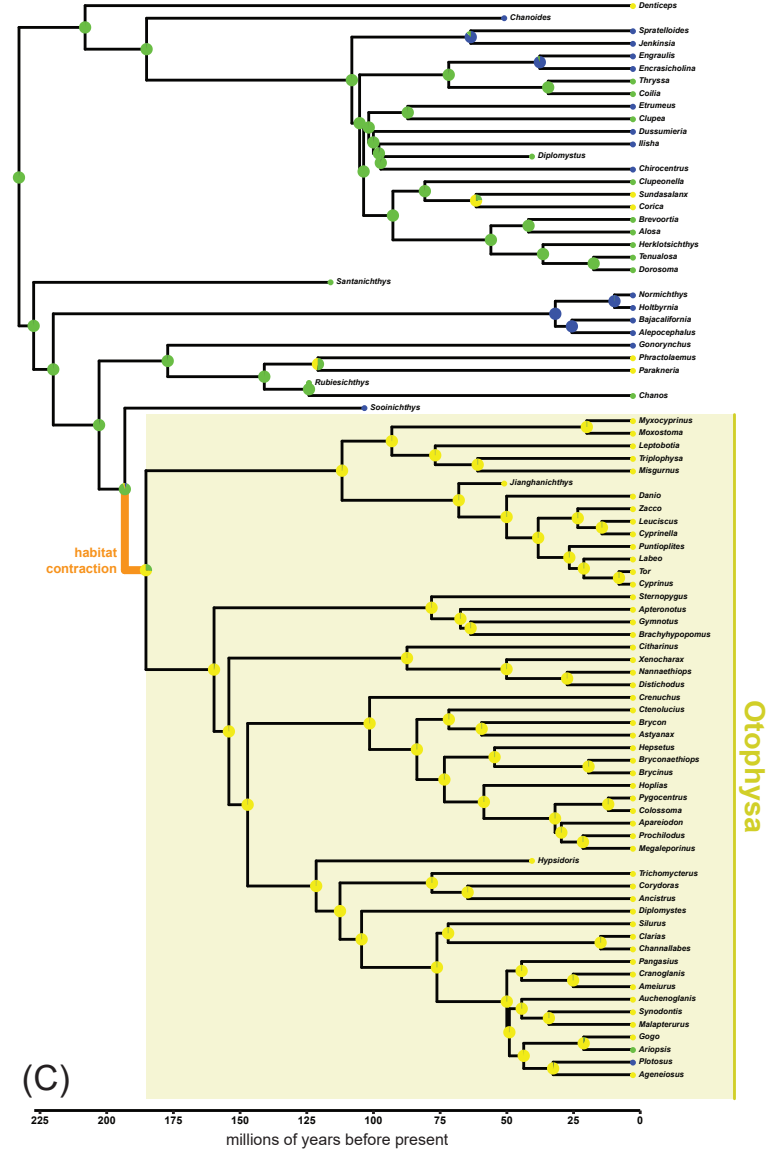

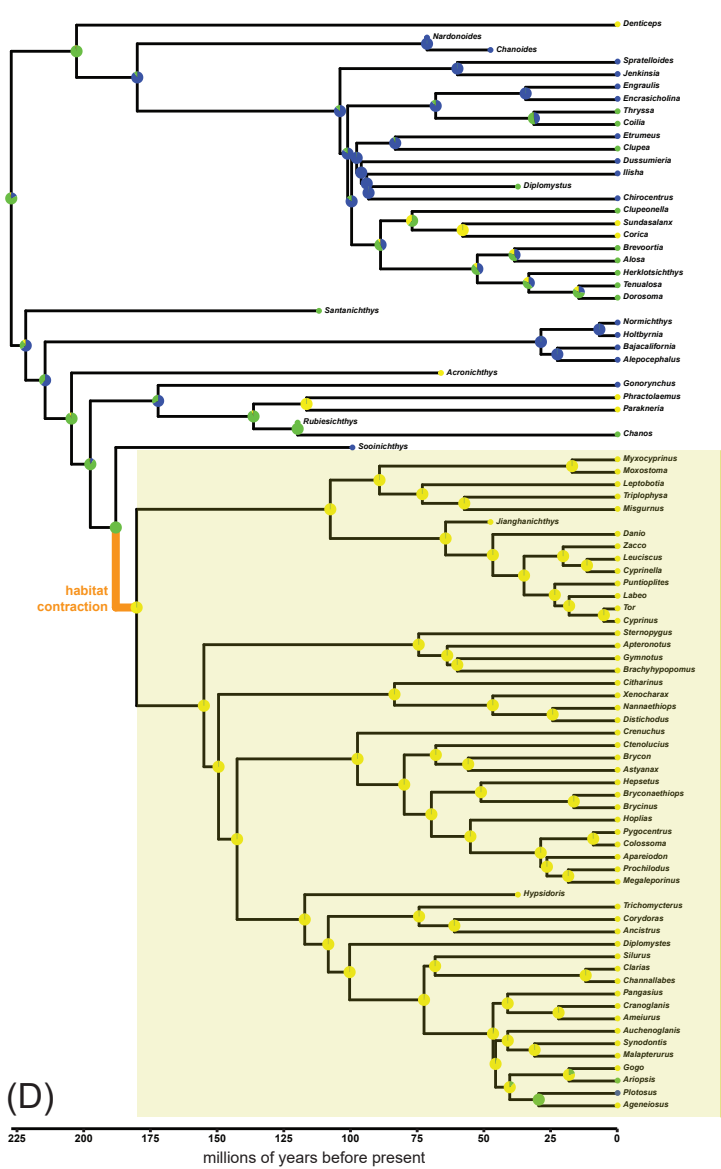

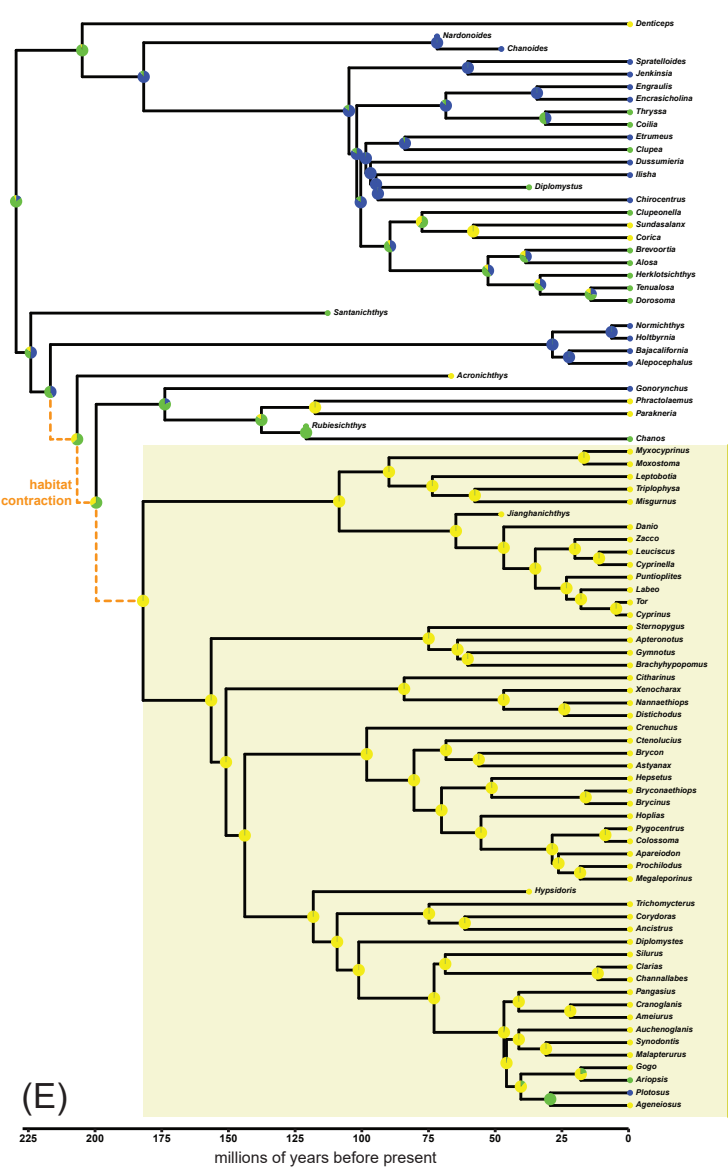

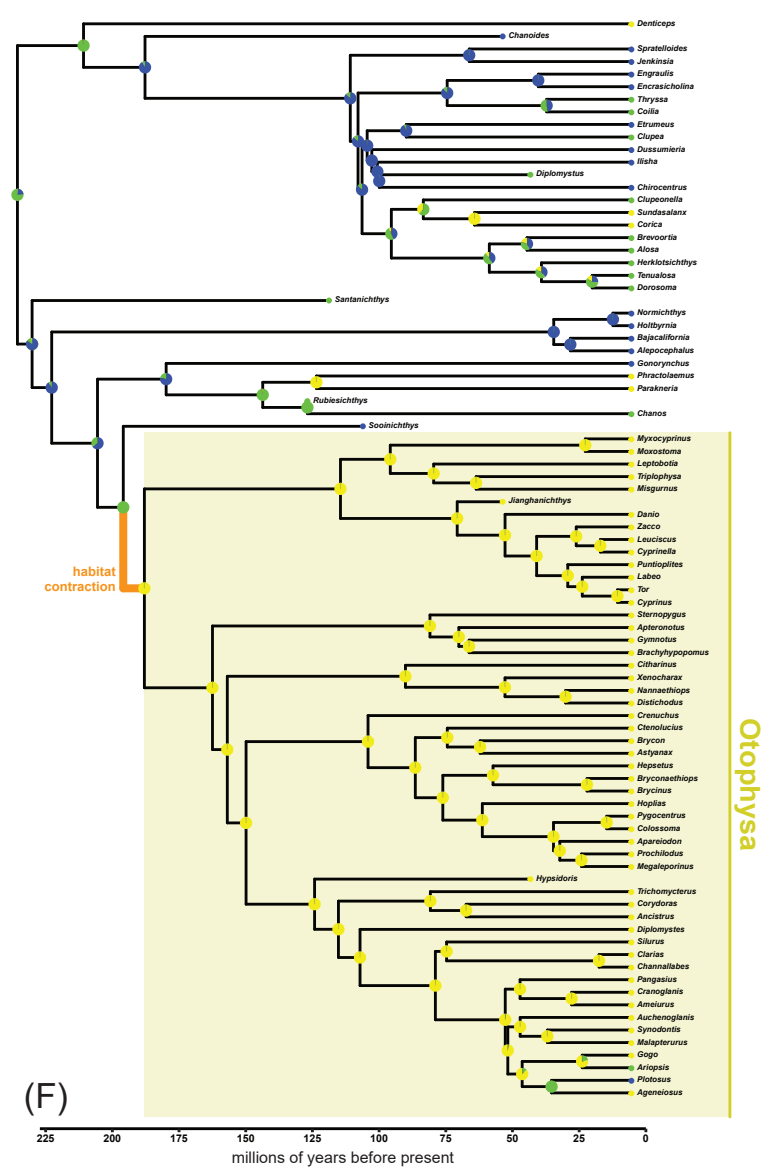

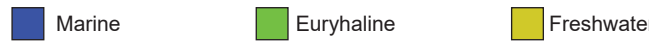

### Figure SX04

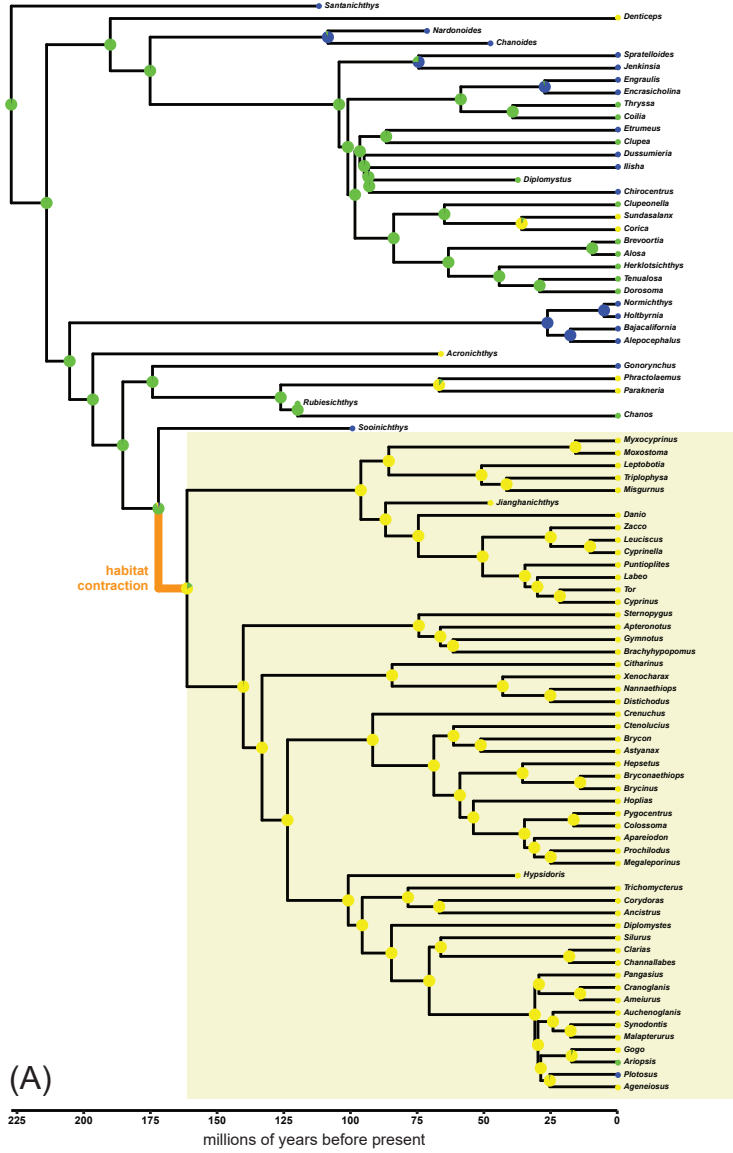

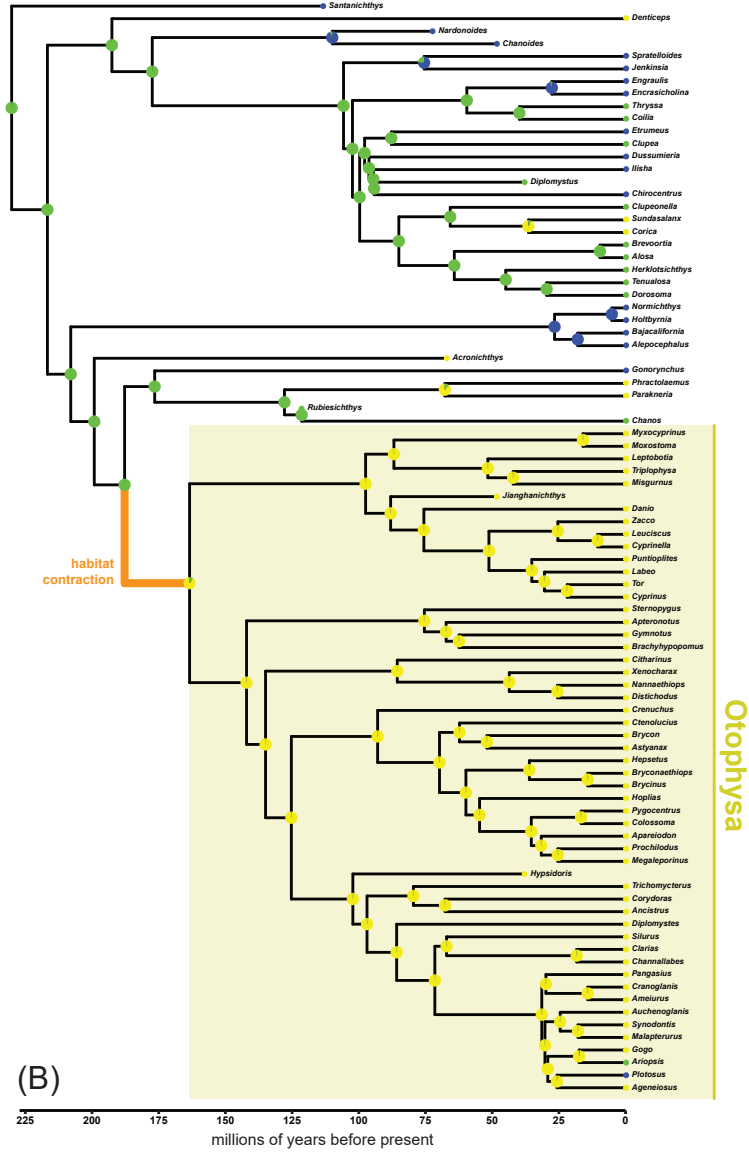

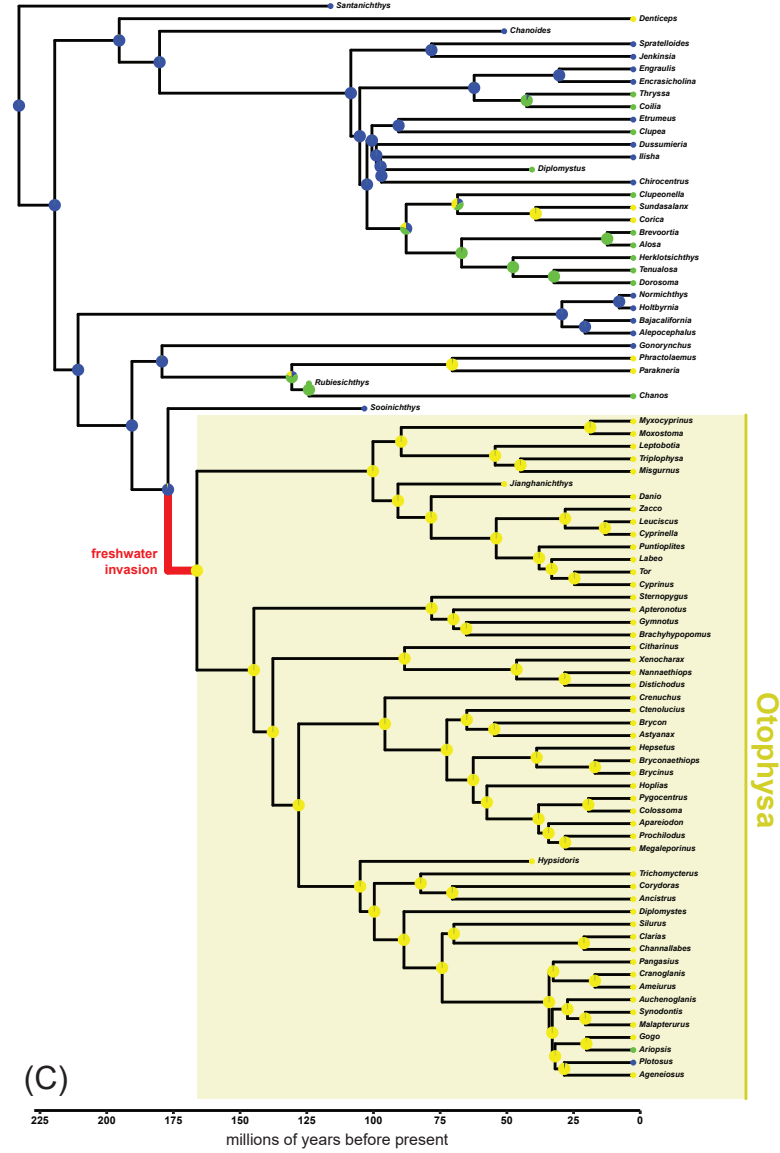

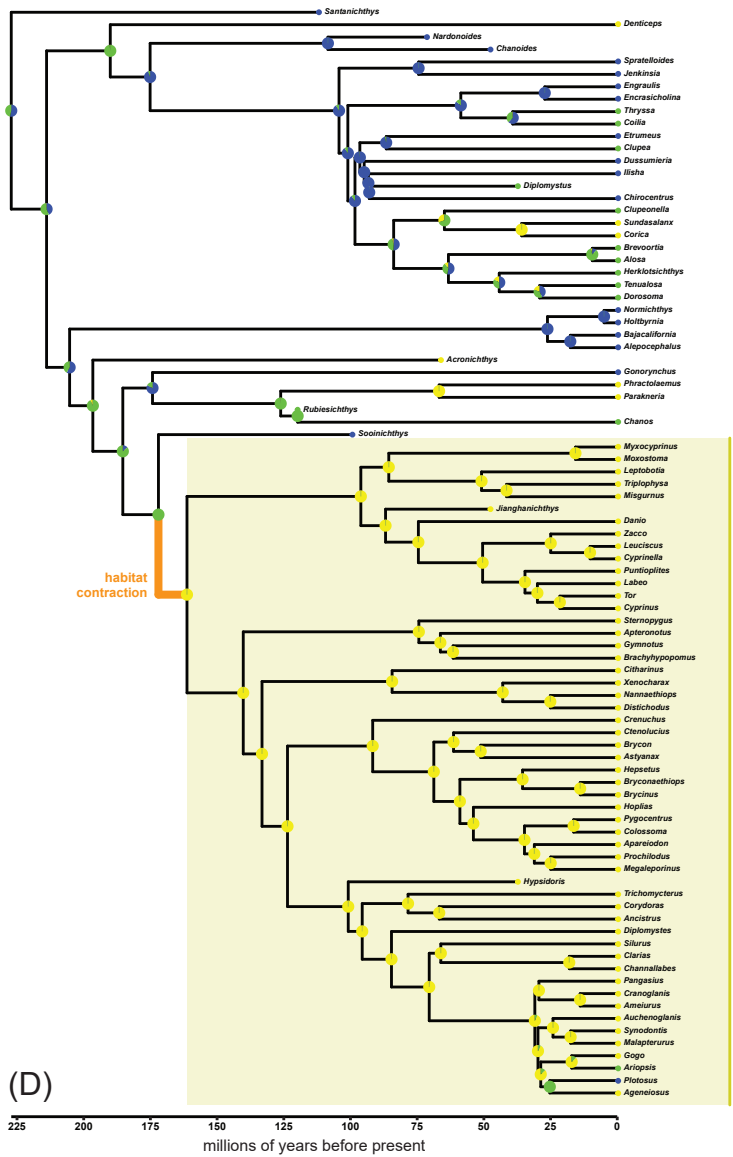

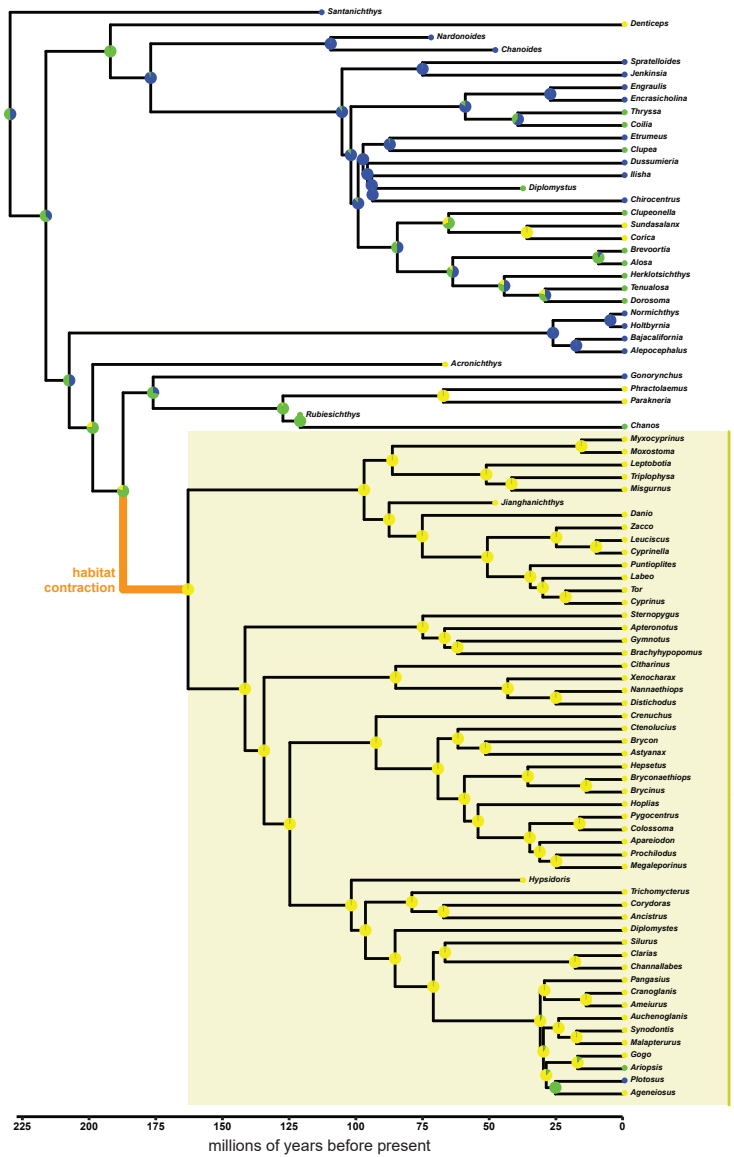

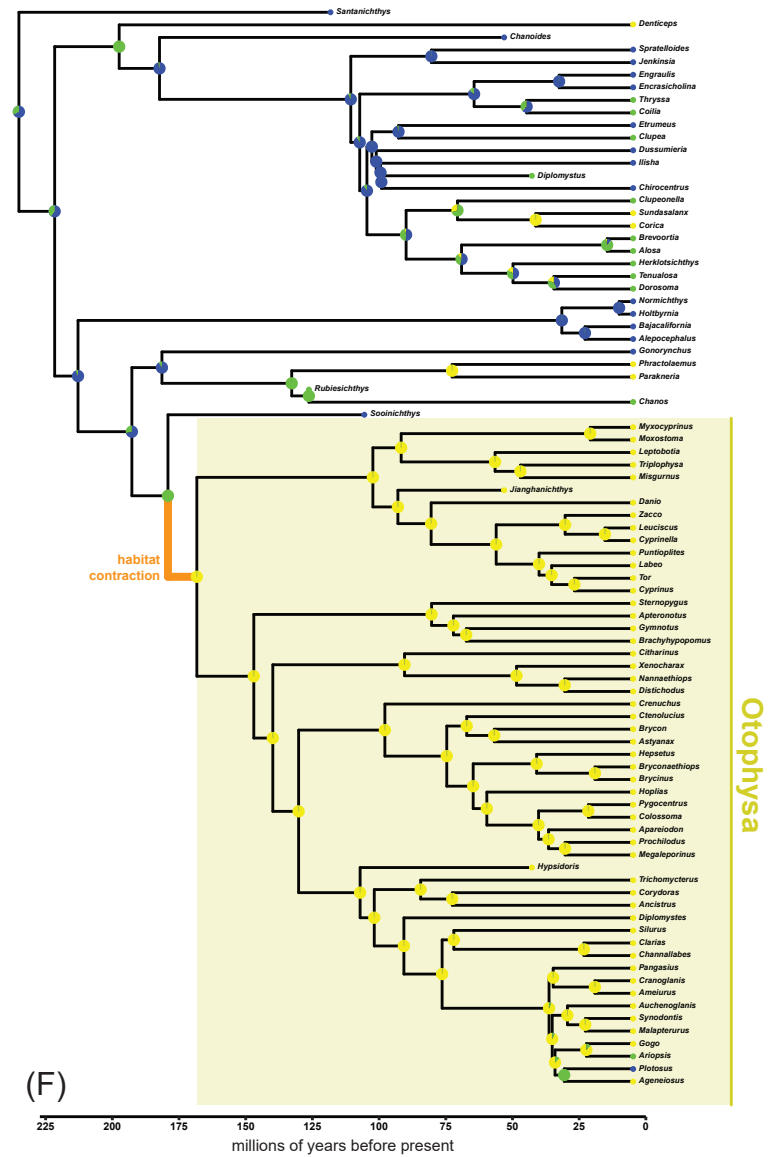

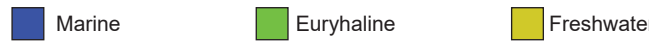

### Figure SX05

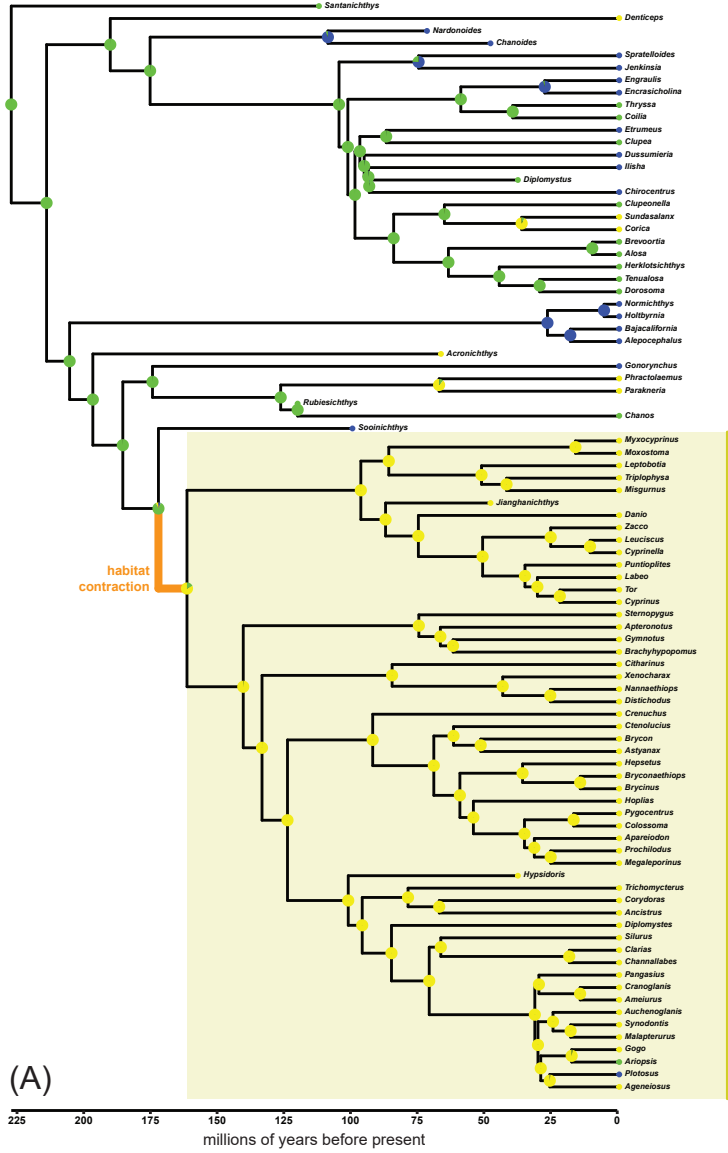

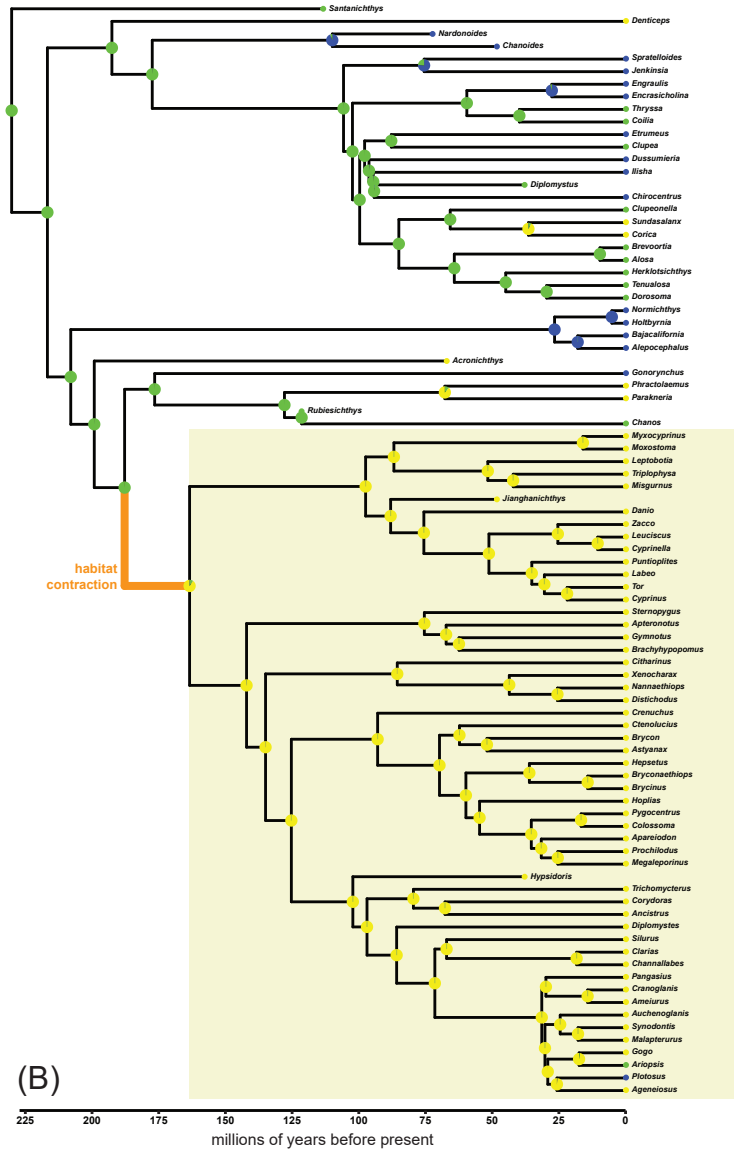

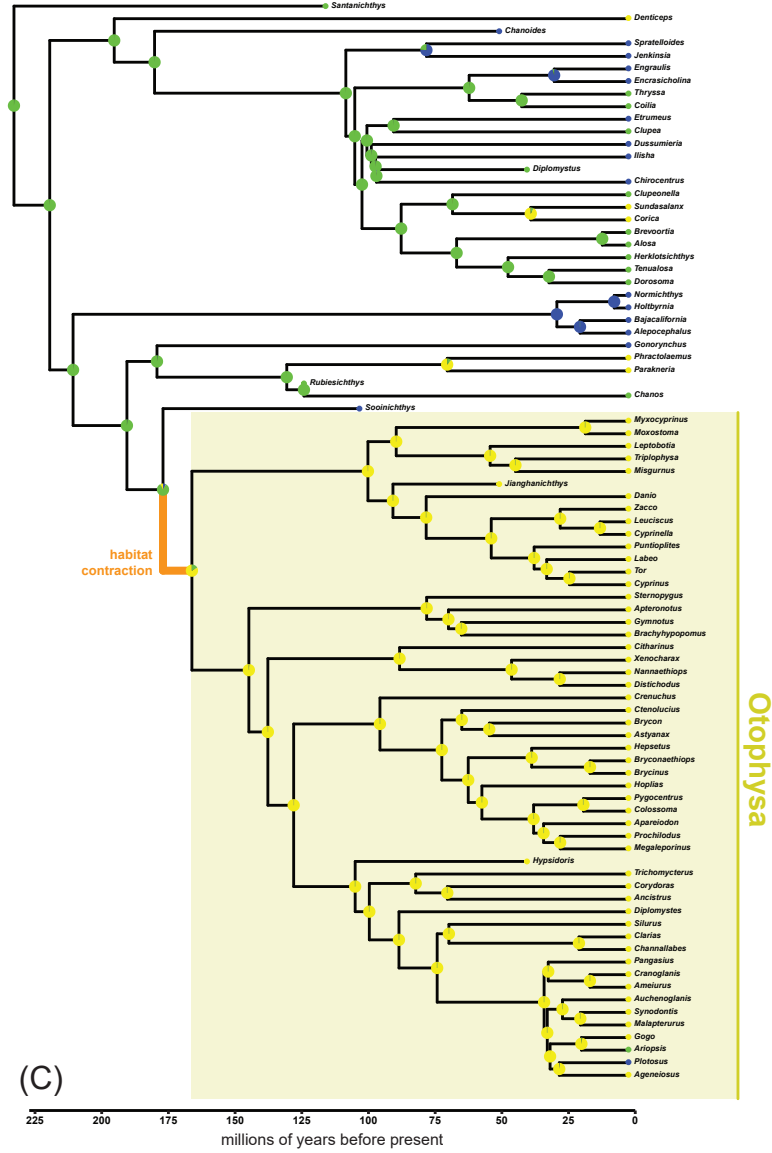

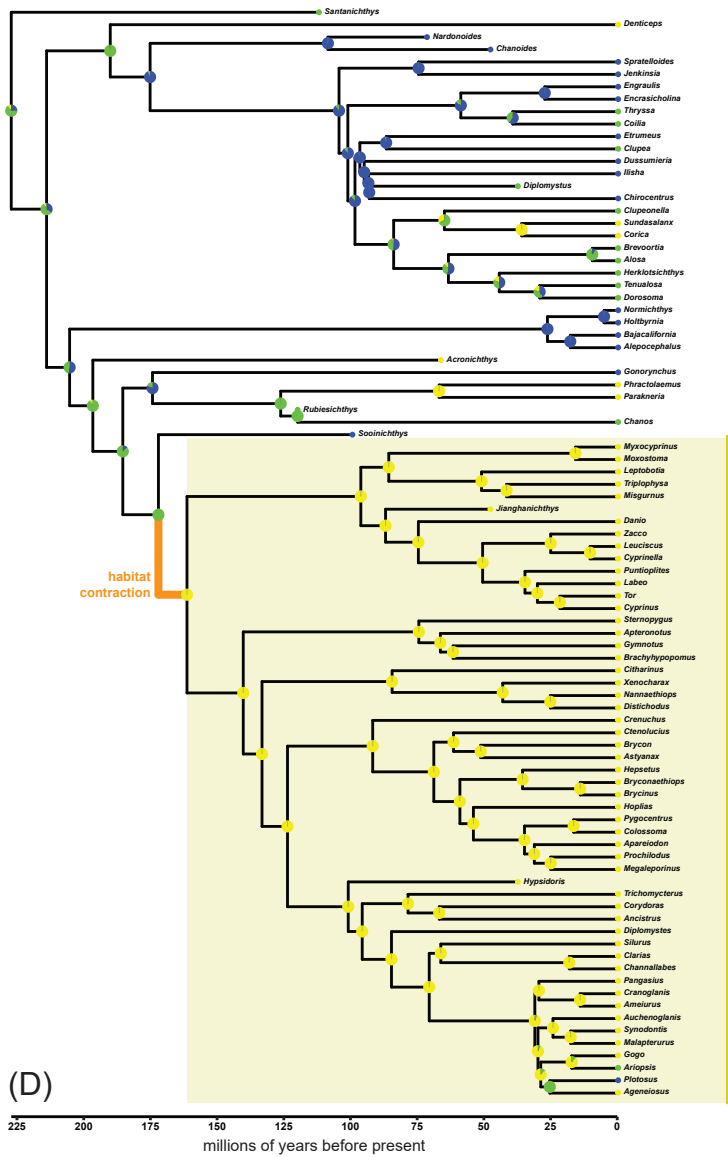

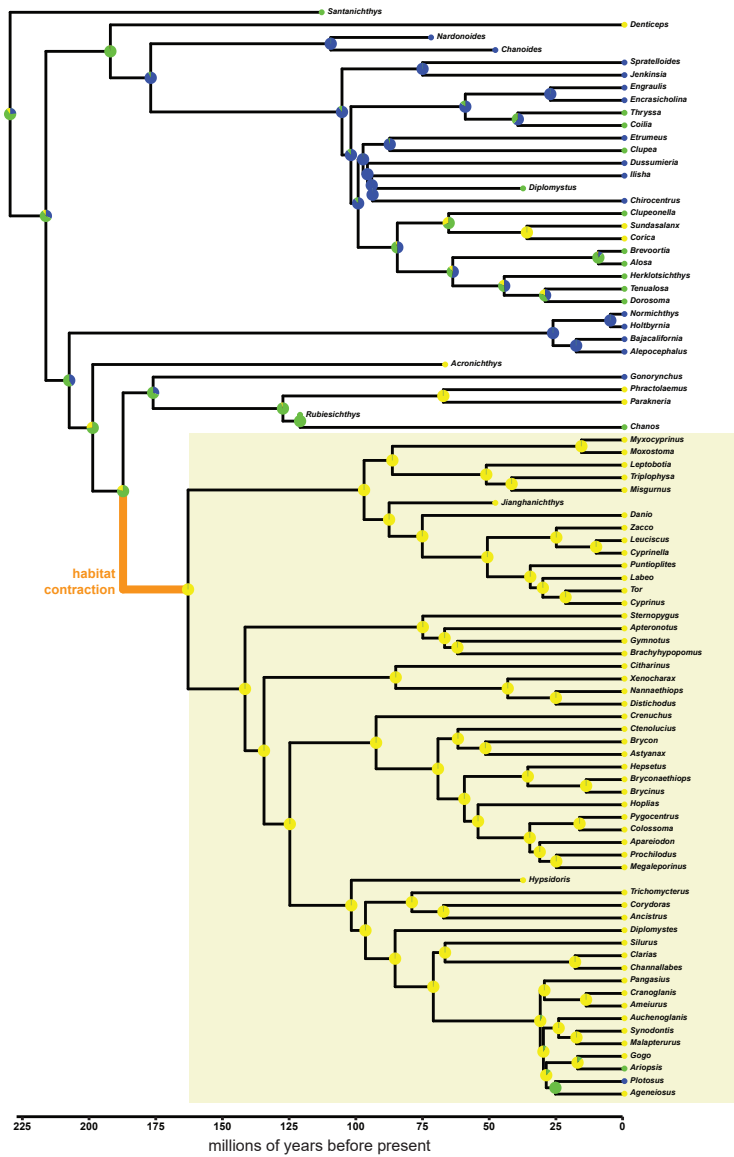

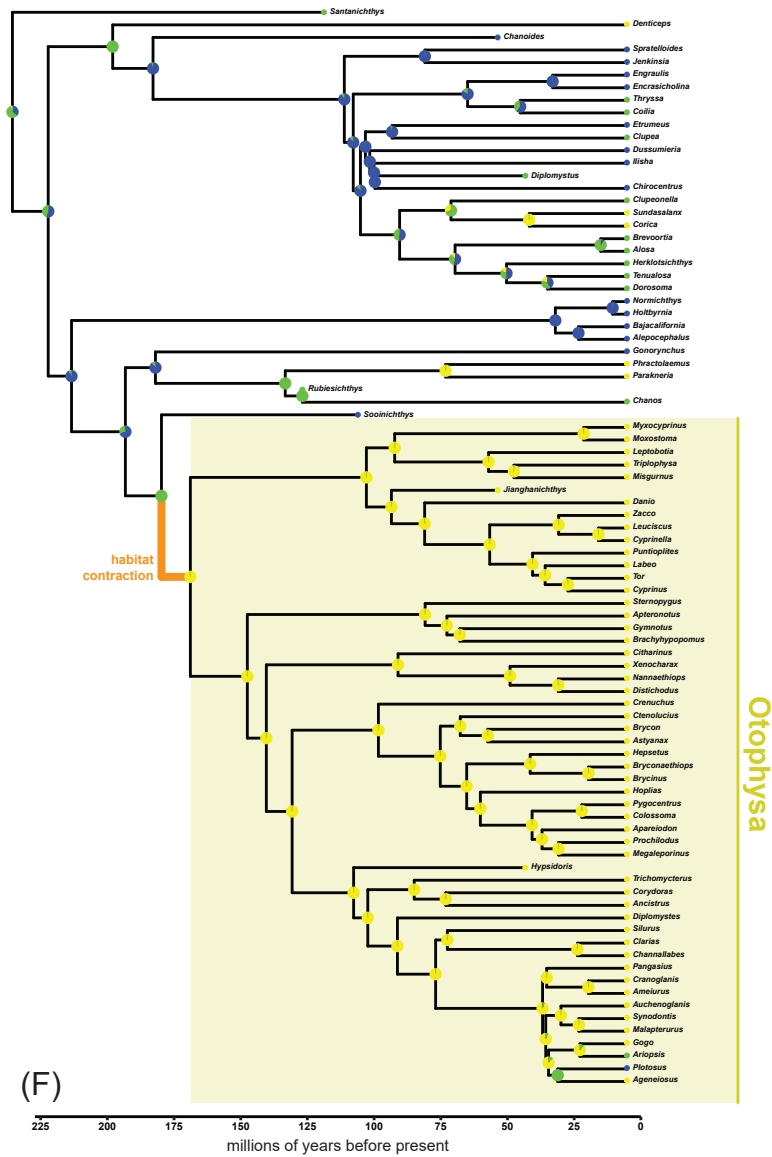

Marine Euryhaline Freshwater

### Figure SX06

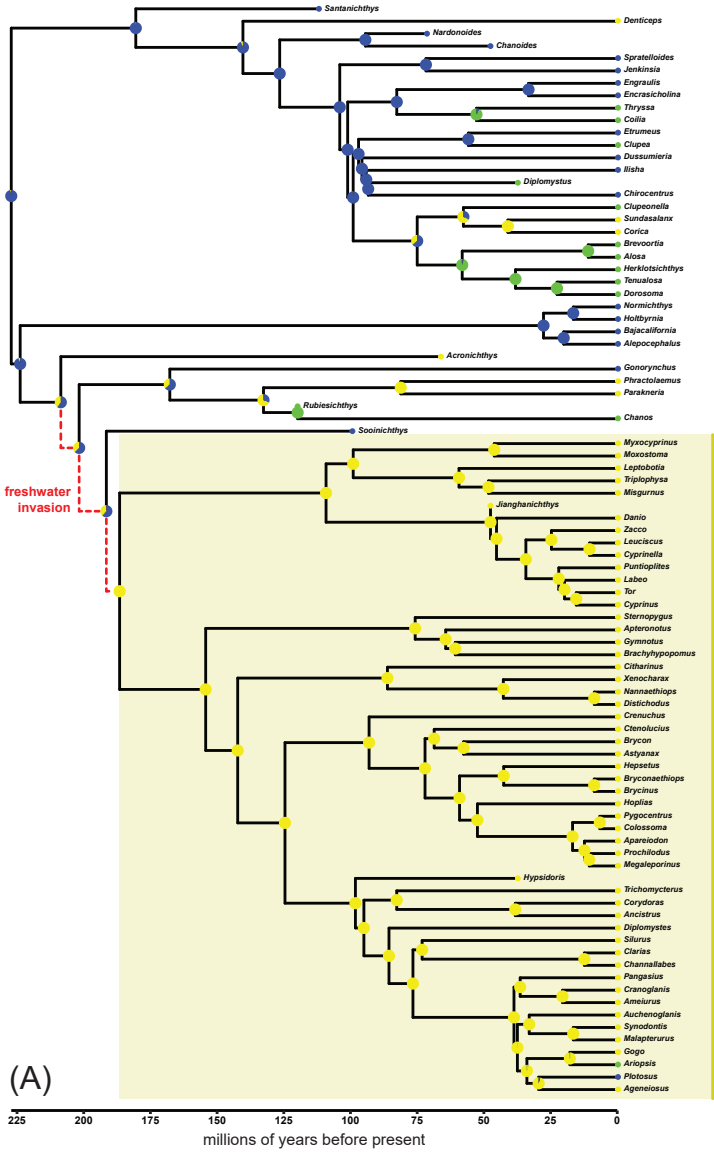

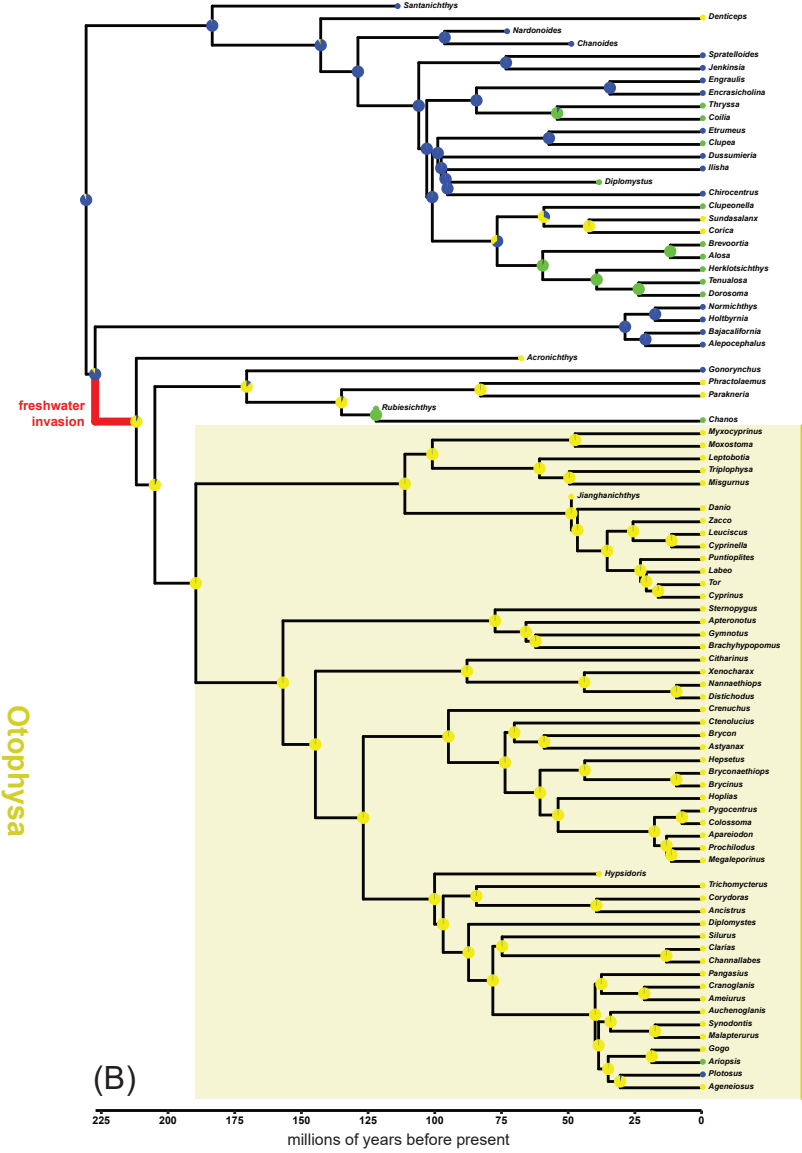

Marine Euryhaline Freshwater

### Figure SX08

Marine Euryhaline Freshwater

### Figure SX10

Marine Euryhaline Freshwater

### Figure SX11

Marine Euryhaline Freshwater

### Figure SX13

Marine Euryhaline Freshwater

### Figure SX14

Marine Euryhaline Freshwater

### Figure SX15

Marine Euryhaline Freshwater

### Figure SX16

Marine Euryhaline Freshwater

### Figure SX17

Marine Euryhaline Freshwater

### Figure SX18

Marine Euryhaline Freshwater

### Figure SX19

Marine Euryhaline Freshwater

### Figure SX20

Marine Euryhaline Freshwater

### Figure SX21

(A)

(B)

(C)

(D)

(E)

(F)

Marine Euryhaline Freshwater

### Figure SX22

Marine Euryhaline Freshwater

### Figure SX23

Marine Euryhaline Freshwater
